# Mobile Integrons Encode Phage Defense Systems

**DOI:** 10.1101/2024.07.02.601719

**Authors:** Nicolas Kieffer, Alberto Hipólito, Laura Ortiz-Miravalles, Paula Blanco, Thomas Delobelle, Patricia Vizuete, Francisco Manuel Ojeda, Thomas Jové, Dukas Jurenas, Meritxell García-Quintanilla, André Carvalho, Pilar Domingo-Calap, José Antonio Escudero

## Abstract

Integrons are bacterial genetic elements that capture, stockpile and modulate the expression of genes encoded in integron cassettes. Mobile Integrons (MI) are borne on plasmids, acting as a vehicle for hundreds of antimicrobial resistance genes among key pathogens. These elements also carry <u>g</u>ene <u>c</u>assettes of <u>u</u>nknown function (*gcu*s) whose role and adaptive value remains unexplored. Here we show that *gcu*s encode phage resistance systems, many of which are novel. <u>B</u>acteriophage <u>r</u>esistance <u>i</u>ntegron <u>c</u>assettes (BRiCs) can be combined and mixed with resistance cassettes to produce multiphage or drug/phage-resistance. The fitness costs of BRiCs are variable, dependent on the genetic context, and can be modulated by changing the order of cassettes in the array. Hence, MIs act as highly mobile, low-cost defense islands.

Summary Figure
Novel phage defense systems identified in Mobile Integrons.
We confronted genes of unknown function from mobile integrons against a panel of phage. We characterized 13 Bacteriophage Resistance integron Cassettes (BRiCs) and confirmed their function in *Klebsiella pneumoniae* and *Pseudomonas aeruginosa*. Combined with other cassettes, BRiCs produce multi-phage/antibiotic resistance. Additionally, their cost can be reduced in an array.

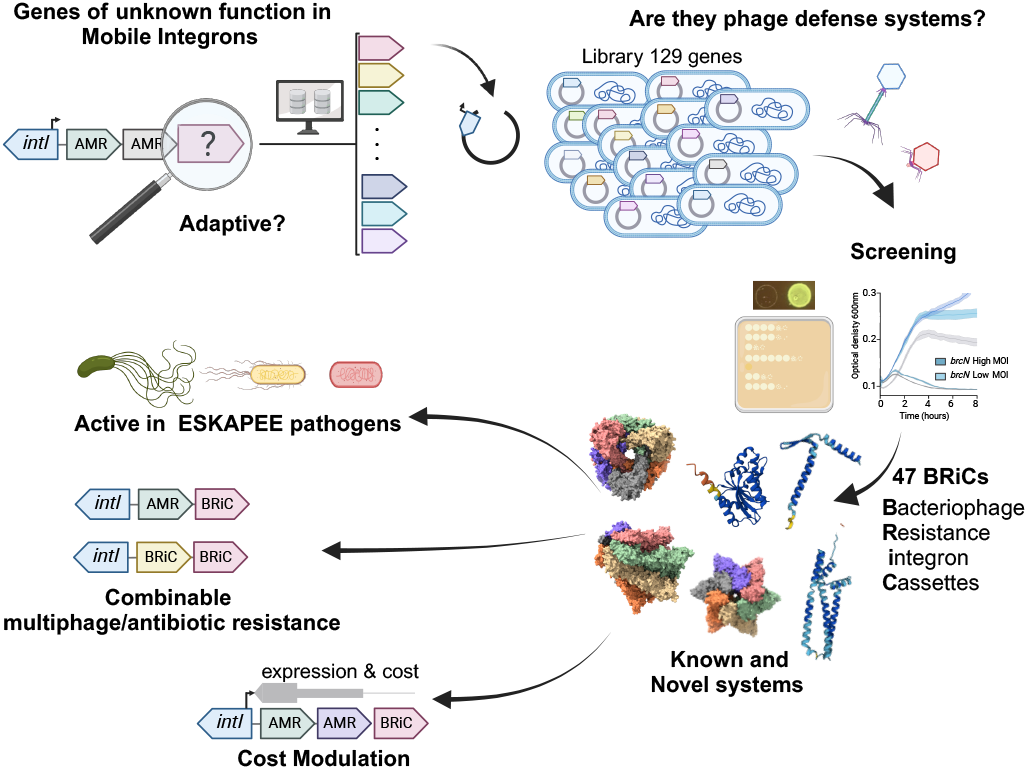

## INTRODUCTION

Antimicrobial resistance (AMR) is a major public health concern worldwide, and the emergence of multi-drug (MDR) resistant bacteria is making increasingly difficult to treat infections with antibiotics (*1*). Phage therapy - the use of viruses that infect and kill bacteriais currently building momentum as an alternative to antibiotics (*2*–*4*). However, its efficacy can be limited too by the emergence of resistance (*5, 6*). In recent years, a plethora of new phage defense systems (PDSs) have been discovered (*7*), often co-localizing in defense islands (*8*–*10*). Some PDSs are encoded in mobile genetic elements (MGEs) such as integrative conjugative elements, transposons, or prophages (*11*). While their spread is a threat to phage therapy, PDSs can entail a fitness cost to their host limiting their dissemination (*12*).

Integrons are genetic elements that play an important role in bacterial adaptation to changing environmental conditions (*13*–*15*). They capture and accumulate new genes embedded in integron cassettes (ICs), acting as genetic memories of adaptive functions (*16*). Integrons typically consist of a conserved platform encoding the integrase gene and the recombination site in the integron (*attI*) (*17*); and a variable region containing the cassettes. Cassettes are incorporated into the *attI* site through site-specific recombination reactions mediated by the integrase and are expressed from the dedicated Pc promoter encoded in the platform (*18*). The integrase can also reorder cassettes in the array to modulate their expression, modifying their distance to the Pc and the polar effects they are subjected to (*19*–*21*). The expression of the integrase is controlled by the SOS response, so that integrons provide to their hosts adaptation on demand (*22, 23*). MIs are a subset of integrons associated with plasmids and transposons, that facilitate their transfer between bacterial cells (*24*–*26*). They are currently commonplace among key Gram-negative pathogens (*27*) carrying almost 200 resistance genes against most antibiotic families (*28*–*30*). Although generally devoted to AMR, MIs also carry <u>g</u>ene <u>c</u>assettes of <u>u</u>nknown function (*gcu*s), whose importance has commonly been overlooked. The working model of integrons suggests that cassettes must be adaptive at the time of integration (*22*). Given the importance of phage predation in the lifestyle of bacteria, we sought to explore if *gcus* encode phage defense systems.

We have selected 129 non-redundant *gcus* from the INTEGRALL database (*31*), and cloned them as cassettes in a mobile integron (*30*). DefenseFinder (*32*) and PADLOC (*33*) predicted potential defense systems in 4 *gcus* in this collection. Screening the library against a panel of phage, we found 43 novel defense systems. We have characterized 13 systems further and confirmed that they are encoded in functional integron cassettes, which we have named <u>B</u>acteriophage <u>R</u>esistance integron <u>C</u>assettes (BRiCs). They are hence mobile and exchangeable between integron platforms that circulate among many species. We demonstrate that they also confer protection in *K. pneumoniae* and *P. aeruginosa*, major pathogens of the ESKAPEE group. As part of an integron, defense systems can be stockpiled in an array where their position allows to modulate their function and strongly diminish their cost. We also show that phenotypes encoded in BRiCs are additive, conferring multi-phage or phage/drug resistance if combined with other BRiCs or antimicrobial resistance cassettes (ARCs). We discovered a natural example of a three-BRiC array, confirming that multiphage resistance integrons are already present in clinical isolates. Altogether, our work shows the involvement of integrons in phage defense, acting mobile and low-cost defense islands.

## RESULTS

### Integron *gcus* contain predicted phage defense systems

To obtain a broad set of *gcu*s in MIs we screened the INTEGRALL database (date: November 2021) and elaborated a curated list of 129 *gcu*s with < 95% nucleotide identity (Fig. S1). We synthesized and cloned them in pMBA, as cassettes in first position of a class 1 MI under the control of a strong Pc promoter (PcS) (*30*). The sequences of our selected collection were submitted in May 2022 to DefenseFinder and PADLOC (*32, 33*) predicting that 4 cassettes (*gcu*59, *gcu*128, *gcu*135 and *gcu*N) contained homologs of known defense mechanisms (Fig. 1A).

**Fig. 1.**
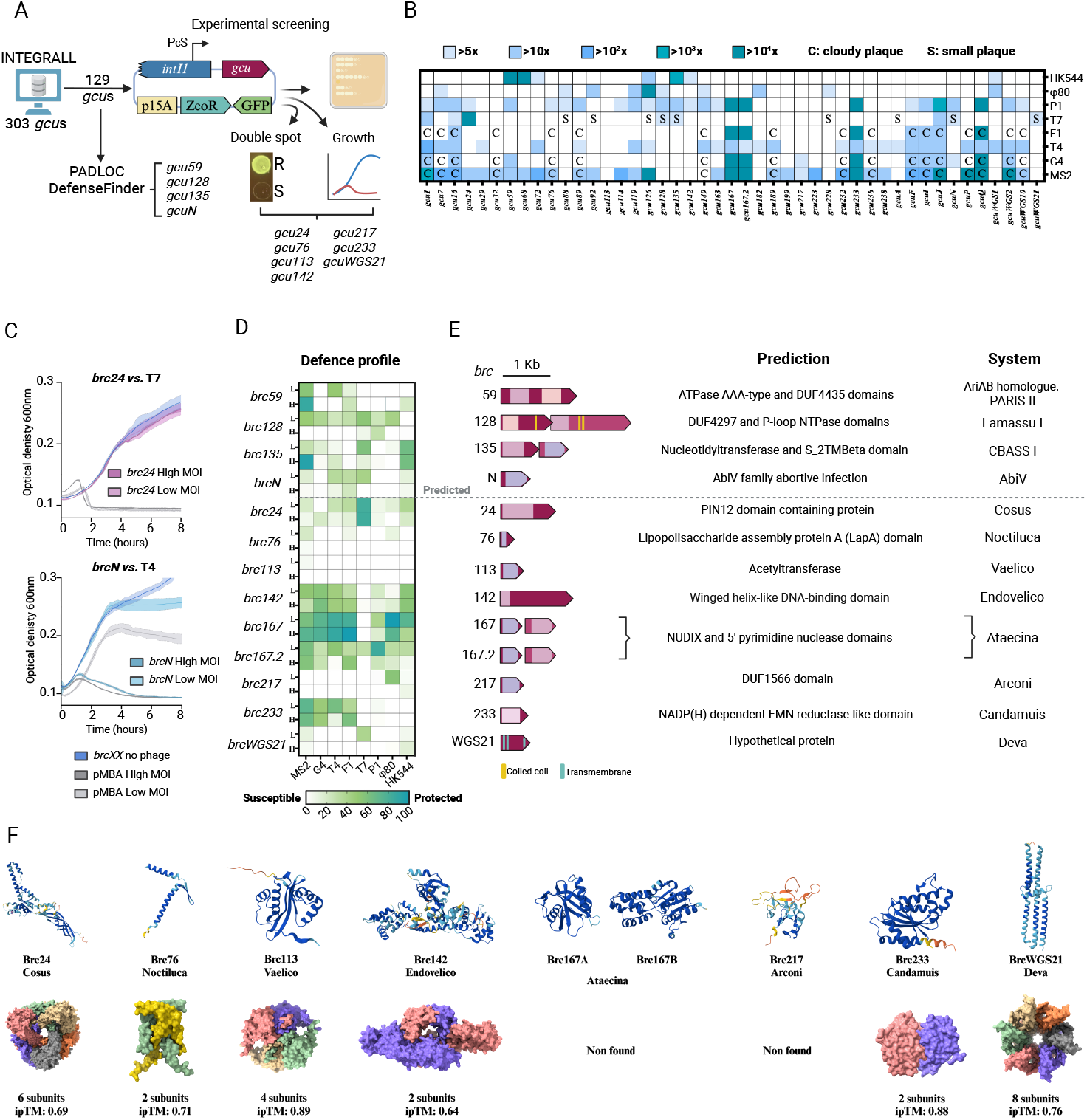
Identification and Characterization of Phage Defense Systems in Integron Cassettes. **(A)** Schematic workflow for detection of PDSs in *gcus*. A curated collection from INTEGRALL was cloned in pMBA vector. Systems were identified using PADLOC and DefenseFinder, (May 2022), double-spot assays, growth curves and plaque assays. **(B)**Profile against a panel of eight *E. coli* phages of *gcu*s providing >5-fold defense against at least one phage. **(C)** Examples of growth curves used to build the heatmap in C: *brc24* confers strong resistance to phage T7 at both high and low MOI while *brcN* shows resistance to phage T4 only at low MOI. Growth curves are represented as the mean of three independent replicates. The standard error of the mean is represented as a lighter color shade. **(D)** Heatmap showing the defense profiles at different MOIs. The degree of protection (resistance) is indicated by the color scale, with dark green representing high protection, light green representing low protection, and white no protection. **(E)** Genetic organization and predicted functions of selected *gcus*. **(F)** Tertiary and quaternary structures of novel phage defense systems, as predicted by AlphaFold3.

Given their newly identified putative function as BRiCs, we propose to rename these *gcus* as *brc*s, while conserving their initial numbers or letters for simplicity. Brc59 (formerly Gcu59) was identified as a homolog of the AriAB abortive infection system (*34*). *brc128* encodes two ORFs (*brc128A and B*) showing similarity to Lamassu Type 1 systems. AlphaFold predicts that Brc128A shares motifs with YfjL-like or AbpA proteins, involved in phage defense, while Brc128B contains a putative exonuclease domain. *brc135* also contains two ORFs encoding proteins with roles in nucleotidyl-transfer and membrane translocation and showing structural similarity with a CBASS Type 1 system (*35, 36*). Last, *brcN* encodes an AbiV family protein (*37*), implicated in abortive infection defense.

### Functional screening reveals additional phage defense systems

To search for novel PDSs that might not be detected by algorithms, we confronted experimentally the whole *gcu* collection (established in DH5α) against phages T4 or T7 using a double-spot screening protocol and growth curve monitoring (Fig. 1A and S1). We successfully detected 7 new candidates (*brc24, brc76, brc113, brc142, brc217, brc233 and brcWGS21*) that enabled growth in the presence of phage, suggesting that PDSs might be abundant among *gcu*s. To better address this, we subcloned the collection in strain *E. coli* IJ1862 (an F’ strain that can be infected by phages targeting the conjugative pilus (*38*)) and subjected it to plaque assays with phages MS2, F1, G4, T4, T7, P1, Φ80 and HK544. This screening revealed that 45 *gcu*s in the collection conferred >5-fold protection against at least one phage (Fig 1B). Because some phenotypes-like cloudy plaques-are difficult to interpret and quantitate, we selected 13 systems with clear phenotypes in the double-spot assay for further characterization, 9 of which are novel. We conducted infection assays with all phages at varying multiplicities of infection (MOIs) and monitored the optical densities (OD_600_) of bacterial cultures over time (Fig. 1C). We used the area under the growth curve compared to an empty vector control to determine the protective effect of the cassette, as in (*39*) (see Materials and Methods). Our data confirmed that all BRiCs conferred resistance to at least one phage at one MOI (Fig. 1D and Fig. S2). Resistance profiles were diverse, with examples of broad-spectrum BRiCs, and others a narrower spectrum but very high resistance against a given phage. The resistance conferred by *brcN* is limited to low MOIs confirming *in silico* predictions of an abortive infection system. Overall, our results confirm that at least one third of *gcus* are indeed phage resistance genes.

Structure modelling and domain predictions (Fig. 1E, 1F and S3) showed that most systems are novel and many do not contain recognizable domains or are of unknown function. Instead, about a third of BRiCs contained transmembrane helices, compatible with direct interference with phage infection (Fig. 1E and S3). Among the predicted domains found, some have been previously related to phage defense, like the Toll-Interleukin receptor in Brc236 (*40*), or the toxin antitoxin system (HigAB) in Brc182 (*41*); while others have roles easily related to phage resistance (mRNA splicing for Brc232 or DNA repair for Brc68) (Fig. S3). Among the selected BRiCs, Brc24 possesses a PIN12 RNA-binding domain (*42*) and potentially acts as a hexameric protein. Brc76 is a small protein of 76 amino acid with a LapA domain related to biofilm formation and forms dimers (*43*). Brc113 has a GNAT N-acetyltransferase domain (also found in BrcP (Fig. S3)). Brc142 is a putative dimeric restriction endonuclease. *brc167* and its close homolog *brc167*.*2* encode proteins with NUDIX hydrolase and nucleotidase activities with no evident multimeric structure. Brc217 contains the domain of unknown function (DUF) 1566 and no identified multimeric structure. Brc233 is possibly a dimeric NADP(H) oxidoreductase; and last, BrcWGS21 has no identified domains but contains transmembrane helices and conforms an octamer potentially forming a pore in the membrane. To adhere to the custom in the field, these novel systems are also given the name of a deity of Celtiberian mythology (Fig. 1E).

### BRiCs are *bona fide* integron cassettes

Finding defense systems in integron cassettes has important implications for the mobility of these elements. Identification of cassettes has not been straightforward until the development of IntegronFinder (*44*). INTEGRALL predates IntegronFinder. The genetic context, suggests that most BRiCs identified here are indeed located within class 1 or 2 MIs, often in pathogenic and multidrug resistant strains (like *brc217, brc113, brc24* or *brcN*) (Fig. 2A). But other, like *brc135* and *brc142*, did not have an MI context. Hence we wanted to verify that PDSs are encoded in *bona fide*-functional-integron cassettes. Integron recombination is semiconservative, involving only the bottom strand of *attC* and *attI* sites (*45, 46*). This unique feature can be detected in a suicide conjugation assay as the difference between recombination rates when the plasmid transfers the bottom or the top strand of an *attC* site to the recipient strain (*46*). We measured this for all cassettes and observed a 100-to 10.000-fold lower recombination of top strands (Fig. 2B). *brcWGS21* had extremely low recombination frequencies for the bottom strand, yet recombination of the top strand was only observed in 1 out of 6 replicate experiments, suggesting that it is probably an integron cassette, albeit with a very poor recombination site. Accordingly, the folded *attC* site of *brcWGS21* has the typical hairpin structure and extra-helical bases of these sites (Fig. S4) (*46*–*48*). Hence, our data confirms that BRiCs are *bona fide* integron cassettes.

**Fig. 2.**
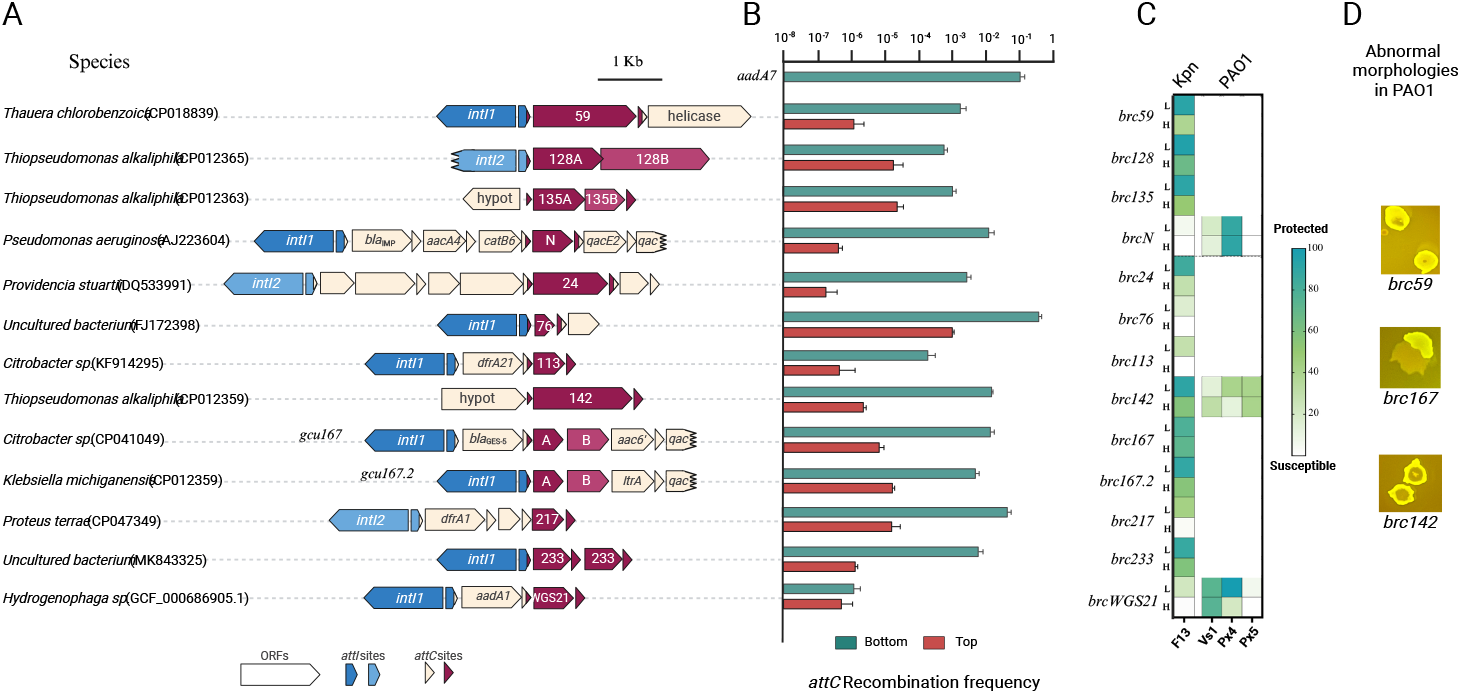
Genetic Context, Recombination Frequency, and Phage Protection Profiles of Identified BRiCs in other species. **(A)** Genetic context of BRiCs as found in databases, highlighting their association with integron integrases (*intI1* or *intI2*, in blue shades). BRiCs are found in different host species. Each BRiC is annotated with its respective identifier (e.g., 59, 128A/B, 135A/B), along with known associated genes such as resistance genes (e.g., *bla*IMP, *aacA4, catB6*). **(B)** Recombination frequency of the *attC* sites of the identified BRiCs. The recombination frequency was measured for the bottom (green bars) and top (red bars) strands of the *attC* sites using a suicide conjugation assay. Bars represent the mean of at least 3 biological replicates. Error bars correspond to the standard error of the mean. **(C)** Heatmap showing the protection profiles of BRiCs against phage infection in different host species, including *K. pneumoniae* (KP5) and *P. aeruginosa* (PAO1) against a panel of cognate phages (F13, Vs1, Px4, Px5). **(D)** Images of transformant colonies of PAO1 showing abnormal morphologies.

### Defense cassettes are protective in other species

Given the mobility of integrons and cassettes among important pathogens, we sought to determine if BRiCs are active in different hosts. To this end, we evaluated their protective effect in *K. pneumoniae* and *P. aeruginosa*, two species of the ESKAPEE group of highly resistant and lethal pathogens where integrons are prevalent. We introduced all BRiCs into *K. pneumoniae* KP5, a MDR clinical isolate that contains three plasmids (Fig. S5) and subjected them to infection with phage F13, a *Drexlerviridae* member co-isolated with the strain. All BRiCs, except *brcN*, demonstrated resistance to F13 at low MOI, and most also conferred resistance at high MOI (except *brc76, brc113, brc217*, and *brcWGS21*) (Fig. 2C). To introduce BRiCs in *P. aeruginosa* PAO1 strain, we changed the p15A origin of replication of pMBA for BBR1, and introduced a tetracycline resistance marker. After several assays, we could only transform *brcN* successfully, while other BRiCs produced no colonies at all or abnormal colony morphologies (Fig. 2D). This led us to hypothesize that some BRiCs might be toxic or extremely costly in this background. To avoid this, we changed the PcS promoter driving the expression of cassettes for a weaker version (PcW, 30-fold less active than the PcS (*49*)). This allowed to introduce *brc24, brc113, brc142* and *brcWGS21* in PAO1. We confronted these strains to a collection of 36 phages infecting PAO1 using a double spot screen. *brcN, brc142* and *brcWGS21* showed growth in presence of phages Vs1, Px4 and Px5. We confirmed their resistance phenotype monitoring growth curves (Fig. 2C). *brc142*-that had a clear phenotype against several phages both in *E. coli* and *K. pneumoniae-* confirmed its broad spectrum and host range, conferring resistance to Px5 at both MOI and to Px4 at low MOI; contrarily, *brcN* and *brcWGS21*, that had only shown low resistance against a handful of phages in *E. coli*, conferred very high resistance against phage Px4 and Vs1 in *P. aeruginosa*. Altogether, our data proves that BRiCs are active in different host species. Although some have a narrow host range, others, like *brc142* conferred resistance in the three species against most phages tested.

### BRiCs protect against prophage activation

Conflicts between defense systems and MGEs can limit their spread through HGT. Mobile integrons can reach new hosts through conjugation, so we asked if BRiCs could interfere with the activation of existing prophages, protecting the host at the cell and/or the population level. To test this, we introduced all BRiCs in lysogens of *E. coli* 594 with either HK544 or Φ80 prophages in their genomes. We induced prophage activation with mitomycin and quantitated the phage titer after 6 hours. *brc128, brc135* and *brc142* showed mild to strong defense against the activation of HK544, with 20 to 1.000 fold decreases in titers. Brc59 completely abolished the production of HK544 virions, showing >10^7^ fold protection (Fig. 3A). Resistance against Φ80 activation was generally very mild, with only *brcN, brc167*, and *brc233* producing low but statistically significant resistance levels. Interestingly, there is not a clear correlation between defense against the free phage (Fig. 1B and 1D) and prophage activation. Certain systems, like *brc135* and *brc142* confer resistance against both forms of a phage HK544; while others, like *brc128, brc167*.2 and *brc217* only conferred resistance against one. These results show that mobile integrons containing BRiCs can interfere with prophage activation, highlighting the potential protective effect at the community level, and the complex interplay between MGEs.

**Fig. 3.**
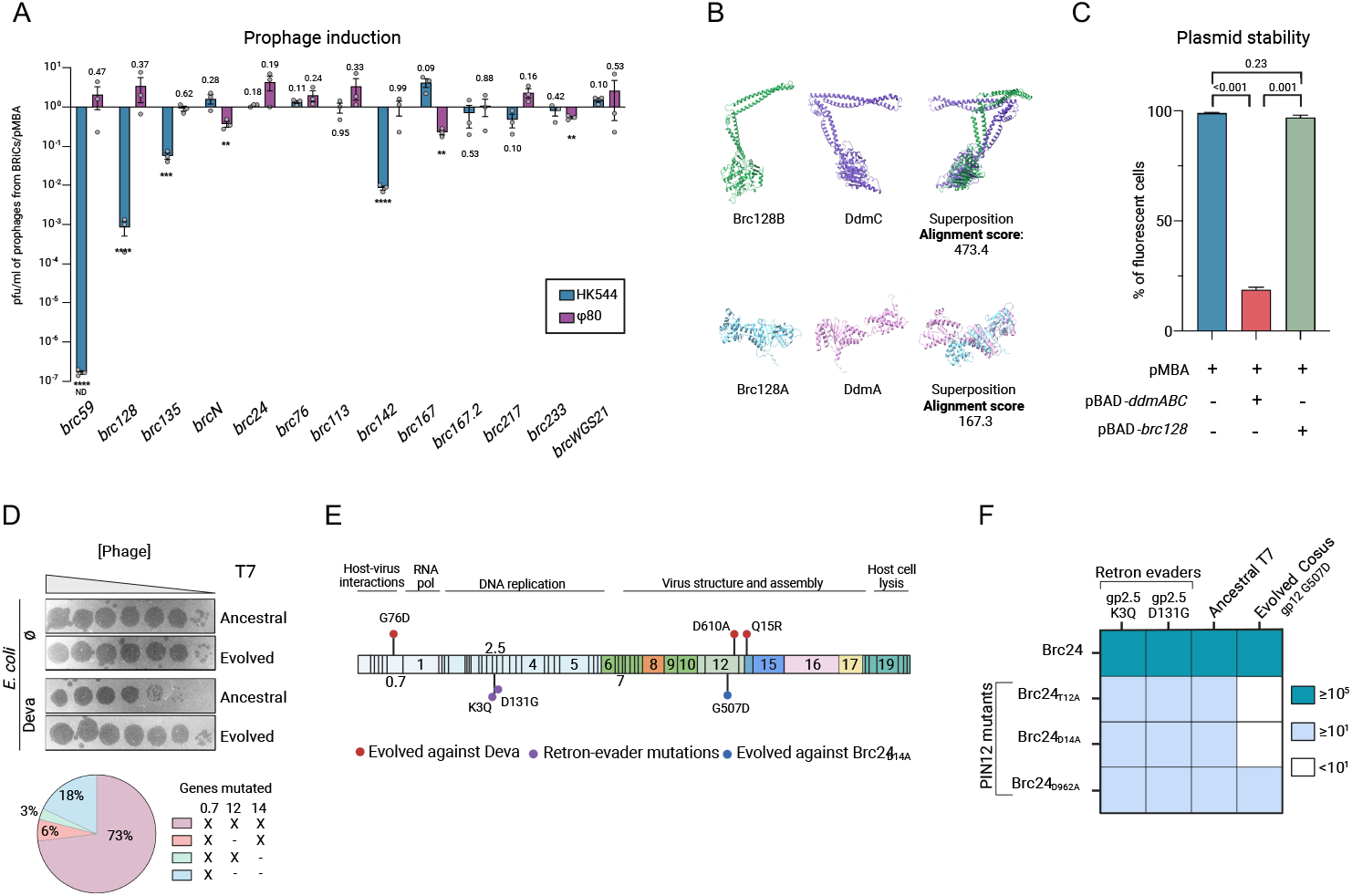
Prophage Induction, Structural Alignment, and Impact of *brc128* on Plasmid Stability. **(A)** Prophage induction measured as the ratio of plaque-forming units per millilitre (pfu/ml) in the presence of pMBA containing BRiCs over the empty pMBA. ND: not detected (Limits of detection plotted). One sample *t* and Wilcoxon test; *P* values are shown when possible; ** < 0.01; *** < 0.001; ****<0.0001. **(B)** Structural alignment and superposition of Brc128A and Brc128B proteins with DdmA and DdmC. Alignment scores indicate the degree of structural similarity. **(C)**Plasmid stability assays in *E. coli*. Percentage of fluorescent cells is used to measure the stability of pMBA. Bars represent the mean of at least 3 biological replicates. Error bars correspond to the standard error of the mean. *P* values of unpaired t-tests are shown. **(D)** Plaque assays showing T7 evasion from Deva. Mutation distribution in the population. **(E)** Location of mutations in the genome of T7. **(F)** Resistance profile of PIN12 mutants of Cosus and defense-evading phages.

### *brc128* does not have anti-plasmid activity

Brc128B shows strong structural similarities with DdmC (Fig. 3B), a Lamassu system with anti-plasmid activity in *Vibrio cholerae* (*50*). To explore if Brc128AB has anti-plasmid activity we measured the stability of pMBA (a p15A replicon) in the presence of *brc128* and *ddmABC*. After ca. 20 generations in the absence of selective pressure, pMBA was present in >97% of cells expressing Brc128AB, while only in 19% of those expressing DdmABC (Fig. 3C). We extended this study to KP5 whose three natural plasmids are distinguishable for their resistance profile (tetracycline, ertapenem or cefotaxime) and again found no effect of *brc128* in plasmid stability (Fig. S5). This shows that, despite their structural similarities, Brc128AB does not have the anti-p15A plasmid activity of DdmABC and suggests it might not be an anti-plasmid system.

### Deva interferes with genome injection

Deva (*brcWGS21*) confers a 10-fold protection and a small plaque phenotype against T7. To provide insight on Deva’s mechanism of action we evolved T7 to evade its activity. After two rounds of infection, we retrieved phages producing large plaques and similar titers in the presence and absence of Deva (Fig. 3D). Population sequencing with long reads revealed mutations in the genes encoding gp0.7, gp12 and gp14 (Fig. 3E). gp0.7 is a kinase that phosphorylates the host RNA polymerase to produce a transcriptional shutoff. The mutation in gp0.7 is pervasive: it is present in all mutant combinations, and is the only one found alone, suggesting that it is the first one to appear and that it plays a primary role in evading Deva. gp12, and gp14 are structural genes encoding tail and internal virion protein B respectively. The latter is ejected into the cell during DNA injection. Hence, mutations in gp12 and gp14 suggest that Deva interferes specifically with genome injection. This is in accordance with its probable localization in the cell membrane (see transmembrane domain and quaternary structure prediction in Fig.1E and F).

### Functional analysis of Cosus

Cosus (*brc24*) provides complete resistance against T7, with no detectable plaques in spot assays. To investigate it further, we tried to evolve phages to evade its activity, but were unable to retrieve plaques even when infecting with high phage titers (10^8^ plaque forming units (pfus)). In the absence of plaques, we tested if T7 mutants that evolved to escape other anti-phage mechanisms could evade Cosus. We tested phages with K3Q and D131 substitutions in gp2.5 (ssDNA binding protein (SSB)) (Fig. 3E) allowing to avoid retron-mediated defense (*51*) but saw no crossed evasion between systems. We changed our approach to altering Cosus and generated 11 mutants in residues predicted to be important in the activity of its RNA-binding PIN12 domain(*52*). Mutations T12A, D14A and D96A led to a strong (>10^5^-fold) decrease in resistance to only 100-fold protection, and a small plaque phenotype, supporting the correct identification of the PIN12 domain (Fig. 3F). Retron evaders did not show enhanced activity in any Brc mutant, suggesting that Cosus does not target T7 SSB. We then evolved T7 to evade Brc24_D14A_ and obtained a 5-fold increase in pfu count and plaques with an intermediate size. The evolved phage could also evade Brc_T12A_, but not Brc_D962A_ nor wild type Brc24. Genome sequencing revealed exclusively a non-synonymous mutation in the tail protein gp12 (Fig. 3E), pointing to interference with genome injection too. Taken together, our data hints at a potential dual activity of Cosus, that could explain the difficulties to evolve T7 evaders.

### BRiCs entail different fitness effects that vary across species

How defense systems affect the fitness of the host has important implications in their accumulation in genomes and their spread between species. Integrons are low-cost platforms that can modulate the expression of genes in the array by shuffling positions (*20, 21, 53*). To determine the cost of BRiCs, we performed competition assays in both *E. coli* and *K. pneumoniae* (Fig. 4). Our results showed a broad distribution of fitness effects among BRiCs, ranging from large costs (ca. 40%) to mildly positive (up to 3%) suggesting that acquisition of certain systems can be costless (Fig. 4A). It is of note that we have used a strong version of the Pc promoter, and that the cost of cassettes is probably lower in MIs with weaker Pcs. Comparison between species shows that fitness effects are not conserved between genetic backgrounds. For instance, *brcWGS21* entailed a small cost (ca. 7%) in *E. coli* but was very costly (ca. 30%) in *K. pneumoniae* (Fig. 4B). Only a few cassettes, like *brcN* showed very similar cost in both species. The lack of cost of BRiCs in certain genetic backgrounds, and the differences in fitness effects across backgrounds can be of importance in the accumulation of cassettes and their preferential distribution of BRiCs among species.

**Fig. 4.**
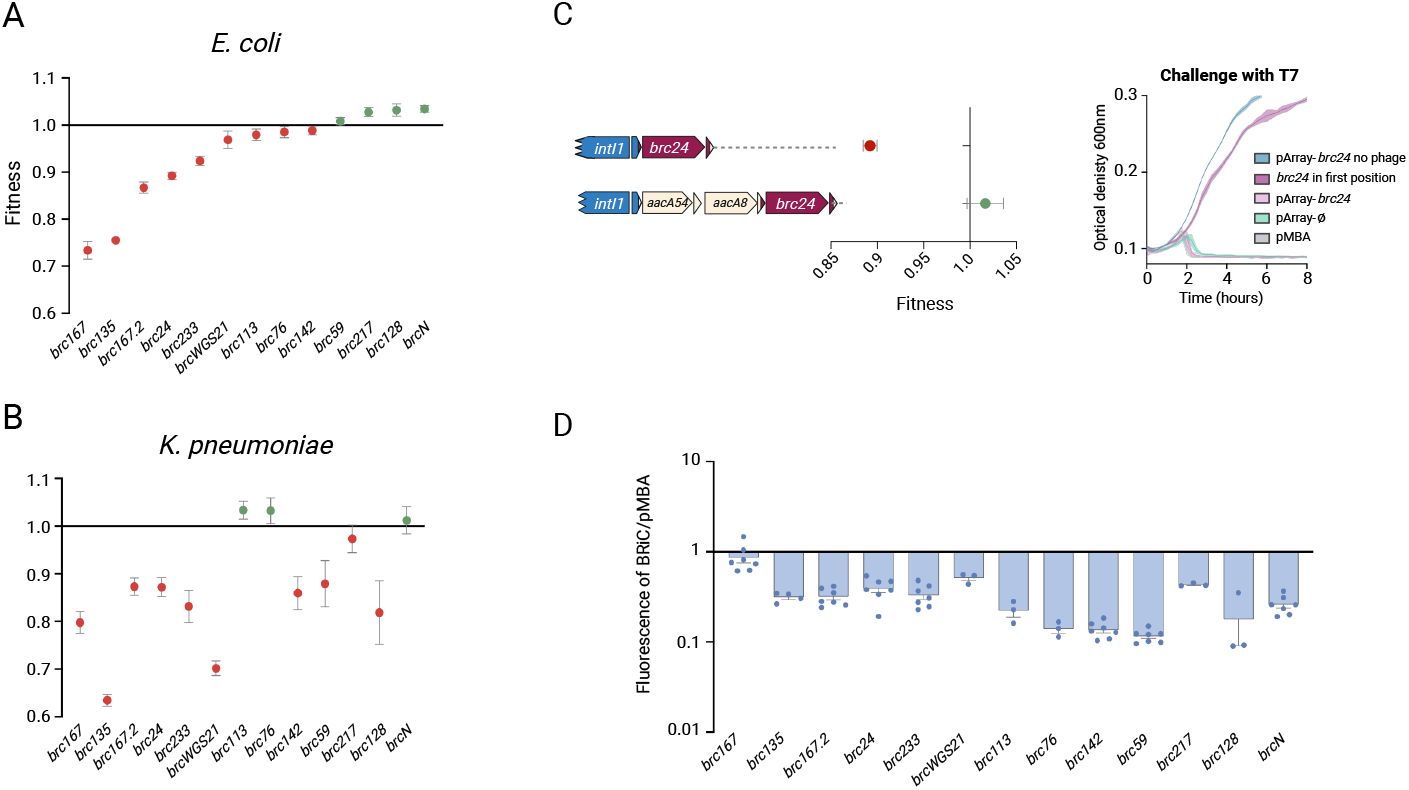
Fitness Effects of BRiCs. Fitness effects of BRiCs in **(A)** *E. coli* and **(B)** *K. pneumoniae*. Red circles represent fitness values below 1 (cost), while green circles indicate fitness values above 1 (gain). Fitness effects vary in sign and magnitude and are not conserved among genetic backgrounds. **(C)** Modulation of fitness cost and function by integron position. The graph shows the fitness effect of *brc24* in first position (competing pMBA vs pMBA-*brc24*) and third position (pArrayø *vs*. pArray-*brc24*). The fitness cost imposed by the BRiC is reduced in third position behind cassettes with strong polar effects. The cost is related to the function of the BRiC since pArray-*brc24* does not confer resistance to T7. **(D)** Polar effects of BRiCs. All cassettes except *brc167* exert polar effects on downstream cassettes, limiting their expression. All assays have been performed at least three times. Error bars represent the standard error of the mean.

### Integrons modulate the cost of BRiCs

In integrons, expression of cassettes depends on their proximity to the Pc and the polar effects that cassettes upstream can exert (*21*). Because the expression and fitness cost of a gene generally correlate, we hypothesize that integrons can modulate the cost of BRiCs. To test this, we measured the cost of *brc24* in first and third position (as it is found in the databases (Fig. 2A)) downstream of cassettes with strong polar effects (*aacA54* and *aacA8*) (*21*). Cost of *brc24* decreased from 11% in first position, to no significant cost in third (Fig. 4C). This was indeed due to the strong repression of its expression, since phage infection experiments showed pArray-*brc24* did not protect against T7 infection. Hence, costly BRiCs can be carried by mobile integrons at no cost to be later reshuffled into first position, providing phage resistance on demand (*23*). We also measured the polar effects exerted by BRiCs on downstream cassettes (Fig. 4D), showing that they too participate in the modulation of function and cost of downstream genes in the array (*21*).

### Mobile Integrons can accumulate resistance to phages and antibiotics

Being recombination platforms capturing cassettes with adaptive value, integrons could potentially combine BRiCs and ARCs to provide multi-phage and drug resistance. To test this, we combined *brc24* (T7^R^) with the *bla*OXA-10 β-lactamase and confirmed that it conferred high T7 and carbenicillin resistance separately and simultaneously (Fig. 5A). We then built an array combining *brc24* and *brc167*.*2* (P1^R^) (Fig. 5B) and showed it conferred resistance against both phages, confirming the additivity of BRiC and ARC phenotypes. Because MIs are enriched in AMR genes, natural examples of arrays combining ARCs and BRiCs are abundant (see Fig. 2A for examples). *gcus* are less abundant, so we did not find co-occurrence of BRiCs within an array. Nevertheless, having found here a variety of two-gene BRiCs, the genetic environment of *brc24* (preceded by two such cassettes) was re-evaluated. An update of DefenseFinder (June 2024) found homologs of sensor protein ThsB from the Thoeris system in the first ORF of both cassettes (*gcu23* and *gcu24*), and a type II restriction modification system downstream the last cassette (Fig. 5C). *gcu23* and *gcu24* share 50% bp identity, and a remarkably conserved predicted structure of their two ORFs. While Brc22A and Brc23A are predicted homologs of Thoeris ThsB, Brc22B and Brc23B are shorter than ThsA (300 *vs*. 500 aminoacids), show a very low protein identity (ca. 13%) and a very different predicted fold, suggesting that these genes are not homologs. The predicted structure of these proteins is instead similar to Deva (*brcWGS21*) (Fig. 1F). Both *gcu22* and *gcu23* were present in our collection, but not selected in our screenings because *gcu22* did not confer resistance to any phage, and *gcu23* showed inconsistent results and was discarded. We found that the cassette in our stock strain was frequently interrupted by the insertion of IS1, which explained the inconsistency. We interpreted this as a sign of high fitness cost, so we cloned both cassettes under a weak Pc (PcW) and tested them against the panel of phages in *E. coli* and *K. pneumoniae. gcu22* conferred low levels of resistance exclusively against phage F13 in *K. pneumoniae*, while *gcu23* had a broader defense profile, including very high resistance levels against G4, MS2, F1 and T4) (Fig. 5D). We hence redefined both cassettes as BRiCs (*brc22* and *brc23*) and, given the apparent hybrid genetic structure mentioned before, we consider this a novel system that we have called Tragantía (a deity with the upper body of a woman and the lower body of a snake). Finally, to show that this integron acts as a natural defense island we built an array containing *brc23* and *brc24* (the only ones with distinguishable phenotypes) and showed that it confers multi-phage resistance against T7 and T4 (Fig. 5E). Hence MIs naturally act as mobile defense islands that already circulate among clinical isolates (*54*).

**Fig. 5.**
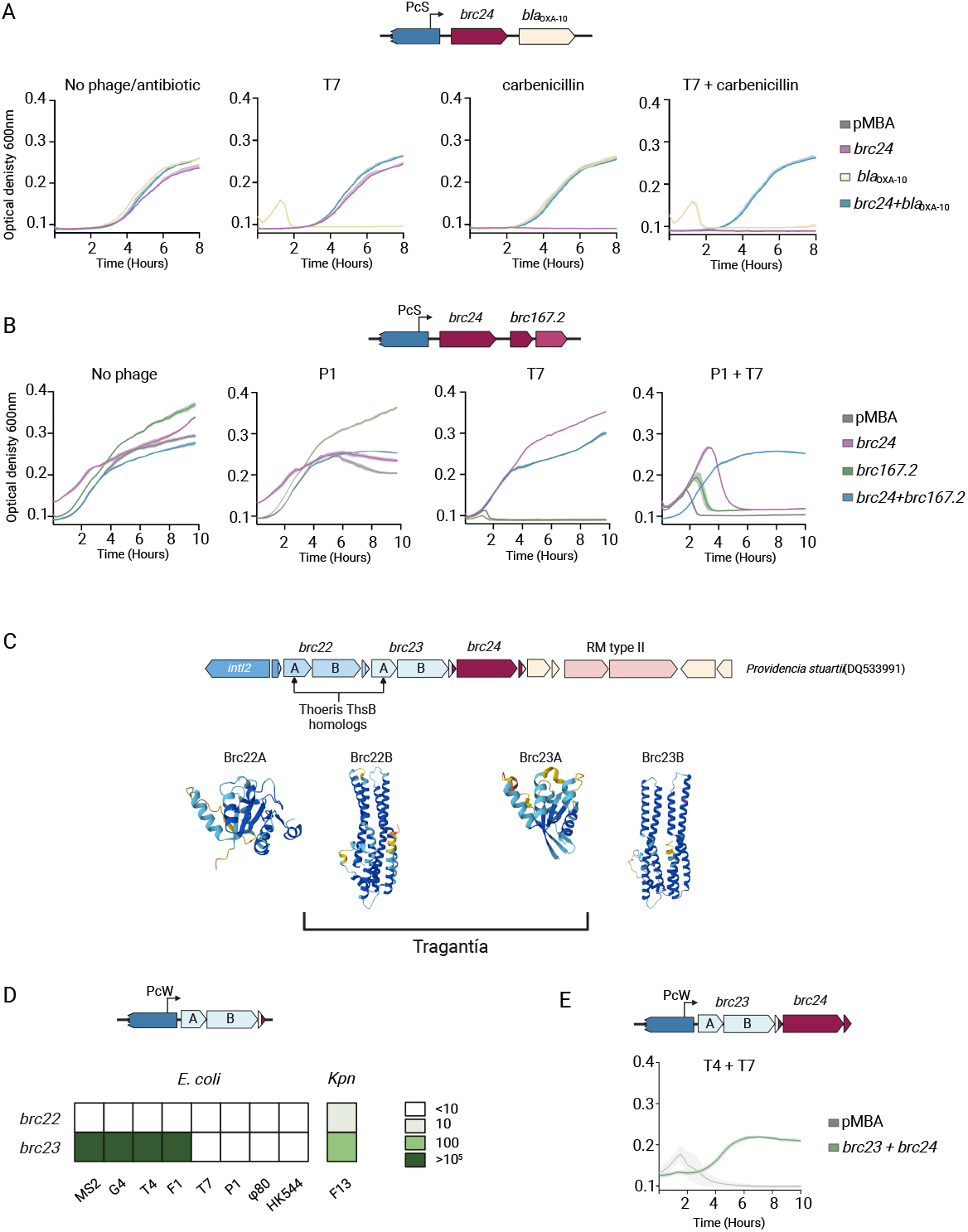
Additivity of Phage and Antibiotic Resistance Phenotypes in Multi-Cassette Arrays. Growth curves (OD_600_) of *E. coli* strains containing **(A)** pMBA, *brc24* and *bla*OXA-10 independently, and combined in an array in the absence of phage or antibiotic, and challenged with phage T7, carbenicillin, and a combination of both. The combined array confers resistance to phage and antibiotics simultaneously, and **(B)** pMBA, *brc24* and *brc167*.*2* independently, and combined in an array; either with no phage, or challenged with phage P1, phage T7, and a combination of both. (**C**) Genetic background of *brc24. brc22* and *brc23* contain homologs to TshB from Thoeris that confer resistance against phages from *E. coli* and *K. pneumoniae* (Kpn). *brc23* and *brc24* can confer multiphage resistance when in the same array. Growth curves are represented as the mean of three independent replicates. The standard error of the mean is represented as a lighter colour shade.

### Defense systems in BRiCs are found outside integrons

Integron cassettes are exchanged between integrons cohabiting the same cell. MIs can scan sedentary chromosomal integrons (SCIs) and bring to clinical settings adaptive functions evolved elsewhere in the biosphere (*55*). This suggests that BRiCs are likely found in SCIs. In an article in this issue Darracq *et al*. provide evidence that the Superintegron in *V. cholerae* contains indeed a variety of BRiCs, that are different from the ones described here (*56*). We hence sought to investigate the potential origin and distribution of our defense systems. We have searched for homologs in databases and examined their genetic context (Fig. S6). Certain systems show a broad distribution among integrons, like *brcN*, that is found in MIs and SCIs of the *Pseudomonadaceae* family (Fig. 6A), consistent with its resistance phenotype in *P. aeruginosa*. Homologs of Brc24 (Cosus) are found as BRiCs in integron arrays in *Marinobacter salexigens* (Fig. 6B). They are also found in genomes of *Vibrio* species, outside their SCIs but with a conserved *attC* site, suggesting that this could be a *bona fide* cassette integrated into an *attG* site in the chromosome (*57*). Interestingly, BrcN and Brc24 homologs were also found isolated without recognizable *attC* sites in the genomes of *Kangiella japonica* and *Escherichia marmotae* (Fig. 6C and Fig. S7). The Tragantía system in *brc22* has homologs often found in the vicinity of other PDSs. We found one in the chromosome of *Marinobacter sp*. CuT6 (Fig. 6D). This isolate has a complete integron and CALIN (cassette array lacking an integrase), but the Tragantía homolog was encoded elsewhere, within a small defense island together with homologs of Thoeris and Gao (*58*). This system shows 95% amino acid identity with Brc22A and B, but has a pseudo *attC* site in which extrahelical bases are located at both sides of the stem, and a mismatch is found in the conserved crossover point. This likely makes it non-recombinogenic (Fig. S8). While this does not seem to be a *bona fide* integron cassette (and it is not recognised by IntegronFinder), it is strikingly close (only a few mutations away) from a canonical one. Altogether, the high plasticity in the genetic context of these systems and their *attC* sites suggest a dynamic recruitment of PDSs from defense islands to integrons.

**Fig. 6.**
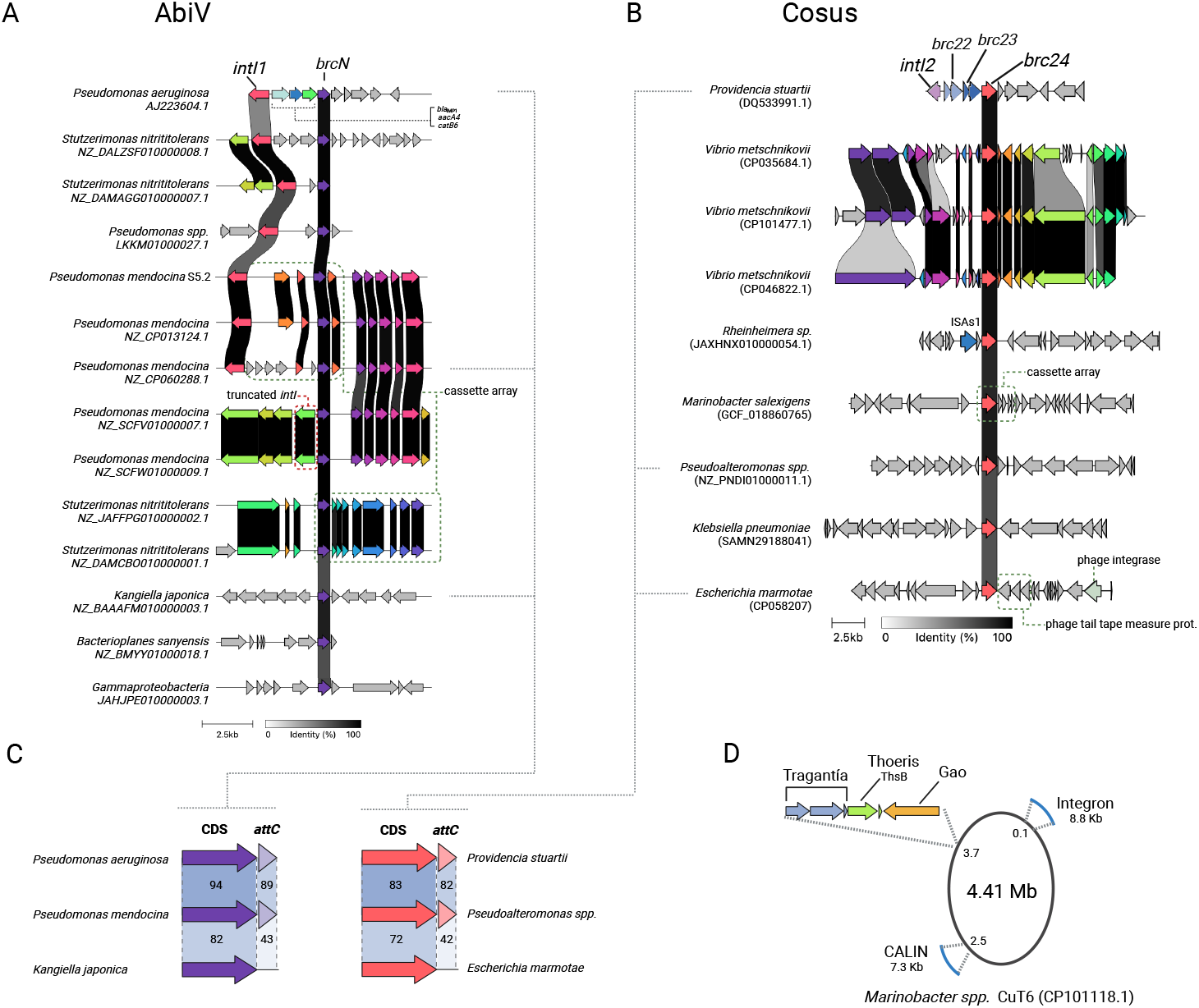
Genetic context of BRiC homologs. Analysis of the genetic environments of AbiV (*brcN*) **(A)** and Cosus (*brc24*) **(B)** showing their distribution in mobile and sedentary chromosomal integrons of different species. Integron integrases and arrays are marked where recognised. Some homologs to BRiCs are also found outside integrons, and, despite a good sequence conservation in the coding region, they lack *attC* sites (**C**). A Tragantia system, homolog of *brc22*, is found within a small defense island in the chromosome of *Marinibacter* spp. and contains a pseudo *attC* site. This isolate contains an SCI and a CALIN (**D**).

## DISCUSSION

In this work we show that mobile integrons act as highly-mobile, low-cost defense islands. We have successfully identified 47 new BRiCs experimentally, and have characterized in depth 15 of them. BRiCs displayed different specificities conferring narrow to broad immunity in our assays.

Our work is not free from limitations. First, while our screening successfully retrieved 45 BRiCs in the *gcu* collection, it is likely that others went undetected given the variety of bacterial species in which *gcu*s have been found. Indeed, it is probable that other PDSs would be discovered if we could test them in other bacterial species against their phages. This rationale could also apply to our plasmid stability assays, where testing more plasmids might reveal an anti-plasmid activity in Brc128. Nevertheless, despite the structural similarities, Brc128B and DdmC belong to different types of Lamassu systems and Brc128B does not contain the Walker B motif found in DdmC.

Hence, we cannot rule out intrinsic functional differences between both systems. In this sense, the lack of anti-plasmid activity in Brc128AB might have been selected for in conjugative plasmids navigating an environment with high antibiotic pressure. Additionally, detection algorithms are being constantly improved, so prediction of BRiCs in *gcus* will likely yield more hits at the time this work is published.

This study reshapes our perception of integrons. The presence of PDSs and AMR genes, proposes that integrons are protection elements against a large breadth of environmental insults. Our data also highlights for the first time the crosstalk between integrons and genomes with a multitude of examples of BRiCs closely related homologs outside integrons. The study of BRiCs might help reveal the genesis of cassettes, a long-standing question in the field. Being part of an integron has a profound impact in the biology and ecology of PDSs and of bacterial defense strategies. Encoded in BRiCs, PDSs become extremely mobile genetic elements that can be combined to confer multi-phage or phage-drug resistance and shuffled to modulate their cost. An extreme case of cost modulation is the Superintegron of *Vibrio cholerae* a large structure carried at no measurable cost (*59*) that contains several BRiCs, as shown by Darracq *et al*. in a paper in this issue. Additionally, their findings highlight that SCIs are extensive repositories of BRiCs for MIs.

Phage therapy is an old approach with renewed interest in the light of the AMR crisis (*60, 61*). It allows for personalized treatments against multidrug resistant bacteria, with encouraging outcomes against multidrug resistant isolates. Mobile integrons have significantly contributed to the AMR crisis bringing to our hospitals a plethora of ARCs from the genomes of environmental bacteria. Being shed to the environment at extremely high quantities (10^23^ per day) (*62*), MIs connect the genomes of pathogenic and environmental bacteria (*63*). The rampant movement of MIs among clinically-relevant bacteria ensures a rapid dissemination of any novel adaptive function in our hospitals. We have shown that BRiCs can confer resistance in 3 of the 5 Gram negative species of the ESKAPEE group of highly resistant and dangerous pathogens: the kind of bacteria aimed by phage therapy assays. This strongly suggests that the spread of BRiCs among them will be quick if selective pressure with phages becomes commonplace. This has implications in what are today considered as exploitable trade-offs in phage therapy. For instance, it is known that many *K. pneumoniae* phages bind the bacterial capsule and that resistant clones can easily arise through capsule loss, albeit at the cost of becoming non-virulent (*64*). Also, in some cases becoming resistant to phage infection through mutations comes at the cost of losing antibiotic resistance (*65, 66*). The acquisition of plasmids containing integrons with BRiCs can abrogate the exploitability of such trade-offs, rendering virulence and antibiotic resistance perfectly compatible with phage resistance. Altogether, our results highlight that the role of integrons in phage defense can be critical for the advent of phage therapy.

The role of MIs in phage resistance showcases the interplay between MGEs. Many of the BRiCs described here confer resistance against phages that target the conjugative pilus of *E. coli* IJ1862. Hence, BRiCs can be beneficial for conjugative plasmids and foster HGT by alleviating the trade-off between acquiring adaptive plasmids and becoming susceptible to a large variety of phages. Nevertheless, the fact that we couldn’t introduce many systems in *P. aeruginosa* (even under a PcW) suggests that some BRiCs can also act as barriers to HGT.

Altogether, we show that MIs can act as mobile and low-cost phage defense islands. Being able to exchange BRiCs between plasmids and to move across genetic backgrounds, MIs are likely important players in the complex interactions between mobile genetic elements.

## Acknowledgements

We would like to thank Dr. José R. Penadés, Dr. Alfred Fillol-Salom and Dr. James J. Bull for providing phages and strains, and Tamara Barcos for technical assistance. We thank other MBA lab members for critical reading of the manuscript.

## Funding

European Union’s Horizon 2020 Marie Skłodowska-Curie grant agreement No 847635 (NK)

Universidad Complutense PhD program (AH)

Juan de la Cierva program FJC 2020-043017-I (PB)

Ministerio de Ciencia, Innovación y Universidades FPU21/03268 (LOM)

Ministerio de Ciencia e Innovación Programa Ramón y Cajal RYC2019-028015-I (P.D-C).

Ministerio de Ciencia e Innovación PID2020-112835RA-I00 (P.D-C)

Ministerio de Ciencia e Innovación Subprograma Miguel Servet CP19/00104 (MG-Q),

Instituto de Salud Carlos III (Plan Estatal de I+D+i 2017–2020) (MG-Q)

European Research Council (ERC) Starting Grant [803375] (JAE);

Ministerio de Ciencia, Innovación y Universidades BIO2017-85056-P (JAE)

Ministerio de Ciencia, Innovación y Universidades CNS2022-135857 (JAE)

Ministerio de Ciencia e Innovación PID2020-117499RB-100 (JAE)

EU HARISSA JPI-AMR program PCI2021-122024-2A (JAE)

Comunidad de Madrid Programa de Atracción de Talento 2016-T1/BIO-1105 (JAE)

Comunidad de Madrid Programa de Atracción de Talento 2020-5A/BIO-19726 (JAE)

## Author Contributions

Conceptualization: JAE

Investigation: NK, AH, LOM, PB, TD, PV, FMO, TJ, MG-Q, AC, PD-C.

Funding acquisition: JAE

Project administration: JAE

Supervision: MG-Q, PD-C, JAE

Writing Original Draft: NK, AH, JAE.

Writing review and editing: NK, AH, DJ, PD-C, MG-Q, JAE.

## Competing interests

Authors declare that they have no competing interests.

## Data and materials availability

The genome of strain KP5 has been deposited in NCBI GenBank (accession numbers: CP162381-CP162384). All other data are available in the main text or the supplementary materials.

## List of Supplementary Materials

Materials and Methods

Figs. S1 to S8

Tables S1 to S3

References (*67-83*)

## Notes

### Competing Interest Statement

The authors have declared no competing interest.

### Summary of Updates

This version of the manuscript has important modifications and enhancements both in the text and in the figures

