## Supplementary for "Mobile Integrons Encode Phage Defense Systems"

#### Materials and Methods

##### Strains and Phage collection

A list of strains can be found in Table S1. *E. coli* DH5a and *P. aeruginosa* PAO1 are laboratory strains. *K. pneumoniae* KP5 is a clinical isolated at Hospital Fundación Jiménez Diaz, Madrid. *E. coli* 594 (*67*), phages HK544, Φ80 and T7 retron-evaders were kindly donated by José Penadés and Alfred Fillol-Salom. Strains were grown at 37 °C in lysogeny broth (LB) or LB agar (1.5 %) (BD, France). Zeocin was added at 100 µg/ml to maintain pMBA and pMBA-derived plasmids in *E. coli* and *K. pneumoniae*. Tetracycline at 100 µg/ml was used to maintain pMBA plasmids in *P. aeruginosa*. Liquid cultures were incubated in an Infors Multitron shaker at 200 rpm (Informs HT, Swiss). Antibiotics were purchased from Sigma Aldrich (Merck, USA) except for zeocin (InvivoGen, USA).

The coliphages included T4, T7, P1, MS2, G4, F1 which belong to the *Myoviridae*, *Autographiviridae*, *Myoviridae*, *Fiersviridae*, *Microviridae* and *Inoviridae*, respectively. *E. coli* phage MS2, F1, G4 and their host *E. coli* IJ1862(*38*) were kindly provided by from Prof. James J. Bull (University of Texas). Phages T4, T7, and P1 were purchased from the DMSZ collection (Leibniz, Germany). The *K. pneumoniae* phage, named F13, (found in effluents from a hospital) was selected for its ability to lyse a clinical isolate called KP5, which produces the class D carbapenemase OXA-48 and used as its host for this study. We computed the genomic data obtained by Illumina technology with PHASTER to determined that F13 belongs to the *Drexlerviridae* family. The *P. aeruginosa* PAO1 strain was used as a host for the Px4, Px5 and Vs1 phages used in this study.

##### Phage production

We prepared phage solutions using previously published protocols (*68*). Briefly, we grew host bacteria in LB broth at 37ºC until they reached the exponential phase. At that point, we initiated and monitored infection with the respective phage for 4 (for T7, T4, P1 phages) to 6 hours (for MS2, G4, F1, HK544, and Φ80 phages). When we observed a decrease in optical density (OD), we allowed the lysis process to continue for an additional hour. We then transferred the cultures to polystyrene Falcon tubes for centrifugation and subsequent filtration using a 0.2 mm filter (Sartorius, Spain) to collect the lysate containing the phage. Next, we tittered the phage solutions to determine the plaque-forming units (pfu). For plating, we mixed bacterial cultures ($OD_{600}$ ~1) with top agar medium and poured it onto agar dishes. After allowing it to dry for 30 minutes, we spotted serial dilutions of the phages onto the agar plates and incubated them overnight at 37ºC (*69*).

##### *In silico* analysis

We generated the list of *gcus* using the INTEGRALL database (integrall.bio.ua.pt) and Integron finder (*31*, *44*). To determine if the *gcus* associated with known phage resistance defense systems, we subjected the sequences of all *gcus* to a BLAST search against the two main defense system databases: DefenseFinder (*70*) and PADLOC (*33*). For *gcus* that showed a resistance phenotype but remained unidentified by these platforms, we used an AlphaFold2 pipeline known as ColabFold for prediction, followed by Foldseek to gain insights into the protein function of each *gcu* (*71*, *72*). Additionally, we employed the Iterative Threading ASSEmbly Refinement (ITASSER) platform to leverage complementary approaches for protein function prediction (*73*). Genetic environments were analyzed with CAGECAT (cagecat.bioinformatics.nl) (*74*). For oligomerization experiments, we used the Alphafold server from Google DeepMind(*75*). Multimers were selected when ipTMs were >0.5. Visualization of oligomer structures were obtained by using the ChimeraX software.

We carried out the sequencing of the *K. pneumoniae* isolate KP5 using a combination of Illumina and Nanopore technologies, with SeqCoast (Portsmouth, USA) performing the sequencing. We performed genome assembly using Unicycler software and conducted subsequent analyses using the Center for Genomic Epidemiology platform. We determined the resistome using the ResFinder web platform, identified the sequence type using MLST, and characterized the plasmids using the PlasmidFinder web platform (*76*–*78*). We assessed the presence of prophages in the genome using the PHASTER web tool (*79*). General DNA sequence analysis was performed with Geneious and figures were created with BioRender.com.

Cloning *gcu*s in pMBA

All *gcus* were synthesized as double-stranded DNA by IDT ((Newark, USA), with an addition of a 20 bp homology region corresponding to the targeted cloning site of pMBA located between the *intI1* and *gfp* gene (*30*). We performed the cloning process using the Gibson assembly method (*80*) and selected clones on an LB agar plate supplemented with zeocin (50µg/mL). We cloned the entire sequences of the *gcus*, including the open reading frame (ORF) and their *attC* region, into the pMBA vector. Our in-house designed pMBA plasmid contains a p15A replication origin, a zeocin resistance marker, and a truncated *intI1* gene followed by a *gfp* gene, acting as a second integron cassette. We inserted the cloned *gcus* as the first cassette in the vector. To ensure a relevant level of expression for each *gcu*, we selected a strong version of the Pc promoter, PcS, located within the *intI1* gene, for our constructions. For the construction of multicassette arrays, we cloned at second position of the integron-like structure of pMBA the *bla*$_{\text{OXA-10}}$ gene or the *brc167.2* instead of having the *gfp* gene (strains C916 and D159 respectively). For the construction of the example of multiphage defense array found in databases we cloned *brc23* and *brc24* at first and second position, respectively, of pMBA under the control of a PcW promoter. To test polar effects and fitness cost, *aacA54* and *aacA8* from (*21*) and *brc24* were cloned into the pMBA-Δ*gfp* vector using Gibson assembly. For the cloning of *gcus* into *Pseudomonas aeruginosa* PAO1, we used a modified pMBA vector (pBTZ). The p15A replication origin was replaced with the BBR1 origin to ensure replication in both *E. coli* and *P. aeruginosa* species. Additionally, the strong PcS promoter was substituted with its weaker version, PcW. Finally, a tetracycline resistance marker was introduced, as *P. aeruginosa* exhibits high resistance to zeocin. The list of oligonucleotides used can be found in Table S2. The sequences of BRiCs can be found in Table S3.

Double Spot Screening

To identify new phage defense integron cassettes, we employed both liquid and solid media screening strategies. Initially, we cultured clones carrying various *gcus* overnight in LB broth with zeocin (50 µg/mL). The following day, a 1 OD dilution of each culture

(10 µL) was spotted onto LB agar plates supplemented with zeocin. After 1 hour of incubation, we applied 7 µL of phage solution at a concentration of $10^9$ pfu/mL to the bacterial spots. After 16-18 hours of incubation, we selected the clones that exhibited growth despite phage presence for further analysis. Additionally, we conducted a high-throughput screening in 96-well plates, challenging the entire *gcu* collection with phages T7 and T4 under two different multiplicities of infection (MOI) conditions.

Plaque assays

We mixed bacterial cultures ($OD_{600}$ ~1) with top agar medium and poured it onto agar dishes. After allowing it to dry for 30 minutes, we spotted serial dilutions of the phages onto the agar plates and incubated them overnight at 37ºC (*69*).

Growth curves and phage lysis monitoring

We inoculated a single colony of a clone carrying a potential phage resistance gene into an overnight culture of Luria Bertani (LB) broth at 37ºC. The next day, we adjusted the culture to an optical density of 0.1 and transferred it to a 96-well plate containing LB broth supplemented with zeocin (50 µg/mL). We then exposed the plate to different concentrations of phage to test various multiplicities of infection (MOI). For all tested phages, we defined a high MOI condition as an MOI of 1 and a low MOI condition as an MOI of 0,01. However, for phages T4 and T7, which exhibit high lytic activity, we set the high MOI condition to 0,01 and the low MOI condition to 0,00001. We monitored growth curves using a Biotek HTX synergy plate reader (Agilent , USA) at 600 nm over a 16-hour period. To assess the phage lysis level, we calculated the area under the curve (AUC) for each growth condition, both with and without phage as described previously (*39*). The inhibition score for each sample was calculated using the following formula:

$$Inhibition\ Score = (\frac{\Delta AUC}{\Delta AUC_{no\ phage}}) \times 100$$

where ΔAUC is the difference in AUC between the conditions with and without phage. Specifically, $\Delta \text{AUC} = \text{AUC}_{no\ phage} - \text{AUC}_{phage}$. We calculated the inhibition scores for both high and low multiplicity of infection (MOI) conditions. These scores were then normalized by comparing them to the inhibition scores obtained from the control samples, which used the empty pMBA vector. The normalization was done by computing the ratio of the inhibition score of the experimental sample to the inhibition score of the control: $Inhibition = \left(\frac{Inhibition\ Score_{pic}}{Inhibition\ Score_{pMBA}}\right) \times 100$. By using the "inhibition score", we normalize the reduction of the area under the curve relative to growth without the phage. This allows for comparison of the relative effects regardless of initial growth differences between strains. The protective effect of the BRiC is shown as the inverse of the inhibition score.

Prophage activation assay

We chemically activate the lytic cycle of the prophages HK544 and Φ80 in E. coli 594::pMBA-*brc* using mitomycin C at a concentration of 0,5 µg/mL. Briefly, we adjusted the culture to an optical density of 0.1, and after 1h we induced prophage activation by adding mitomycin C. Then, we quantitated the production of active phages after 6 hours of induction. To do so, we centrifuged the culture we performed serial dilutions of the supernatant and spotted it on top agar plate containing IJ-pMBA to evaluate the phage concentration (*81*).

Plasmid stability experiment

We co-transformed *E. coli* IJ1862 with both the empty pMBA plasmid -carrying the p15A replication origin and a GFP gene- and the L-arabinose inducible pBAD plasmid (pBR322 *oriV*) carrying either the *brc128* or *ddmABC* defense system. For five days, three replicates were put in culture in LB broth supplemented with carbenicillin (100 µg/ml) and L-arabinose (0,1%). Daily, we transferred 50 µl of each culture into fresh 5 ml LB broth and used another 50 µl to count GFP-expressing colonies on LB agar plates supplemented with carbenicillin. The presence of pMBA was assessed as the number of fluorescent cells in the mixture.

To assess the stability of the endogenous clinical plasmids in KP5 *K. pneumoniae* isolate, we cultivated clones harboring *brc128* in LB broth with zeocin (50 µg/ml) over five days. On days 1 and 5, we selected >120 colonies from LB zeocin agar plates and subcultured them onto separate LB media containing either tetracycline (15 µg/ml), cefotaxime (50 µg/ml), or ertapenem (30 µg/ml). This step aimed to detect any loss of resistance, which would suggest the corresponding loss of one of the three natural plasmids.

Phage evolution

Overnight cultures of *E. coli* IJ1862 or DH5α strains carrying BrcWGS21 and $Brc24_{D14A}$, respectively, were diluted 1:50 in LB + zeocin and grown at 37°C until reaching an $OD_{600}$ of 0.3-0.4. Then, 50µl of these cells were mixed with 100µl of different dilutions of a T7 lysate and left at room temperature for 10 minutes to allow adsorption. The mixture was then added to 3ml of LB top agar (0.4%) and poured in a LB agar plate which was left to dry at room temperature for 10 minutes. Plates were incubated overnight at 37°C. Next day, phage plaques were recovered in 3mL of LB and centrifuged at 4402 x *g* for 5 minutes. The supernatant was filtered using a 0.2 mm Minisart filter unit. The filtered lysate was then used for a new round of infection until the evolved T7 phage displayed increased infectivity in strains carrying the different BRiCs (i.e. increased titer on drop assays or displaying bigger plaques). The genomic DNA of evolved T7 phages was extracted and sequenced as linear DNA by Plasmidsaurus. Analysis of mutation distribution for Deva evaders was performed by using reads in which phasing of all mutations was possible.

Extraction of phage genomic DNA

To extract genomic DNA from phages, 360µL of a high titer phage lysate (>$10^{10}$ pfu/mL) were treated with 1uL DNAseI (1U) at 37°C for 30 minutes to eliminate host DNA. Following this, 20µl of 0.5M EDTA, pH 8.0 were added and samples were incubated at 65°C for 10 minutes to inactivate DNAseI. From this point, the DNeasy gDNA Extraction Kit (Qiagen) was used to isolate DNA. 180µl buffer ATL and 20µl of proteinase K were added to 200µl of DNAseI-treated lysate and incubated at 56°C for 1 hour. Then, 40µl of

RNaseA (10mg/ml) were added and samples incubated at RT for two minutes. After this step, DNA extraction was carried out following the instructions of the manufacturer. DNA was eluted in 35µl DNAse/RNAse free water.

Site-directed mutagenesis of *brc24*

To obtain the Brc24 single mutants, we designed primers carrying the desired single amino acid substitutions (D6A, T12A, D14A, E42A, D47A, S57A, D96A, D120A, D146A, D153A, and D177A) and performed Gibson assembly using the pMBA-*brc24* plasmid extracted from *E. coli* DH5α as a template. Individual transformants were screened by PCR, and the presence of the intended mutations was confirmed by DNA sequencing. The resulting mutants, along with the wild-type Brc24 and a control lacking Brc24, were then tested for susceptibility to phage T7 by spotting method, ensuring that the observed susceptibility was directly attributable to the single amino acid substitution.

*attC* recombination experiments

Recombination experiments were conducted as previously described (*82*). Briefly, we cloned the 3' region of each *brc* using a pSW23T mobilizable plasmid containing a chloramphenicol resistance marker into *E. coli* β2163 strain (*dapA*$^-$, *pir*$^+$) requiring diaminopimelic acid (DAP) to grow. We used *E. coli* β2163 as donor strain for conjugation experiments using *E. coli* DH5α strain carrying both a p3938 plasmid (Carb$^R$) containing an L-arabinose inducible *intI1* gene and a p929 plasmid (Kana$^R$) containing an *attI* site as recipient. We performed conjugation experiments onto LB agar supplemented with DAP and L-arabinose (0,2%) overnight. The next day, after cell recover, we performed ten-folded dilutions onto LB agar supplemented with carbenicillin (100 µg/ml) and glucose (1%) or with carbenicillin, glucose and chloramphenicol (25 µg/ml) to select for the total population and recombinants, respectively.

Fitness experiment

To assess the fitness cost of each BRiC, we performed *in vitro* competition assays following the protocol described in (*83*) with some modifications. Briefly, we subcloned the panel of the 13 BRiCs into a modified version of the pMBA vector, which includes a knock-out *gfp* gene (with a stop codon instead of the start codon), along with an empty vector control. We then subjected this new set of clones to a fitness competition experiment against the wild type pMBA vector carrying the intact *gfp* gene. Briefly, we grew six replicates of each clone overnight at 37ºC in a 96-well plate filled with LB broth supplemented with zeocin (50 µg/ml). The next day, we mixed cells containing the target *gcu* in the pMBA-Δ*gfp* in a 1:1 volume ratio with the clone carrying the pMBA-WT vector. We diluted these mixtures 1:20 in 96-well plates containing NaCl 0,9%. This plate was further diluted 1:20 in a plate with NaCl and another with LB broth supplemented with zeocin. The first plate was used to estimate the percentage of pMBA-WT and pMBA-*brc*-Δ*gfp* cells in the mixture on day 1, using a Cytoflex S cytometer (Beckman Coulter, USA). The second plate (with LB) was incubated overnight. The next day these cultures were diluted 1:400 in NaCl to determine the proportion of strains in the mixture as explained above. Following the formula described in (*83*), fitness cost was calculated as the difference in ratios of GFP-fluorescence between day 1 and day 2 for each BRiC.

Quantitation of polar effects

To determine the polar effects of cassettes, we measured the fluorescence of the Green Fluorescent Protein (GFP) that acts as a second cassette in pMBA. At least three independent colonies of each pMBA derived strain were inoculated in LB zeocin and incubated at 37ºC overnight. Cultures were then diluted 1/400 in filtered saline solution to measure fluorescence using a Cytoflex-S flow cytometer (Beckman Coulter). The 488 nm laser was used to detect GFP expression through 525/40 nm (FITC) band pass filter. 20,000 events were recorded per sample. Data were analysed using CytExpert software (v.2.4; Beckman Coulter).

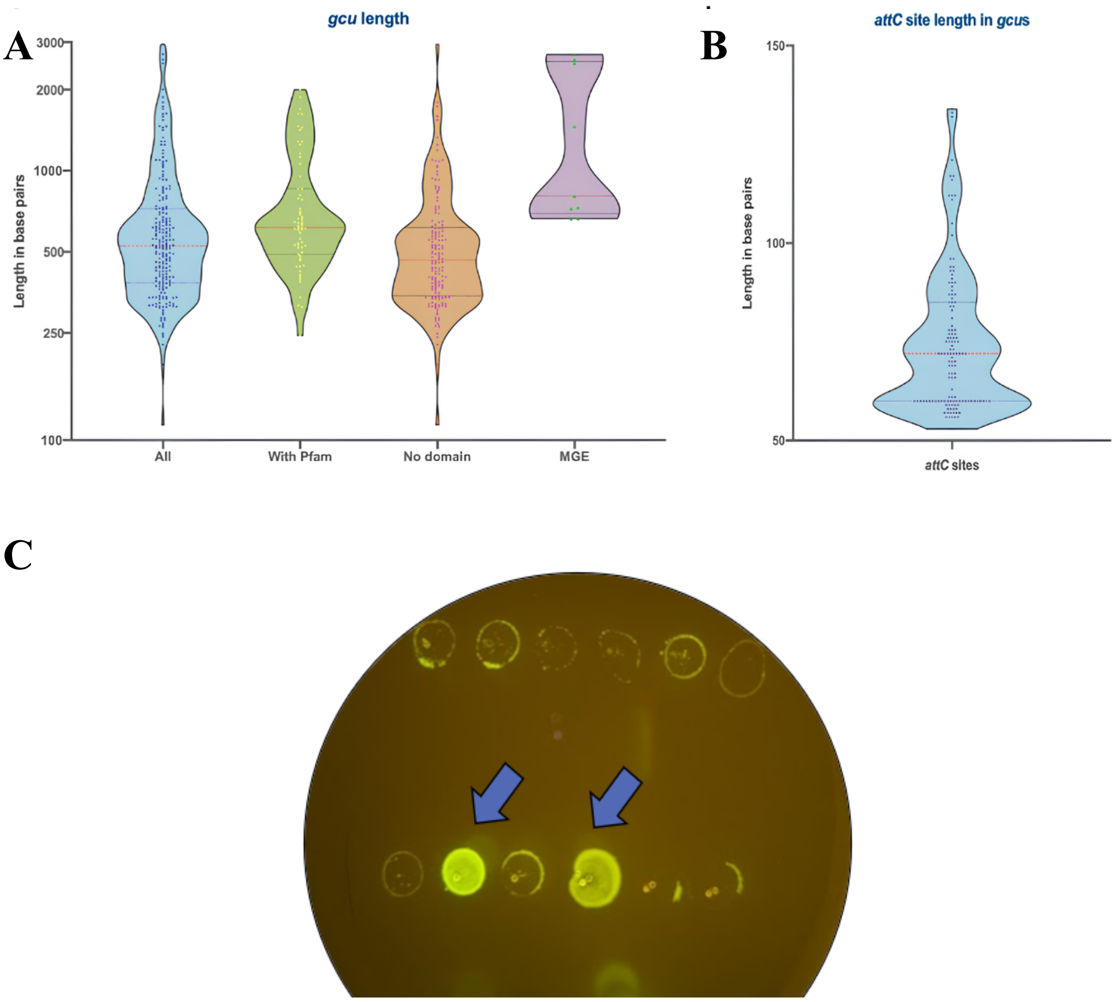


**Fig. S1. Length distribution of *gcus* and *attC* Sites**. **(A)** Violin plot depicting the length distribution of *gcus* in base pairs across various categories. **(B)** Violin plot showing the length distribution of *attC* sites in *gcus*, measured in base pairs. The *attC* sites are essential for site-specific recombination mediated by integrases. **(C) Double spot screening for phage resistance** Identification of *gcu24* and *gcu142* through double spot screening. The image shows the results of a phage resistance assay, where bacterial spots containing different *gcus* were challenged with phage T4. The arrows indicate the spots corresponding to *gcu24* and *gcu142*, which show growth despite the presence of the phage, indicating successful identification and resistance.

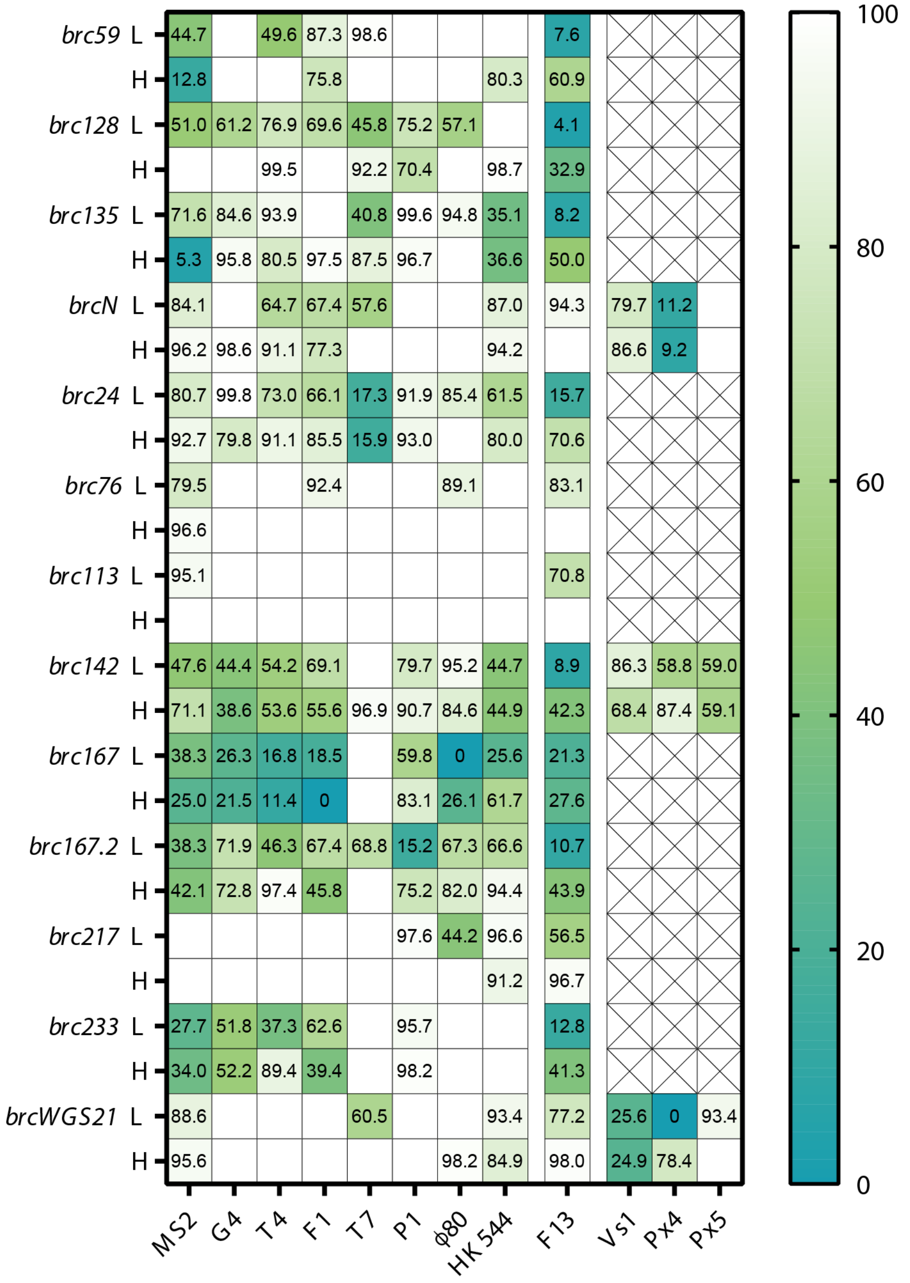


**Fig. S2. Heatmap of defense profiles across different phage types**. Heatmap showing the defense profiles of identified Bacteriophage Resistance integron Cassettes (BRiCs) against a panel of different phages. Each cell represents the level of inhibition, with 100% indicating total susceptibility (death) and 0% indicating total protection. The colour intensity reflects the degree of inhibition, where dark green represents high protection and blue indicates lower protection. The defense profiles are shown for various BRiCs against multiple phages, demonstrating the specificity and broad spectrum of phage resistance conferred by different *gcus*.

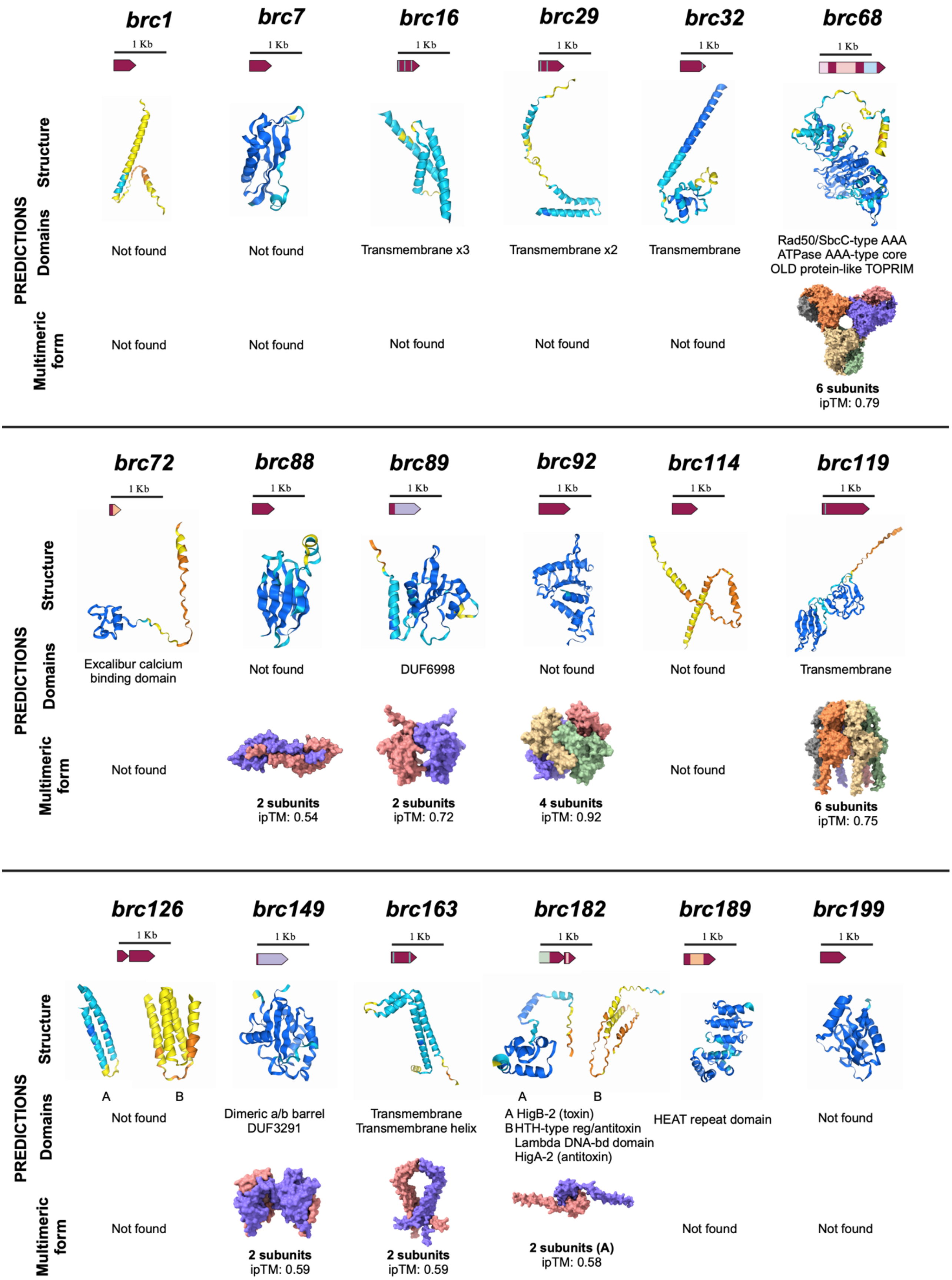
PREDICTIONS
Structure
Domains
Multimeric form
brc1
1 Kb
Not found
Not found
brc7
1 Kb
Not found
Not found
brc16
1 Kb
Transmembrane x3
Not found
brc29
1 Kb
Transmembrane x2
Not found
brc32
1 Kb
Transmembrane
Not found
brc68
1 Kb
Rad50/SbcC-type AAA
ATPase AAA-type core
OLD protein-like TOPRIM
6 subunits
ipTM: 0.79
brc72
1 Kb
Excalibur calcium binding domain
Not found
brc88
1 Kb
Not found
2 subunits
ipTM: 0.54
brc89
1 Kb
DUF6998
2 subunits
ipTM: 0.72
brc92
1 Kb
Not found
4 subunits
ipTM: 0.92
brc114
1 Kb
Not found
Not found
brc119
1 Kb
Transmembrane
6 subunits
ipTM: 0.75
brc126
1 Kb
A
B
Not found
Not found
brc149
1 Kb
Dimeric a/b barrel
DUF3291
2 subunits
ipTM: 0.59
brc163
1 Kb
Transmembrane
Transmembrane helix
2 subunits
ipTM: 0.59
brc182
1 Kb
A
B
A HigB-2 (toxin)
B HTH-type reg/antitoxin
Lambda DNA-bd domain
HigA-2 (antitoxin)
2 subunits (A)
ipTM: 0.58
brc189
1 Kb
HEAT repeat domain
Not found
brc199
1 Kb
Not found
Not found

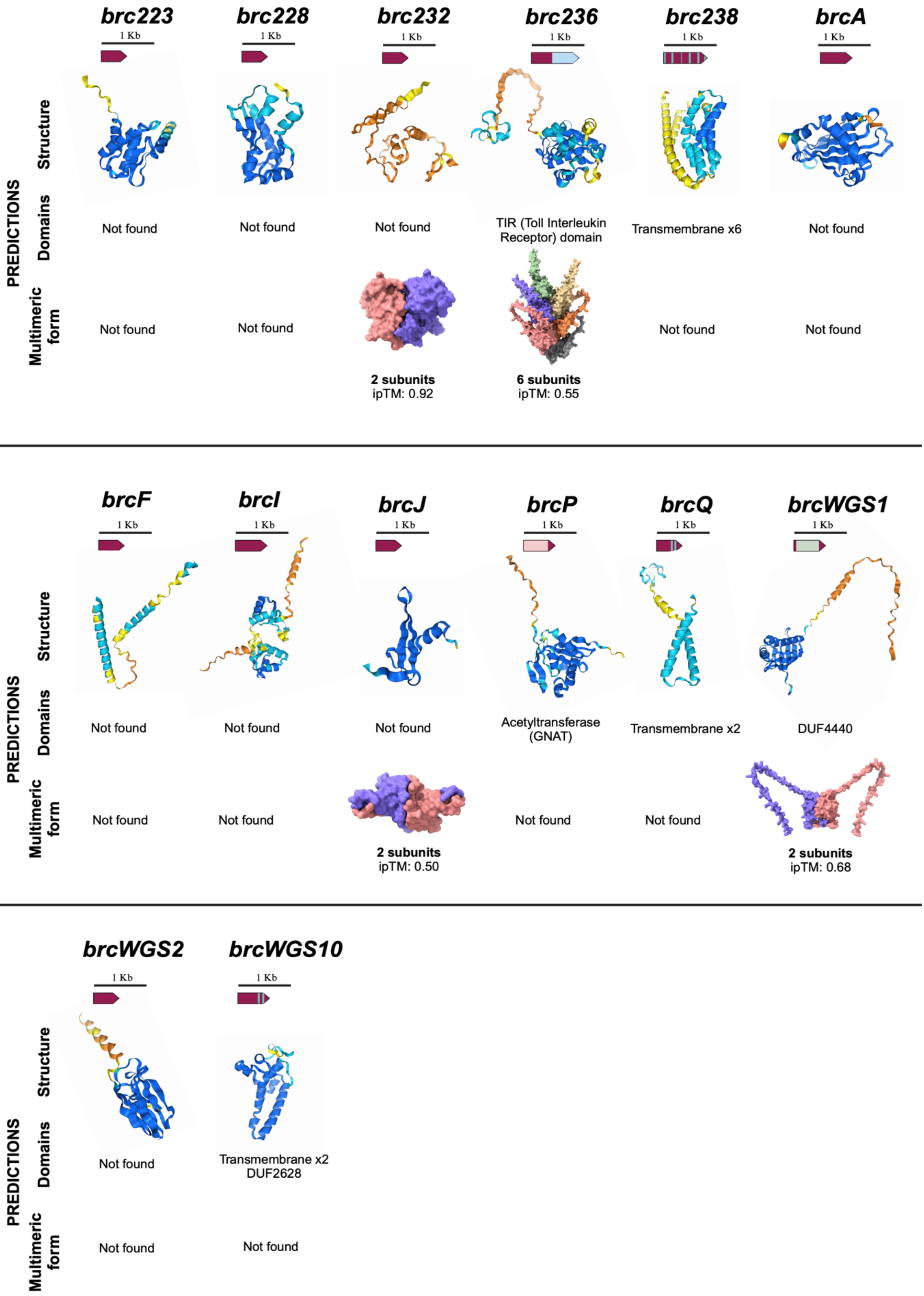


**Fig. S3. *gcus* with bacteriophage defense activity: *brcs*.** Genetic organization, and predicted function and domains of selected *gcus*. Tertiary and quaternary structures were predicted using AlphaFold3 and visualized with ChimeraX.

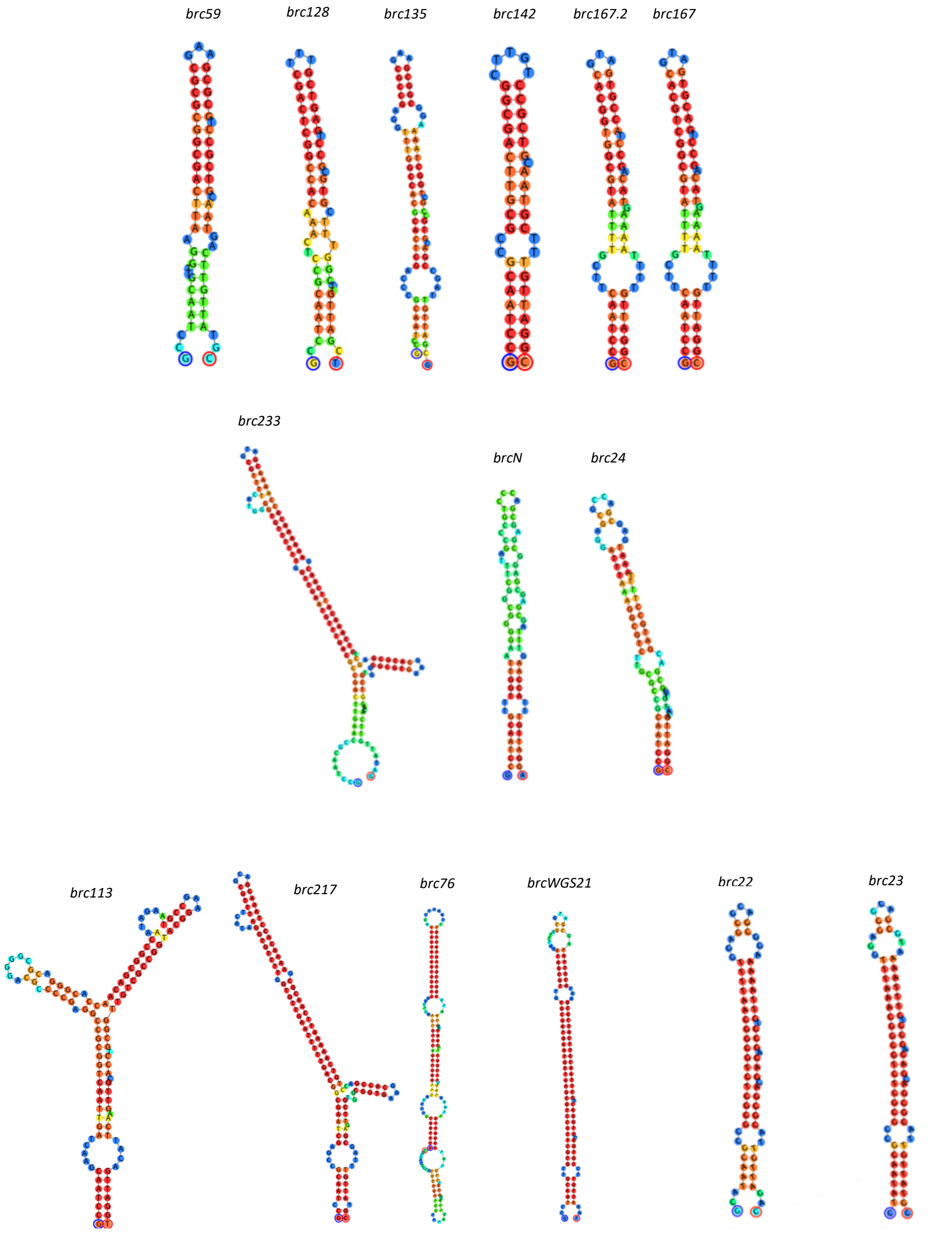


**Fig. S4. Predicted secondary structures of *attC* sites in identified BRiCs**. Secondary structures of attC sites from various BRiCs are shown. Each structure represents the predicted folding pattern for the *attC* sites associated with each *brc*. The structures are color-coded based on nucleotide positions, with blue representing the outer loops and red indicating the stems of the hairpins. These structural predictions highlight the diversity and complexity of *attC* site configurations, which are critical for site-specific recombination events mediated by integrases.

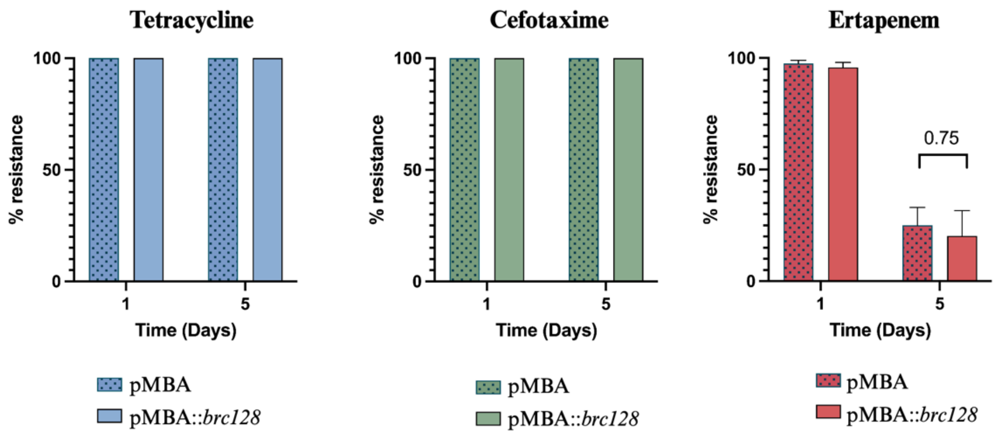


**Plasmid content of KP5 *K. pneumoniae* isolate.**

| Name | Plasmid type | Resistome |
|---|---|---|
| Kp5 | IncR (38kb) | *bla*$_{OXA-1}$, *bla*$_{TEM-1}$, *bla*$_{CTX-M-15}$, *sul2* |
| | IncL (63kb) | *bla*$_{OXA-48}$ |
| | IncH1B(200kb) | *sul1*, *dfrA15*, *aadA1*, *tet(A)* and *catA1* |

**Fig. S5. Stability of the three natural plasmids found in KP5.** Stability of the plasmids encoded in KP5 was followed over ca. 30 generations (5 days) in the presence and absence of *brc128.* Three replicates of each strain were passaged in the absence of antibiotic and plated at every time point in LB without antibiotics. Plasmid stability was measured replicating 40 colonies of each with and without the corresponding antibiotic. Plasmids conferring tetracycline and cefotaxime resistance were 100% stable in all cases. Plasmid conferring ertapenem resistance was not stable, but the differences between the empty pMBA and pMBA harbouring *brc128* were not statistically significant. Bars represent the standard error of the mean.

### PARIS II

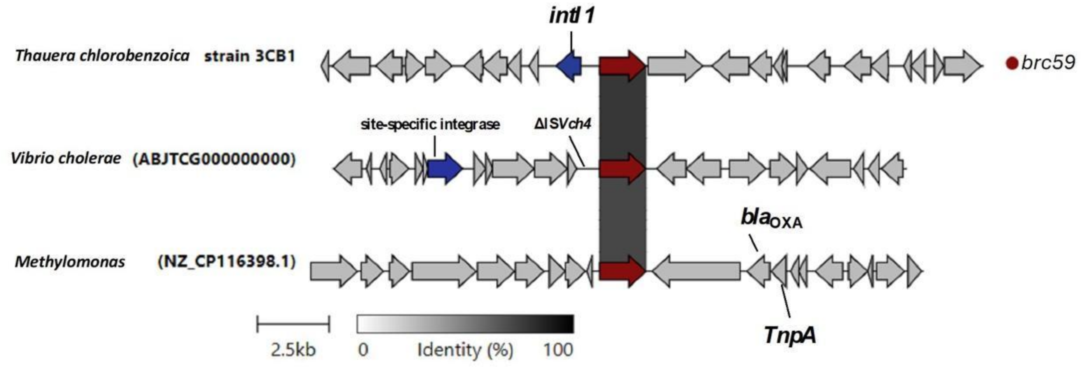


### Lamassu I

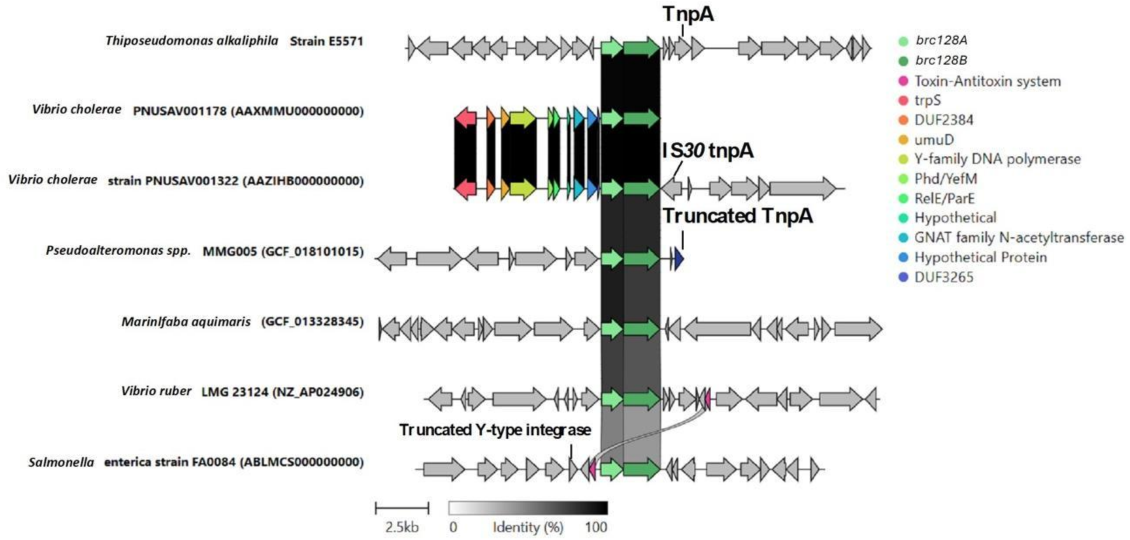

#### CBASS I

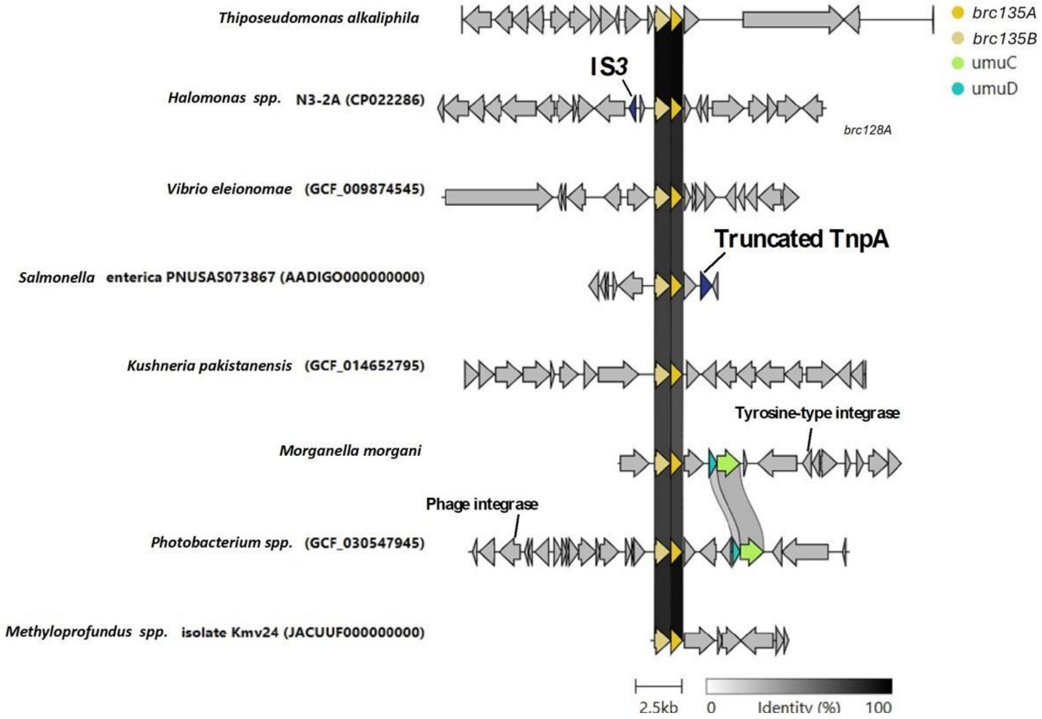


#### Ataecina

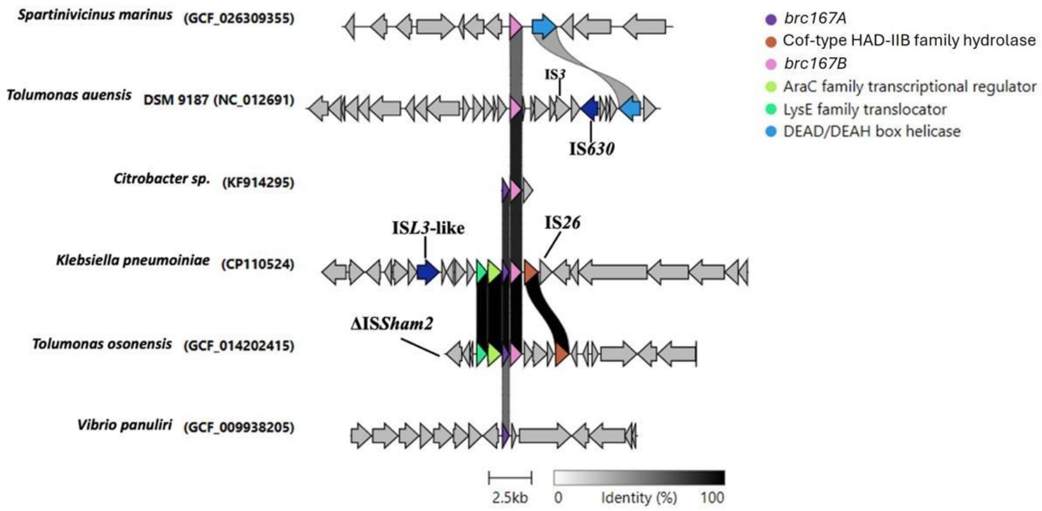

#### Endovelico

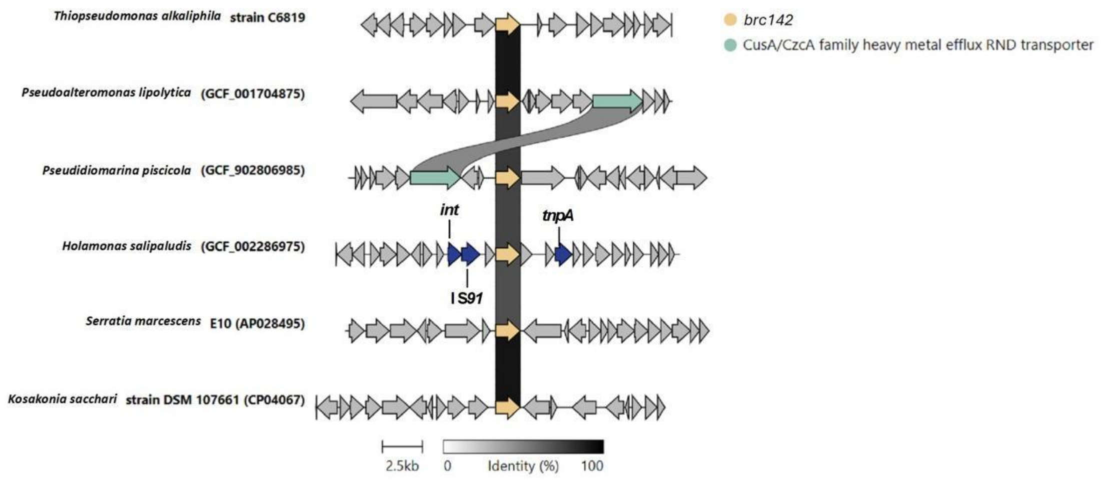


#### Cosus

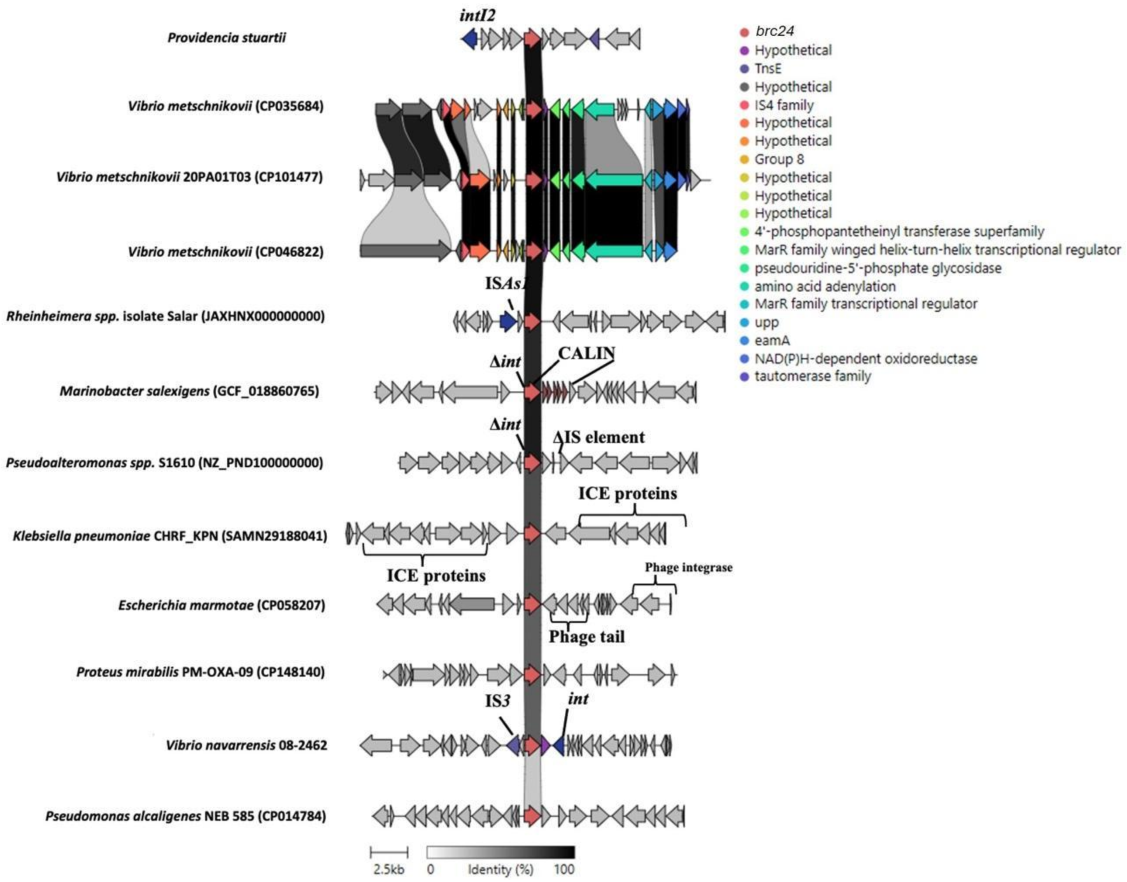

### Candamuis

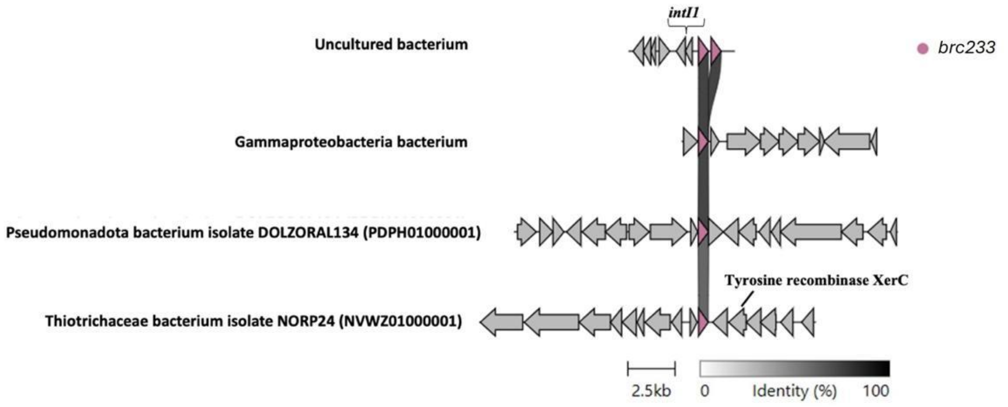


### AbiV

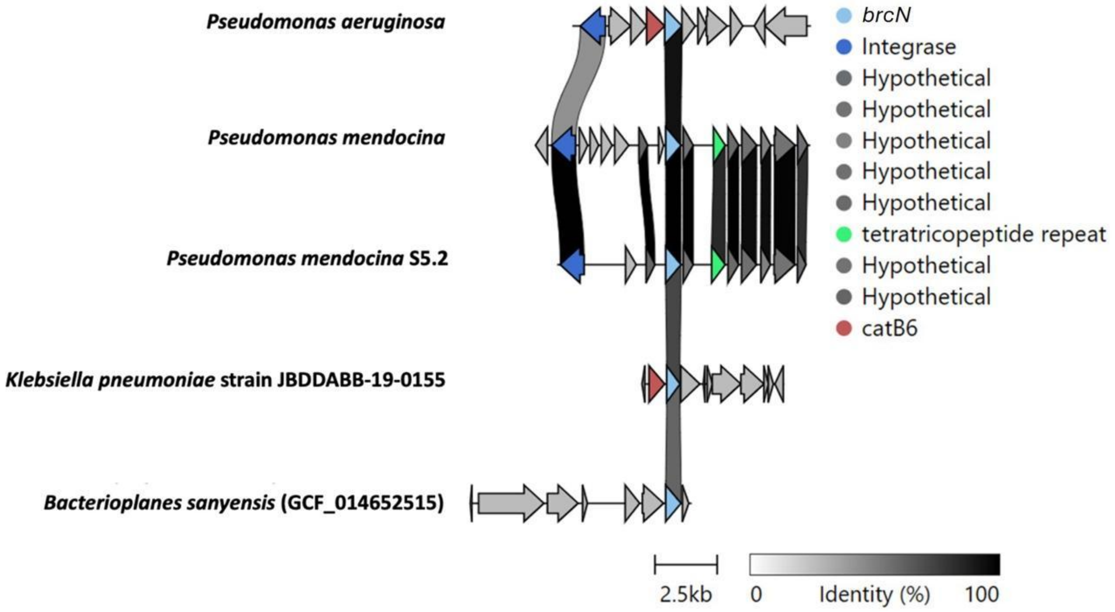


### Arconi

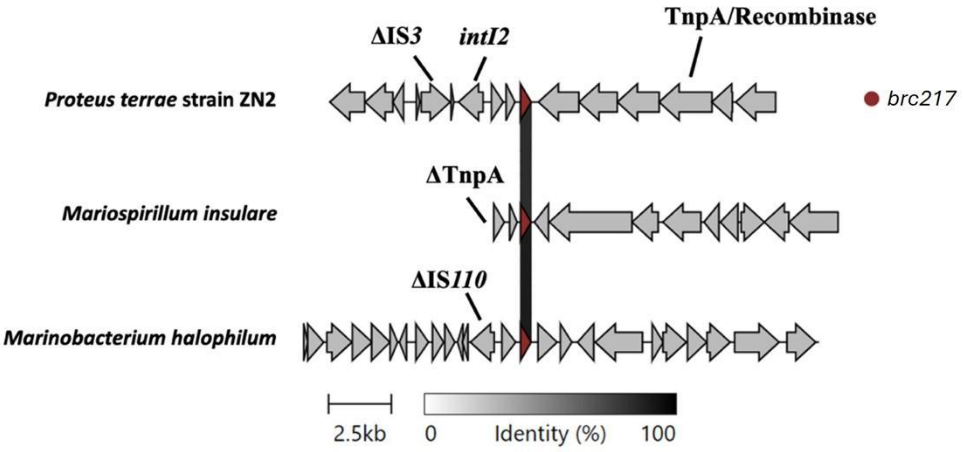

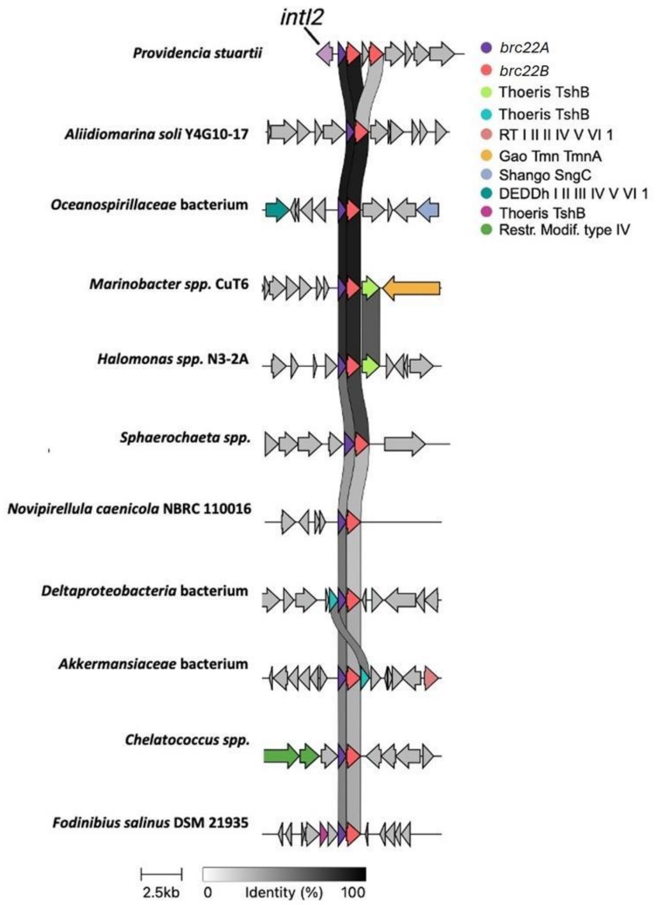


**Fig. S6. Genetic contexts of *brc*s.** Images show the genetic context of *brc* hits in the databases using CAGECAT (Comparative Gene Cluster Analysis Toolbox).

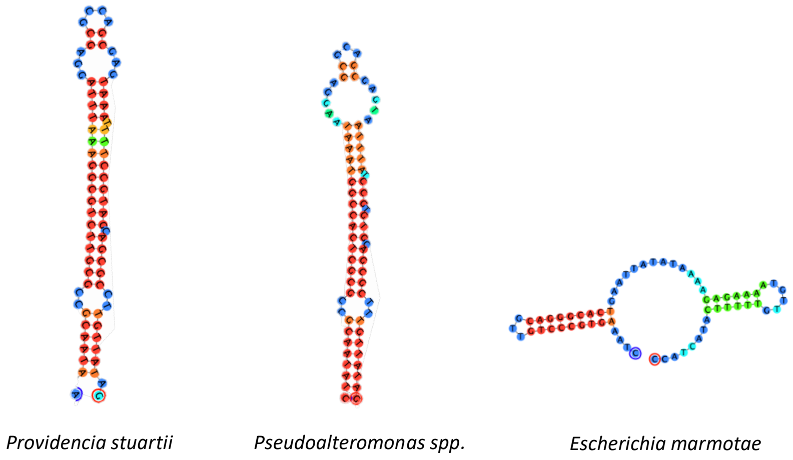


**Fig. S7. Predicted secondary structures of *attC* sites identified in Cosus homologs**. Each structure represents the predicted folding pattern for the *attC* sites associated with each Cosus homolog. The structures are color-coded based on nucleotide positions, with blue representing the outer loops and red indicating the stems of the hairpins.

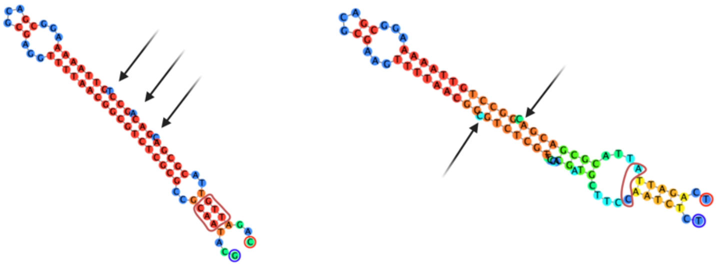

Tragantía in *brc22*

Tragantía in *Marinobacter*

**Fig. S8. Predicted secondary structures of *attC* sites identified in Tragantía homologs**. Arrows show the extrahelical bases that are recognised by the integrase as landmarks in *attC* sites. The red shapes mark the correct formation of the R box in the *attC* of *brc22*, with a perfect GTT/AAC binding between both arms of the stem. Instead, in the pseudo *attC* in *Marinobacter* there is no GTT/AAC box due to a single mutation, and one extrahelical base is found in the opposite side of the stem.

#### Table S1. Strains and plasmids used in this work.

| Bacterial Strains | | | | |
|---|---|---|---|---|
| **Number** | **Plasmids** | **Bacterial species** | **Experiments** | **Reference** |
| A249 | pMBA | *E. coli* MG1655 | General use | *Hipólito et al.* 2022 |
| B800 | None | *E. coli* IJ1862 | | *Bull et al.* 2004 |
| B931 | IncL, IncR, IncH1A | *K. pneumoniae* KP5 | | This study |
| A760 | None | *P. aeruginosa* PAO1 | | Lab collection |
| A115 | None | *E. coli* β2163 | | *Demarre et al.* 2005 |
| A123 | pSW23T::$attC_{aadA7}$ bs | *E. coli* β2163 | | *Demarre et al.* 2005 |
| A785 | pSU38Δ::*attI*/p3938 | *E. coli* TOP10 | | Lab collection |
| C246 | None | *E. coli* 594 (prophage φ80) | | *Alqurainy et al.* 2023 |
| C247 | None | *E. coli* 594 (prophage HK544) | | *Alqurainy et al.* 2023 |
| C665 | pMBA *gcu175* | *E. coli* DH5α | Screening | This study |
| C666 | pMBA *gcu93* | *E. coli* DH5α | | " |
| C667 | pMBA *gcu149* | *E. coli* DH5α | | " |
| C668 | pMBA *gcu26c* | *E. coli* DH5α | | " |
| C669 | pMBA *gcu173* | *E. coli* DH5α | | " |
| C670 | pMBA *gcu80* | *E. coli* DH5α | | " |
| C671 | pMBA *gcu153* | *E. coli* DH5α | | " |
| C672 | pMBA *gcu179* | *E. coli* DH5α | | " |
| C673 | pMBA *gcu168* | *E. coli* DH5α | | " |
| C674 | pMBA *gcu79* | *E. coli* DH5α | | " |
| C675 | pMBA *gcu18* | *E. coli* DH5α | | " |
| C676 | pMBA *gcu145* | *E. coli* DH5α | | " |
| C677 | pMBA *gcuWGS7* | *E. coli* DH5α | | " |
| C678 | pMBA *gcu32* | *E. coli* DH5α | | " |
| C679 | pMBA *gcu88* | *E. coli* DH5α | | " |
| C680 | pMBA *gcuA* | *E. coli* DH5α | | " |
| C681 | pMBA *gcu228* | *E. coli* DH5α | | " |
| C682 | pMBA *gcu170* | *E. coli* DH5α | | " |
| C683 | pMBA *gcu20* | *E. coli* DH5α | | " |
| C684 | pMBA *gcu87* | *E. coli* DH5α | | " |
| C685 | pMBA *gcu17* | *E. coli* DH5α | | " |
| C686 | pMBA *gcuWGS5* | *E. coli* DH5α | | " |
| C687 | pMBA *gcu211* | *E. coli* DH5α | | " |
| C688 | pMBA *gcu144* | *E. coli* DH5α | | " |
| C689 | pMBA *gcu28* | *E. coli* DH5α | | " |

| | | | | |
|---|---|---|---|---|
| C690 | pMBA *gcuI* | *E. coli* DH5α | | " |
| C691 | pMBA *gcuC* | *E. coli* DH5α | | " |
| C692 | pMBA *gcu19* | *E. coli* DH5α | | " |
| C693 | pMBA *gcu12* | *E. coli* DH5α | | " |
| C694 | pMBA *gcu225* | *E. coli* DH5α | | " |
| C695 | pMBA *gcuWGS10* | *E. coli* DH5α | | " |
| C696 | pMBA *gcuWGS15* | *E. coli* DH5α | | " |
| C697 | pMBA *gcu72* | *E. coli* DH5α | | " |
| C698 | pMBA *gcu189* | *E. coli* DH5α | | " |
| C699 | pMBA *gcu100* | *E. coli* DH5α | | " |
| C700 | pMBA *gcuJ* | *E. coli* DH5α | | " |
| C701 | pMBA *gcu1* | *E. coli* DH5α | | " |
| C702 | pMBA *gcu107* | *E. coli* DH5α | | " |
| C703 | pMBA *gcu76c* | *E. coli* DH5α | | " |
| C704 | pMBA *gcuWGS2* | *E. coli* DH5α | | " |
| C705 | pMBA *gcu76* | *E. coli* DH5α | | " |
| C706 | pMBA *gcuP* | *E. coli* DH5α | | " |
| C707 | pMBA *gcu78* | *E. coli* DH5α | | " |
| C708 | pMBA *gcu7* | *E. coli* DH5α | | " |
| C709 | pMBA *gcu57* | *E. coli* DH5α | | " |
| C710 | pMBA *gcu232* | *E. coli* DH5α | | " |
| C711 | pMBA *gcu221* | *E. coli* DH5α | | " |
| D153 | pMBA *gcu23* | *E. coli* DH5α | Screening | " |
| C713 | pMBA *gcu114* | *E. coli* DH5α | | " |
| C714 | pMBA *gcuQ* | *E. coli* DH5α | | " |
| C715 | pMBA *gcu11* | *E. coli* DH5α | | " |
| C716 | pMBA *gcu89* | *E. coli* DH5α | | " |
| C717 | pMBA *gcu92* | *E. coli* DH5α | | " |
| C718 | pMBA *gcu96* | *E. coli* DH5α | | " |
| C720 | pMBA *gcuF* | *E. coli* DH5α | | " |
| C721 | pMBA *gcu199* | *E. coli* DH5α | | " |
| C722 | pMBA *gcu65* | *E. coli* DH5α | | " |
| C723 | pMBA *gcu217* | *E. coli* DH5α | | " |
| C724 | pMBA *gcu238* | *E. coli* DH5α | | " |
| C725 | pMBA *gcu236* | *E. coli* DH5α | | " |
| C726 | pMBA *gcuWGS1* | *E. coli* DH5α | | " |
| D148 | pMBA *gcu14* | *E. coli* DH5α | | " |
| C728 | pMBA *gcu177* | *E. coli* DH5α | | " |
| C729 | pMBA *gcu227* | *E. coli* DH5α | | " |
| C730 | pMBA *gcuWGS21* | *E. coli* DH5α | | " |
| C731 | pMBA *gcu185* | *E. coli* DH5α | | " |
| C732 | pMBA *gcu231* | *E. coli* DH5α | | " |
| C733 | pMBA *gcu46* | *E. coli* DH5α | | " |
| C734 | pMBA *gcu9* | *E. coli* DH5α | | " |
| C735 | pMBA *gcu83* | *E. coli* DH5α | | " |

| | | | | |
|---|---|---|---|---|
| C736 | pMBA *gcu48* | *E. coli* DH5α | Screening | " |
| C737 | pMBA *gcu113* | *E. coli* DH5α | | " |
| C738 | pMBA *gcu197* | *E. coli* DH5α | | " |
| C739 | pMBA *gcuWGS3* | *E. coli* DH5α | | " |
| C740 | pMBA *gcu262* | *E. coli* DH5α | | " |
| C741 | pMBA *gcu184* | *E. coli* DH5α | | " |
| C742 | pMBA *gcu137* | *E. coli* DH5α | | " |
| C743 | pMBA *gcu143* | *E. coli* DH5α | | " |
| C744 | pMBA *gcu141* | *E. coli* DH5α | | " |
| C745 | pMBA *gcu142* | *E. coli* DH5α | | " |
| C746 | pMBA *gcu121* | *E. coli* DH5α | | " |
| C747 | pMBA *gcu24* | *E. coli* DH5α | | " |
| C748 | pMBA *gcu8* | *E. coli* DH5α | | " |
| C749 | pMBA *gcu109* | *E. coli* DH5α | | " |
| C750 | pMBA *gcu101* | *E. coli* DH5α | | " |
| C751 | pMBA *gcu21* | *E. coli* DH5α | | " |
| C752 | pMBA *gcu77* | *E. coli* DH5α | | " |
| C753 | pMBA *gcu230* | *E. coli* DH5α | | " |
| C754 | pMBA *gcuH* | *E. coli* DH5α | | " |
| C755 | pMBA *gcu163* | *E. coli* DH5α | | " |
| C756 | pMBA *gcu223* | *E. coli* DH5α | | " |
| C757 | pMBA *gcu29* | *E. coli* DH5α | | " |
| C758 | pMBA *gcu190* | *E. coli* DH5α | | " |
| C759 | pMBA *gcu172* | *E. coli* DH5α | | " |
| C760 | pMBA *gcu16* | *E. coli* DH5α | | " |
| C761 | pMBA *gcu108* | *E. coli* DH5α | | " |
| C762 | pMBA *gcu196* | *E. coli* DH5α | | " |
| C763 | pMBA *gcu5b* | *E. coli* DH5α | | " |
| C764 | pMBA *gcu37* | *E. coli* DH5α | | " |
| C765 | pMBA *gcu135* | *E. coli* DH5α | | " |
| C766 | pMBA *gcuWGS6* | *E. coli* DH5α | | " |
| C767 | pMBA *gcu253A* | *E. coli* DH5α | | " |
| C768 | pMBA *gcu133* | *E. coli* DH5α | | " |
| C769 | pMBA *gcu158* | *E. coli* DH5α | | " |
| C770 | pMBA *gcu39* | *E. coli* DH5α | | " |
| C771 | pMBA *gcu74* | *E. coli* DH5α | | " |
| C149 | pMBA *gcu157* | *E. coli* DH5α | | " |
| C773 | pMBA *gcu122* | *E. coli* DH5α | | " |
| C774 | pMBA *gcu128* | *E. coli* DH5α | | " |
| C775 | pMBA *gcu59* | *E. coli* DH5α | | " |
| C776 | pMBA *gcu154A* | *E. coli* DH5α | | " |
| D150 | pMBA *gcu138* | *E. coli* DH5α | | " |
| C778 | pMBA *gcu118* | *E. coli* DH5α | | " |
| C779 | pMBA *gcu68* | *E. coli* DH5α | | " |
| C780 | pMBA *gcu52* | *E. coli* DH5α | | " |

| | | | | |
|---|---|---|---|---|
| C782 | pMBA *gcuWGS13* | *E. coli* DH5α | Screening | “ |
| C784 | pMBA *gcu182B* | *E. coli* DH5α | | “ |
| C785 | pMBA *gcu233* | *E. coli* DH5α | | “ |
| C786 | pMBA *gcu22A* | *E. coli* DH5α | | “ |
| C788 | pMBA *gcu41* | *E. coli* DH5α | | “ |
| C789 | pMBA *gcu43* | *E. coli* DH5α | | “ |
| C790 | pMBA *gcu26b* | *E. coli* DH5α | | “ |
| C792 | pMBA *gcu6* | *E. coli* DH5α | | “ |
| C793 | pMBA *gcu111* | *E. coli* DH5α | | “ |
| C795 | pMBA *gcu102* | *E. coli* DH5α | | “ |
| C796 | pMBA *gcu54* | *E. coli* DH5α | | “ |
| C797 | pMBA *gcu117* | *E. coli* DH5α | | “ |
| C798 | pMBA *gcu119* | *E. coli* DH5α | | “ |
| C799 | pMBA *gcu136* | *E. coli* DH5α | | “ |
| D383 | pMBA-*brcI* | *E. coli* IJ1862 | Positive BRiCs in the screening against the complete panel of phages | “ |
| D398 | pMBA-*brcWGS10* | *E. coli* IJ1862 | | “ |
| D400 | pMBA-*brc72* | *E. coli* IJ1862 | | “ |
| D403 | pMBA-*brc1* | *E. coli* IJ1862 | | “ |
| D410 | pMBA-*brc7* | *E. coli* IJ1862 | | “ |
| D478 | pMBA-*brc16* | *E. coli* IJ1862 | | “ |
| D475 | pMBA-*brc29* | *E. coli* IJ1862 | | “ |
| D414 | pMBA-*brc32* | *E. coli* IJ1862 | | “ |
| D493 | pMBA-*brc68* | *E. coli* IJ1862 | | “ |
| D441 | pMBA-*brc88* | *E. coli* IJ1862 | | “ |
| D418 | pMBA-*brc89* | *E. coli* IJ1862 | | “ |
| D419 | pMBA-*brc92* | *E. coli* IJ1862 | | “ |
| D415 | pMBA-*brc114* | *E. coli* IJ1862 | | “ |
| D508 | pMBA-*brc119* | *E. coli* IJ1862 | | “ |
| D517 | pMBA-*brc126* | *E. coli* IJ1862 | | “ |
| D431 | pMBA-*brc149* | *E. coli* IJ1862 | | “ |
| D473 | pMBA-*brc163* | *E. coli* IJ1862 | | “ |
| D497 | pMBA-*brc182* | *E. coli* IJ1862 | | “ |
| D401 | pMBA-*brc189* | *E. coli* IJ1862 | | “ |
| D422 | pMBA-*brc199* | *E. coli* IJ1862 | | “ |
| D474 | pMBA-*brc223* | *E. coli* IJ1862 | | “ |
| D443 | pMBA-*brc228* | *E. coli* IJ1862 | | “ |
| D412 | pMBA-*brc232* | *E. coli* IJ1862 | | “ |
| D425 | pMBA-*brc236* | *E. coli* IJ1862 | | “ |
| D424 | pMBA-*brc238* | *E. coli* IJ1862 | | “ |
| D442 | pMBA-*brcA* | *E. coli* IJ1862 | | “ |
| D421 | pMBA-*brcF* | *E. coli* IJ1862 | | “ |
| D402 | pMBA-*brcJ* | *E. coli* IJ1862 | | “ |
| D408 | pMBA-*brcP* | *E. coli* IJ1862 | | “ |
| D416 | pMBA-*brcQ* | *E. coli* IJ1862 | | “ |
| D426 | pMBA-*brcWGS1* | *E. coli* IJ1862 | | “ |

| | | | | |
|---|---|---|---|---|
| D406 | pMBA-*brcWGS2* | *E. coli* IJ1862 | Positive BRiCs in the screening against the complete panel of phages | “ |
| D529 | pMBA-*brc24*-T12A | *E. coli* DH5α | | “ |
| D534 | pMBA-*brc24*-D96A | *E. coli* DH5α | | “ |
| D538 | pMBA-*brc24*-D14A | *E. coli* DH5α | | “ |
| C809 | pMBA | *E. coli* IJ1862 | High/low MOI Phage panel | “ |
| C800 | pMBA *brc59* | *E. coli* IJ1862 | | “ |
| C801 | pMBA *brc128* | *E. coli* IJ1862 | | “ |
| C802 | pMBA *brc135* | *E. coli* IJ1862 | | “ |
| C803 | pMBA *brc167* | *E. coli* IJ1862 | | “ |
| C804 | pMBA *brc167.2* | *E. coli* IJ1862 | | “ |
| C805 | pMBA *brc142* | *E. coli* IJ1862 | | “ |
| C806 | pMBA *brc24* | *E. coli* IJ1862 | | “ |
| C807 | pMBA *brc233* | *E. coli* IJ1862 | | “ |
| C808 | pMBA *brcN* | *E. coli* IJ1862 | | “ |
| C933 | pMBA *brc76* | *E. coli* IJ1862 | | “ |
| C934 | pMBA *brc217* | *E. coli* IJ1862 | | “ |
| C935 | pMBA *brcWGS21* | *E. coli* IJ1862 | | “ |
| C936 | pMBA *brc113* | *E. coli* IJ1862 | | “ |
| C819 | pMBA | *K. pneumoniae* KP5 | BRiCs in other species | “ |
| C810 | pMBA *brc59* | *K. pneumoniae* KP5 | | “ |
| C811 | pMBA *brc128* | *K. pneumoniae* KP5 | | “ |
| C812 | pMBA *brc135* | *K. pneumoniae* KP5 | | “ |
| C813 | pMBA *brc167* | *K. pneumoniae* KP5 | | “ |
| C814 | pMBA *brc167.2* | *K. pneumoniae* KP5 | | “ |
| C815 | pMBA *brc142* | *K. pneumoniae* KP5 | | “ |
| C816 | pMBA *brc24* | *K. pneumoniae* KP5 | | “ |
| C817 | pMBA *brc233* | *K. pneumoniae* KP5 | | “ |
| C818 | pMBA *brcN* | *K. pneumoniae* KP5 | | “ |
| C939 | pMBA *brc76* | *K. pneumoniae* KP5 | | “ |
| C940 | pMBA *brc217* | *K. pneumoniae* KP5 | | “ |
| C941 | pMBA *brcWGS21* | *K. pneumoniae* KP5 | | “ |
| C942 | pMBA *brc113* | *K. pneumoniae* KP5 | | “ |
| C971 | pBTZ PcW *brc24* | *P. aeruginosa* PAO1 | | “ |
| C972 | pBTZ PcW *brc142* | *P. aeruginosa* PAO1 | | “ |

| | | | | |
|---|---|---|---|---|
| C973 | pBTZ PcW *brcWGS21* | *P. aeruginosa* PAO1 | BRiCs in other species | “ |
| C974 | pBTZ PcW *brc113* | *P. aeruginosa* PAO1 | | “ |
| C975 | pBTZ-PcW | *P. aeruginosa* PAO1 | | “ |
| C898 | pBTZ PcS *brcN* | *P. aeruginosa* PAO1 | | “ |
| C915 | pMBA *bla*$_{\text{OXA-10}}$ | *E. coli* IJ1862 | Multicassette arrays | “ |
| C916 | pMBA *brc24-bla*$_{\text{OXA-10}}$ | *E. coli* IJ1862 | | “ |
| D159 | pMBA *brc24-brc167.2* | *E. coli* IJ1862 | | “ |
| D379 | pMBA *aacA54-aacA8-brc24* | *E. coli* DH5α | | “ |
| D022 | pMBA-ΔGFP | *E. coli* IJ1862 | Fitness cost measurement | “ |
| C962 | pMBA-ΔGFP *brc76* | *E. coli* IJ1862 | | “ |
| D063 | pMBA-ΔGFP *brc113* | *E. coli* IJ1862 | | “ |
| D068 | pMBA-ΔGFP *brc217* | *E. coli* IJ1862 | | “ |
| D023 | pMBA-ΔGFP *brc76* | *E. coli* IJ1862 | | “ |
| D024 | pMBA-ΔGFP *brc128* | *E. coli* IJ1862 | | “ |
| D025 | pMBA-ΔGFP *brc135* | *E. coli* IJ1862 | | “ |
| D026 | pMBA-ΔGFP *brc167* | *E. coli* IJ1862 | | “ |
| D027 | pMBA-ΔGFP *brc142* | *E. coli* IJ1862 | | “ |
| D028 | pMBA-ΔGFP *brc24* | *E. coli* IJ1862 | | “ |
| D029 | pMBA-ΔGFP *brcN* | *E. coli* IJ1862 | | “ |
| D064 | pMBA-ΔGFP *brc167.2* | *E. coli* IJ1862 | | “ |
| D065 | pMBA-ΔGFP *brcWGS21* | *E. coli* IJ1862 | | “ |
| D066 | pMBA-ΔGFP *brc59* | *E. coli* IJ1862 | | “ |
| D067 | pMBA-ΔGFP *brc233* | *E. coli* IJ1862 | | “ |
| D068 | pMBA-ΔGFP *brc217* | *E. coli* IJ1862 | | “ |
| D063 | pMBA-ΔGFP *brc113* | *E. coli* IJ1862 | | “ |
| D092 | pMBA-ΔGFP *brcN* | *K. pneumoniae* KP5 | | “ |
| D093 | pMBA-ΔGFP *brc59* | *K. pneumoniae* KP5 | | “ |
| D094 | pMBA-ΔGFP *brc128* | *K. pneumoniae* KP5 | | “ |
| D095 | pMBA-ΔGFP *brcWGS21* | *K. pneumoniae* KP5 | | “ |
| D096 | pMBA-ΔGFP *brc24* | *K. pneumoniae* KP5 | | “ |
| D097 | pMBA-ΔGFP *brc76* | *K. pneumoniae* KP5 | | “ |
| D098 | pMBA-ΔGFP *brc142* | *K. pneumoniae* KP5 | | “ |
| D099 | pMBA-ΔGFP *brc113* | *K. pneumoniae* KP5 | | “ |
| D100 | pMBA-ΔGFP | *K. pneumoniae* KP5 | | “ |
| D101 | pMBA-ΔGFP *brc167.2* | *K. pneumoniae* KP5 | | “ |

| | | | | |
|---|---|---|---|---|
| D102 | pMBA-ΔGFP *brc167* | *K. pneumoniae* KP5 | Fitness cost measurement | " |
| D103 | pMBA-ΔGFP *brc217* | *K. pneumoniae* KP5 | | " |
| D104 | pMBA-ΔGFP *brc135* | *K. pneumoniae* KP5 | | " |
| D105 | pMBA-ΔGFP *brc233* | *K. pneumoniae* KP5 | | " |
| D002 | pMBA | *E. coli* 594 (prophage φ80) | Prophage induction | " |
| D286 | pMBA *brc59* | *E. coli* 594 (prophage φ80) | | " |
| D016 | pMBA *brc128* | *E. coli* 594 (prophage φ80) | | " |
| C998 | pMBA *brc135* | *E. coli* 594 (prophage φ80) | | " |
| D018 | pMBA *brc167* | *E. coli* 594 (prophage φ80) | | " |
| D011 | pMBA *brc167.2* | *E. coli* 594 (prophage φ80) | | " |
| D001 | pMBA *brc142* | *E. coli* 594 (prophage φ80) | | " |
| D010 | pMBA *brc24* | *E. coli* 594 (prophage φ80) | | " |
| C999 | pMBA *brc233* | *E. coli* 594 (prophage φ80) | | " |
| D021 | pMBA *brcN* | *E. coli* 594 (prophage φ80) | | " |
| C957 | pMBA *brc76* | *E. coli* 594 (prophage φ80) | | " |
| C951 | pMBA *brc217* | *E. coli* 594 (prophage φ80) | | " |
| C948 | pMBA *brcWGS21* | *E. coli* 594 (prophage φ80) | | " |
| C954 | pMBA *brc113* | *E. coli* 594 (prophage φ80) | | " |
| C996 | pMBA | *E. coli* 594 (prophage HK544) | | " |
| D285 | pMBA *brc59* | *E. coli* 594 (prophage HK544) | | " |
| D008 | pMBA *brc128* | *E. coli* 594 (prophage HK544) | | " |
| C995 | pMBA *brc135* | *E. coli* 594 (prophage HK544) | | " |
| D017 | pMBA *brc167* | *E. coli* 594 (prophage HK544) | | " |
| D019 | pMBA *brc167.2* | *E. coli* 594 (prophage HK544) | | " |
| C997 | pMBA *brc142* | *E. coli* 594 (prophage HK544) | | " |
| D003 | pMBA *brc24* | *E. coli* 594 (prophage HK544) | | " |
| C994 | pMBA *brc233* | *E. coli* 594 (prophage HK544) | | " |

| | | | | |
|---|---|---|---|---|
| D020 | pMBA *brcN* | *E. coli* 594 (prophage HK544) | Prophage induction | " |
| C959 | pMBA *brc76* | *E. coli* 594 (prophage HK544) | | " |
| C953 | pMBA *brc217* | *E. coli* 594 (prophage HK544) | | " |
| C950 | pMBA *brcWGS21* | *E. coli* 594 (prophage HK544) | | " |
| C956 | pMBA *brc113* | *E. coli* 594 (prophage HK544) | | " |
| B918 | pSW23T:: *brc59* bs | *E. coli* β2163 | Cassette recombination assays | " |
| B920 | pSW23T::*brc128* bs | *E. coli* β2163 | | " |
| B924 | pSW23T::*brc135* bs | *E. coli* β2163 | | " |
| D291 | pSW23T::*brcN* bs | *E. coli* β2163 | | " |
| C984 | pSW23T::*brc24* bs | *E. coli* β2163 | | " |
| C988 | pSW23T::*brc76* bs | *E. coli* β2163 | | " |
| C986 | pSW23T::*brc113* bs | *E. coli* β2163 | | " |
| D292 | pSW23T::*brc142* bs | *E. coli* β2163 | | " |
| B922 | pSW23T::*brc167* bs | *E. coli* β2163 | | " |
| D293 | pSW23T::*brc167.2* bs | *E. coli* β2163 | | " |
| C985 | pSW23T::*brc217* bs | *E. coli* β2163 | | " |
| D294 | pSW23T::*brc233* bs | *E. coli* β2163 | | " |
| C987 | pSW23T::*brcWGS21* bs | *E. coli* β2163 | | " |
| B917 | pSW23T:: *brc59* ts | *E. coli* β2163 | | " |
| B919 | pSW23T::*brc128* ts | *E. coli* β2163 | | " |
| B923 | pSW23T::*brc135* ts | *E. coli* β2163 | | " |
| D295 | pSW23T::*brcN* ts | *E. coli* β2163 | | " |
| C989 | pSW23T::*brc24* ts | *E. coli* β2163 | | " |
| C993 | pSW23T::*brc76* ts | *E. coli* β2163 | | " |
| C991 | pSW23T::*brc113* ts | *E. coli* β2163 | | " |
| D296 | pSW23T::*brc142* ts | *E. coli* β2163 | | " |
| B921 | pSW23T::*brc167* ts | *E. coli* β2163 | | " |
| D297 | pSW23T::*brc167.2* ts | *E. coli* β2163 | | " |
| C990 | pSW23T::*brc217* ts | *E. coli* β2163 | | " |
| D298 | pSW23T::*brc233* ts | *E. coli* β2163 | | " |
| C992 | pSW23T::*brcWGS21* ts | *E. coli* β2163 | | " |
| D278 | pMBA PcW *brc22* | *E. coli* IJ1862 | Naturally occurring defense islands | " |
| D279 | pMBA PcW *brc23* | *E. coli* IJ1862 | | " |
| D283 | pMBA PcW *brc22* | *K. pneumoniae* KP5 | | " |
| D284 | pMBA PcW *brc23* | *K. pneumoniae* KP5 | | " |
| D287 | pMBA PcW *brc23-brc24* | *E. coli* DH5α | | " |
| D289 | pMBA PcW *brc23-brc24* | *E. coli* IJ1862 | | " |

**Table S2. Oligonucleotides Used for Cloning and Amplification**.

| Name | Sequence 5'-> 3' | Role |
|---|---|---|
| gBLOCK F | GCAGTCGCCCTAAAACAAAG | Amplifying *gcu*s for cloning in pMBA |
| gBLOCK R | GCTGCAGCGTCGACGCCTAA | |
| GFP F BackBone pMBA | TTAGGCGTCGACGCTGCA | Amplify pMBA backbone |
| Int R BackBone pMBA | CTTTGTTTTAGGGCGACTGC | |
| Array-Ø F | GCAGTCGCCCTAAAACAAAGTTA GGCCGCACAAAATCAACG | Amplify *aacA54* and *aacA8* to clone it into pMBA KO |
| Array-Ø R | GCTGCAGCGTCGACGCCTAACGT TTGACATGAGGGGCGGC | |
| *brc24*-ArrayØ F | GCCGCCCCTCATGTCAAACGTTA TAGCTCAAGCAGCTCATCTCG | Amplify *brc24* to clone it in pArray Ø |
| *brc24*-ArrayØ R | GCTGCAGCGTCGACGCCTAACGC CGCGTTCTGCGGAAATT | |
| BackBone pArray *brc24* R | CGTTTGACATGAGGGGCGGC | Used with GFP_F_BackBone_pMBA on pArrayØ to amplify the backbone to clone *brc24* |
| BB pSW23T GA-R | ACTTTGTTTTAGGGCGACTGCGAT ATCAAGCTTATCGATAC | Amplify pSW23T backbone to clone *gcu*s amplified with gBLOCK F and gBLOCK R, to deliver the bottom strand |
| BB pSW23T GA-F | TTAGGCGTCGACGCTGCAGCACT AGTTCTAGAGCGGCCGC | |
| BB pSW23T AG-R | TTAGGCGTCGACGCTGCAGCGAT ATCAAGCTTATCGATAC | Amplify pSW23T backbone to clone *gcu*s amplified with gBLOCK F and gBLOCK R, to deliver the top strand |
| BB pSW23T AG-F | ACTTTGTTTTAGGGCGACTGCACT AGTTCTAGAGCGGCCGC | |
| *brc128* pBAD tail F | TTAACCATGGATCCGAGCTCATG GTAAATAGTAATTCTGGTGTAGA GG | Clone *brc128AB* into pBAD |
| brc128 pBAD tail R | TGAGATGAGTTTTTGTTCTAGAG GATATTGAAATATTAAGTTAG | |
| pBAD BBExt F | TCTAGAACAAAAACTCATCTC | Amplify pBAD backbone |
| pBAD BBExt R | GAGCTCGGATCCATGGTTAA | |
| *gcu24*-BB 167-2 | CGCCGCGTTCTGCGGAAATTT | Backbone amplification to create pArray1 (*brc24-brc167.2*). Use with GFP_F_BackBone_pMBA |
| 167-2 for p24 F | AATTTCCGCAGAACGCGGCGTTA GGCTTTAAAACCATCACTTCCG | Amplify *brc167.2* to clone in second position to create pArray-1 |
| 167-2 for p24 R | GCTGCAGCGTCGACGCCTAACTT CGTTTTATGCGGTGGCA | |
| DdmABC pBAD F | TTAACCATGGATCCGAGCTCGAG AATCGAGTTGCTATTGGT | Amplify *ddmABC* for cloning in pBAD |
| DdmABC pBAD R | TGAGATGAGTTTTTGTTCTAGAGC ACAACTTGTAAGATAGCCTTGC | |
| *gcu24* for 22/23 F | AATTTGCCGCAGAGCGCGGCGTT ATAGCTCAAGCAGCTCATCTC | Amplify *brc24* to clone it in pMBA::*brc23* |
| *gcu22/23* for 23/24 R | TAACGCCGCGCTCTGCG | |
| D6A-ext-5 | AAATAGAATTAGCTTTTGGCGCG ATAACCATAGATAACGC | Used to mutate *brc24* |
| D6A-ext-3 | GCCAAAAGCTAATTCTATTTCGTC CATAAAACCTTCACGTTTC | " |
| T12A-ext-5 | GATTTTGGCGCGATAGCCATAGA TAACGCGACCTATAAGG | " |
| T12A-ext-3 | ATGGCTATCGCGCCAAAATCTAA TTCTATTTCGTCCATAAAAC | " |

| | | |
|---|---|---|
| D14A-ext-5 | GCGTTAGCTATGGTTATATCGCG CCAAAATCTAATTCTATTTCG | “ |
| D14A-ext-3 | GCGATAACCATAGCTAACGCGAC CTATAAGGGCGAAGGTTATAG | “ |
| E42A-ext-5 | GGGCTTGCTTTAAATTGGCGCAT CTGAGCAAGTAAGCCCCTC | “ |
| E42A-ext-3 | CGCCAATTTAAAGCAAGCCCAGT TAAAGTTCTTCAAACCG | “ |
| D47A-ext-5 | GTGCACTATAGCGGTTTGAAGAA CTTTAACTGGGCTTTCTTTAAATT GG | “ |
| D47A-ext-3 | CTTCAAACCGCTATAGTGCACAA TGAAGCGATCAATCATATC | “ |
| S67A-ext-5 | CTGCTCAATCGCGGATCGGGTTTT TGATACTCTTGCCCG | “ |
| S67A-ext-3 | CCCGATCCGCGATTGAGCAGGCA CTAAGGTCAGAAATAAAC | “ |
| E69A-ext-5 | CTGCGCAATCGAGGATCGGGTTT TTGATATCTCTTGCCCG | “ |
| E69A-ext-3 | CCCGATCCTCGATTGCGCAGGCA CTAAGGTCAGCAAATAAAC | “ |
| D96A-ext-5 | CTTCGCTGCCAGCAACTGATAGC AACGCTCGAGCATTTTC | “ |
| D96A-ext-3 | ATCAGTTGCTGGCAGCGAAGCTG AAATAGCGGAAGCGAGG | “ |
| D120A-ext-5 | CTTCCAGAATCGAGCTTCTCAGCT CCAATAAATTCATAATATTTTTC | “ |
| D120A-ext-3 | GAGAAGCTCGATTCTGGAAGATA TGCAGATCTATCTCGTC | “ |
| D146A-ext-5 | TTTTTGGCCTTTCCGGATTCAAAA GGAGCTTCTGTAGCAAAATAC | “ |
| D146A-ext-3 | GAATCCGGAAAGGCCAAAAAAA ATGAATTTCCAGATGCGA | “ |
| D153A-ext-5 | AATAGTGCAATCGCAGCTGGAAA TTCATTTTTTTGTCCTTTCCGG | “ |
| D153A-ext-3 | CCAGCTGCGATTGCACTATTAGC CCTTGAAGGTTGGGCTG | “ |
| D177A-ext-5 | CCTTAGCTTGACTTACGGCGATA ATATTAATTTCATTTTCTTCAGCC CAAC | “ |
| D177A-ext-3 | CGCCGTAAGTCAAGCTAAGGGCT GGAAGAACTTTTCTGAAG | “ |

**Table S3. List of the *gcu* sequences with bacteriophage defense activity.** In green are the ORFs. When there is an overlap between ORFs the sequence is underlined.

*gcu1*

TTAGGCTTGTTGGAATACACCAGTATTCATCGGCCTTGTGGGGAGCCTGTCCTGCGGCAGTCGTCCGCACCACCGAGCAGCTCGGGCTCAGTG
CTGGCAGGGCTTCATCGCAGGTTGAAGCTGGGTTAGATTGTGGTCATCGTCAGCGCATGGCTGACGCCATACGCAAAGGGAGGCGTGATGGC
AAGGTTCGGTTGGGCAACCACCACAGCACATCGCACACACCAGGCTGTCGCCTCGTGTGCGGGCTCGCTGTCCGCGCGACCACCGGCTTACTT
CCACCAGCCTAACTGGTCGGTCAAGCGGACGCCAACACAGGCCATGCCTTCGGCATTCTCATGGCCTGTGTCGGTGCCCTACGCCCTGTCGGG
CTCCGGCGCCGCTTACCTTGGGCG

*gcu7*

TTAGGCTGCAGACCAAACAATGCCGTATTTCAAAGTCATTCTTTCCGGGCGTGGCATTGATCTGCCGTTTGATGGCGATTCGGCCATTGGTTTC
TTCACCACCAGGCTTGTCCGTTCTATGGACCCAGCAAGCGCTGAATCCTTGGCCAAAGGCCTAGTTCAGGCAGAGTGGCTCCCCGGCGGCACC
TATGCAACGGTTAATCGCGGCTCGCAGCCCACCTTGTCTGTCGAGCGGTCGTCCCCCGTCGGCTGGCTGACGGGCACTTTCGGTCGCAAACCA
TCCGGCTACTCATTCTACCGCCATGAGGACTAGACTGTGGCCTAACAATTCATTCAAGCCGACGCCACTTCGCGGCGCGGCTTACTTCAGGCG

*gcu16*

TTATGAGTATCTGTGCCCTGAGAGTATGAAAGCAGAATGGAAGCTCTATTCGTAGTATGCATTTATTTGGTGGTAGCCGGCATAGCCATTCTTC
CAGCCGTATTGGTTTGGAGCTTAGCTGGCCTATGGCTCCCCGCTTGGTTTTCACTACCTCCTATTCGTATGGCTTTAATATGTTTGGCATTATCC
ATTGTTACGGCATTGCTATTTAGTCTTGAACTTCCGGGCGGTATGCTTTCAGTTGGGCCATCTATCGTAGTTCTGTCATTGACCTGGTCTTTACT
TCTACTTCCCGTACTTTGTCTAGTTAGGTATTACCGCTATGCCACAACAAAACAGTCATAACAATCTGTTTCAGTACATTCCGGCACCTTTGGT
GCCTCCATCGGACAGCCTTTACAGCGTCGCGCCTCCAAGGCTGCCGCTGAACAAGGGCG

*gcu22*

TTAGACCTTCAGGAGGTGATAGGTGAGTTACAGAAATAAAACGTACGTCATCTTTGATGGCGATGAAGACATGTGGGCATACCGATACATGC
GTGGCTGGAAAGCTAACGAAAATATTGATTTCAACTTCTTTGATGCTCATGATTTGAAGCCGCTTACAGATCGTGCAGGAGAAGATACGGTTA
AGAGGCGGCTGAGGGAAAGGCTTGGGGATACCAAGCAGGCCATAGTTCTTATTGGCGAAAAAACAAAAAATCTGTATCGATTTGTGCGCTGG
GAACTTGAAACATGCATGAACCTGGATATACCAATCATAGCTGTTAATTTAAATGGACAAAGAAGCCAGGACGAGAATCTTTGTCCTCCAATA
ATTCGTGATGAGTATGTAGTCCACATCCCGTTCAAGCTAAAAATAATTCAGTATGCCTTGGACAACTTTCCAGGAGAGTTCCATCGGCGCAAT
CTAAGCGATAAAGGACCAAGGCTATACAACGACTCCGTCTACAAGCAGCTGGGGATCGAATGAAAAACGACGAATACCCTGCGCTGTACTGC
GCCACCAACAAGGCATCAATGGATGCCCAAAACCAGTACTTAAAGTTCGTTAAAGTTTATTCTCTTCTTCTTATTGCTGCGGCAGGGTTGGGCG
TTTACGGAATAAATGAAAAAGCATCTGCGGTCGCAGCCGCCTTGCTCGTTATAGGATCTATATTCCTATCTGTAATGATGCTGCTGAGACGAG
ATGAAGATACATGGTATCGGGCACGCTCTGTCGCCGAATCCGTTAAAACAAGCTCTTGGCGTTTCATGATGAGGTGCGAGCCTTACGTTGACG
CCCCTGATGTAAGAGTAGTGAAGTCAAATTTCAGGACACGACTGAAAAGTATCTTGAGTGAGCATAAGGACTTGGCGGAGCATCTAGGAGGT
TCGGTGTCTGAACAAGAGCAGATTACCGACAAGATGTGCGAAGTAAGAAATCTATCGTGGGAGCAGAGGGCTGACTTTTATCGCACGCATAG
AATTGACGAGCAACGTTCTTGGTATGCAAACAAGTCTGCTTGGAACCGCAAAAAAGGAAGGCTGTGGTTTGCTGTTTTAATCGGCTGCCAAGC
TTTTGCCGTTTTGTTCCTAATTTTGCGTGTTGCTTATCCGGATTGGGGATATTGGCCTGCGGATGTATTCGTTGTTGCGGCTGGATCGGCTTTAA
CCTGGATACAGGTGAAACGGTTTAAGGAGCTAGCAGCTGCATATGGATTGACAGCGCACGAGATTGGAGTTGTGAGAGGTGAGCTAGAGCAA
ATCGACTCTGGAGAAAAGTTGGCGCAGTTTGTAGCTGATAGCGAGAACGCATTTTCTAGAGAGCACACTCAATGGCTAGCTCGCAAAGACTCT
ATATAGGGTCTAACAATGCGCTGCTGTCGGACAATTTTTCCGCTGCGCTCCAAAATTGCCGCAGAGCGCGGCG

*gcu23*

TTATGCAGTCCAATGTCTGAATTAAGGAACTAAAAATGAGCACACGCCAAATCCATGTATTCATCAGTCATGCATGGAAATATTCTGGCCATT
ATCAAACGCTTGCTGATTGGATTTTTAATCAGAATTGGAGCGTGGGGCAAGCATCTCTGGATTTTAGAAATTATTCGGTTCCTAAAGATGATCC
AATTCATGATGCGCCAAATGATAGGCAGCTTAGAGATGCGATTATGAGGCAGATATCCATGAGTCATGTGGTAGTTATTCCTACAGGGATGTA
CACCAACTACAGCAAATGGATCGCAAAAGAAATTGAAGGTTCGACTGGTTTTAACAAACCTATTCTTGCTGTTAACCCCTGGGGACAGCAAAG
AACGTCAAGCGTAGTTGCGAATGCGGCAGCAAAGATAGTCGGATGGAATAAGCAGTCAGTAGTAGATGGTATCTGGGAGTTATACAAATAGT
GAGTAAGTTAGAAATAACCTCAGATGAGCTACCGGGTCTGTACCAATCAGCAAATCAAGCATCTCTCAACGCACAGGATAACTACTTCAGAG
GCTTGCGATGGTATCTTATCCTTTTGGTATGCGCGGCATTTATTTCATATGCCATGCCGAGAGATGCATTGGGCGCGCTGCTGTCAGCGGGACT
TTTCCTAGTTACATTAGGAATTCTTATTTTCATTCGTGTACAGCGCCCCGATGACACATGGTATAACGGGCGGGCGGTTGCTGAATCAGTGAAA
ACAAGAAGCTGGCGGTGGATGATGAGGGCTGAGCCGTATGAAGACTGTGAGAGTATGGAAATTGTTGCTAAGCAGTTTATCAACGACCTGAA
GACAATCTTGGAGCAAAACAAAAGCCTTTCTCACTCGCTGCAATCAACAAGTGCCGCCAAAGACCCTATCTCGCAGACGATGAAGGATGTTC
GCTCAAGAAACGTTAAAGATCGCTTGTCGATCTACATCGATCAAAGAGTGCAGAATCAAGTTGAATGGTACTGGCACAAGGCTCGCTTTAACA
AGCGAAGGGCGCAACAGTGGTTTTGGGTCTCGGTGATTCTGCATGCGCTTGCCATTGCCATGTTATTGTATCGCATTAAGGATCCTTCTTTTTC
GCTTCCTGTAGAGGTTATTGCAACAGGTGCGGGGGCTGCATTAACTTGGCTTCAAGCGAAAAAGCACAACGAATTAAACTCTGCATATGCTTT
AACGGCGCATGAAATTGTATTAATTAAGGGCGAGTCGGATTCTGTCCACGACGAAAAGCAACTGTCTGAGTATGTAATTAATAGCGAGGCTG
CATTTTCGCGCGAGCACACTCAATGGGTCGCTCGAAAGGGCGATTAATGCATAACAATGCGCTGCTGTCGGACAAATTTTACGCTGCGCTCCA
AATTTGCCGCAGAGCGCGGCG

*gcu24*

TTATAGCTCAAGCAGCTCATCTCGAAACGTGAAGGTTTTATGGACGAAATAGAATTAGATTTTGGCGCGATAACCATAGATAACGCGACCTAT
AAGGGCGAAGGTTATAGATTTTATGAGGGCTTACTTGCTCAGATGCGCCAATTTAAAGAAAGCCCAGTTAAAGTTCTTCAAACCGATATAGTG
CACAATGAAGCGATCAATCATATCGGGCAAGAGATATCAAAAACCCGATCCTCGATTGAGCAGGCACTAAGGTCAGCAAATAAACAGCTAAA
GATAAAATCCGAAGTTATAGAAAATGCTCGAGCGTTGCTATCAGTTGATGGCAGCGAAGCTGAAATAGCGGAAGCGAGGCTGGAAAAATATT
ATGAATTTATTGGAGCTGAGAAGATCGATTCTGGAAGATATGCAGATCTATCTCGTCTTATGGAAATGTATTTTGCTACAGAAGCTCCTTTTGA
ATCCGGAAAGGACAAAAAAAATGAATTTCCAGATGCGATTGCACTATTAGCCCTTGAAGGTTGGGCTGAAGAAAATGAAATTAATATTATCG
CCGTAAGTCAAGATAAGGGCTGGAAGAACTTTTCTGAAGGTTCAGATAGGATTACGTTGGTATCCTCTCTAGCGGAAGCGCTAGAAAAGTTTC
AGCCACATTACAAGGTTGTAAGTATTATCTCTCATATCAGAGAGGACTCTCTTCTTGATGGCGAGAATCATGTACTTGAAGAAATTGAGCAAG
CAATTATCAACAGCGTCGATGGTTGCGATATTTGGGTTGAAGCAAGTTCATATATGCACTTTGAATGGGAAGACGTTTCAGCCTCCTATATTTC
CCATGAGCTGGATAAAGATCAAGATGGCTTAGTTAAAGTCAGAGTTGTAAGTATAAATGATGAGGAAATAGTTCTTAAAGTTGGTGCTACAGT
TGAAGTAGAGGTTGAGGCTAGCTTTGATTTCTCGGTAAGAGATTCGATAGATAAAGATTATGTTGGTATGGGTGGAAATGTTTGTACTACAAC
GGAGTCTTATCATACTGACATATTGTTATCTCTCACTGGAGATTTTTCTCAGGACTTTGACGACATCGATGTTGCTGAGATTGAAGTTCTTGAA
ACTATTGGGCGCGCAGACTTCGGCGAAGTTGAGCCTGATTGGCGGAGTGAGTACGAAGATGAAGAGCTATAACAAGGCGCTGCTACGGAAAA
TTTACTCGCTGGCGCTCCTAAATTTCCGCAGAACGCGGCG

*gcu29*

TTAGGCCGCTAAAGCAATTCCATGAGATCGCAGGCGTGCAATGACTAAATTCGCTCTCATCATATTGCTGGTCGCTTCAATCACGATCAGTGT
GATTCTGATAGTCGGCGTTGCCCGGAGCAACCGCAAGCCGATGGATAAAGCTTGGCGCATCCTCTTGCTCTTAGTCCCTGTTGCTGGCCCTCTT
CTCTACTGGCTCATCTATTCCGATACCCAGCCGCAGCCTCAGCATCTGCATAACCGGGAGCCGCGCGGCCACTACACCCACACTTGGATCGCC
ATCAAGCCCATACTTGAGCAAGGCCTGCGTCGGCGGGCTGAGGCACAACCTCAGGAAGATGACGATCATCATGGCAAGTAAAGCCACGTTTG
TGCCGCGATCTCATCTGGCGCGGCCTAACAATTCATTCAAGCCGATGCCGCTTCGCGGCACGGCTTAATTCAGGCG

*gcu32*

TTATGCGAATCAAGATATGAAATTAATCTCTAGTGAAAACATACCTGGTAATATTTCTGATTCAGTACAGTTTCTGAGGAGTAAAGGTATTTTA
GTTAGTGGCACTGATAGCTATGCAAATATTCCTCGCTATAGCCATGCGGCAAAATCAATGAAGCAAGGTATTTGGGTTCACTTAAATAGCCAA
TACTCTGATGCACTATTGCTACTTGCAAAGCCAAATGCAGAAATTAAAAGCGGTTTAACTGAAGAGCAAATGCTGGCAATCGAAACAGAAGC
AAAATCTGCTCTATTCAAGGCATCAAGTTTGTTTTTAGTCGGCTAGCTACTACTGTTCTAGTATTGTTTCTTATTTCTTTTGCAGCATATACTCT
TTTTAATATTTACAAGGCATAACAAAGTGCTGCAACACGCGCACTACGTACGCTGGACAGTTTATAAGTCGCGGTTTTATGGTTTTGCTGCGCA
AAAGTATTCCACAAAACCACAACTTATAAACTGCCGTTGAGCACGGCG

*gcu59*

TTATGCCCTATAAAATTATCTCCGTGTGATGGTCAAATTATGAAAATGAATGTGCATCTACCACCAAACAGTGCTGAGTTATCGGTCGTCGAAT
CATCAAGAAGCCTGCTTGTCGTAGGCGCCAATGGTTCCGGTAAAACCAGGCTAGGAACTTGGATAGAGTTTGATTCACCAGATCGCGATAAA
GTACATCGGATATCCGCTCAGAAATCCCTAGCAATGCCCGACTCAACGACACCAAAGTCGATTGACTTGGCAACATCCGAACTATGGACGGGT
TATGAGCGAGCTATCGAGCAGAACAATATCCAGGGATACAAACAAGGGCACAGGTGGCAAAGTAAGCCAGCGGTTAGCCCTCTGAACGACT
ATCACCAGTTAATGGTATTTCTTTTTTCAGACCATACTGAGGAAAGCGCCAAATACCTTTCAGCATCAAAACTGACAACCAGTAGAGTAGAGC
CACCTAAAACAAAGCTAGATGTAGTTAAGGAAATTTGGGAAAAAATACTTCCCCATAGAGAACTTGTGATTGGCGGCCTTCGAATCCAAACA
AAAACAAAGGACTCGCAACAAAGTATCTATAATTCCTCGGATATGAGTGATGGCGAGCGGGTTATTTTTTATCTTATTGGGCAATGCTTGGCC
GCCCCGTCCAGTGCAATTATTGTCGTTGACGAGCCAGAACTACATCTCCATAAATCAGTCCAAGCCCCTCTCTGGGATCAAATAGAGCAATCC
AGACCTGACTGCTTGTTTATATATCTAACCCATGACGTTGACTTTGCTGCGGCAATGGAAGAGACAACTAAGATTTGGTTAAAATCATTTAATG
GAAATCGCTGGGATTGGGAGTTAATTAAACAAGATGAGGCTATCCCGGAAGACTTATTGCTAGAGGTATTGGGCAGTAGGAAACCAGTTGTG
TTTGTAGAGGGTGTTGCAGGAAGTTTTGATTCAGCCCTATACACCTCAATTCTTAGCAACTACCTAGTCATACCAGTAGGCAATTGTAGTCAAG
TAGTACAATCAGTTAAAGCATTGAAAGCTAATCCGCAACTTCACCACCTTAACGTATTTGGCATCATTGATCGAGACAGAAGAGTTCCCGCTG
AAATCCAGAAGCTAGAGCAAGATTCAATATTCGTTCTTTCGGTCGCTGAGGTAGAGAATCTATTTTGCACCAAGGAACTATTAGCATTGGTCA
GTACACGACTGGCCCGTGATGAAAATGCTGATTTTTTATCTGTTTCTTCCGCTATATTTAAGAGGCTGCAAAGTGAGCTAGAAACTCAGGTCTC
ACTCCGTGCCGCAAGCGAAATAAAATTTCAACTAAATATGTTTGATGAAAAAGCAAAAGGTGCTGCTGCACTAACTTCTGCGCTAGAGTCGCT
AGCGGGAAAAATTGACGCAGGCGCCATTTACAATCAAGCGCTTTCAGAATTAAATTTGGTATTGTCTGCGAAGAATTATGAAGGCCTCCTATC
ACTATACAATCGCAAATCTCTATCTACTCAAGCAAGCTCAGCGCTCGGCCTTGCCAATGGAGAACTTCCCGAGCTTATAGTTCGGCTCGCGAA
AGGCGAATGTCGAAATGAAATTACGCAATCGCTAAAGAAGTATTTTGGCAACTTCGCTCCCTATATGGCATAACAAGTCATTGCAGCGGACGC
GCTTCGCGCGCCGCTGAATTCCAACG

*gcu68*

TTAGGAGTTAACACAACTCATGGCTCTTACAAGAATGGAGATCTTGGGTTTTCGGGGGTTCAGGACGCTCGGCACGATAAATTTTTCCGTTCC
AAACGGAGAGATCGGAAGCGGCTTAACAGTCATTACGGGGCCAAACAATGCTGGGAAGTCTTCAATCTTGGAATGCCTCAAGGCTCGCGCAG
GCCATCAACCGCCAAGCTTTACGGTTGGAGCGCGTAACGCCAACGTTGAAGAGGTTGGAATTAAATACGTTATCAACGGCAAAGAGGAAACA
ATTAAGTCCATAAAGAAAGGCGCGAGTGAAACCAAGAAAGAAGGAGTCGATTCTAATTTCCACGTCTTTGTACTTCCATCACGCAGGGCATTC
AATCCTTACTTTGGCCGATCTGAGCACTCCAGAGAGCAACATCTAAACAACTCAACTCTTACACCTCAACGCTCATCGATGCTGAGCGGATTC
GAGTACCGACTTTTCACAGTGCTCAAGAATCAAACGGCATTCAATGAAATCCTCCATGAAGTTCTGACCTTTAAGCCAGAGTGGTCGATAGAC
CAGTCCGACCAAGGACAGTATTTTCTGAAGTTCTTTAACGGGGATGACTCACATTCAAGTGATGGAATGGGTGAAGGAATCGTAAGCATTTTT
TCCATAGTCGACTCCCTTTACGACTCGAAGCCTGGAGACGTCATCGTAATCGACGAGCCAGAACTTTCTCTTCACCCCGCCCTTCAAAAGAGA
GTTGCCAACTTGCTGTGTAGATTTGCGAAAGATAGGCAGATTGTTGTTTCAACCCATTCCCCATATTTCGTCGACCTCAAAGCACTTTCAAATG
GTGGCCACTTGGCTCGTGTGGCGACAGGGGACGAAGGCACAATAATCTACGAAGTTTCTGCTGCAGCGAAAGACTCAATCAGCCGGCTATCA
GAGGGAAACTTGTACAACCCACACGTGTTTGGTCTTGATGCGCGCGAGCTATTTTTTCAGGAAGATCAAATAATCCTGACTGAAGGGCAAGAA
GATGTTCTGCTATTGCCACGGGTAGCGGAACAGGTAAAAGCAGAGATCACAGGGAATTTTTTCGGCTGGGGTGCTGGCGGCGCTAGCAACAT
CCGGCATCTATGTCGGATTCTTAAAGACCTTGGATACAAGAAGGTGGCAGGCCTATTTGATGGAGACAAGACCGAAGAGCGGGATAAGGCGG
CGCTCGAATTTCCGGAATTCCACTTTGCATGTATCCCGGCCAAGGACATTCGCACAAAACCACCGCGCAAAGCTACGGACGAGGTTCAAGGAT
TGCTAAACGAAAAATTGGAACTCAAGGAAGAATACGCAATGCAGCTTAAGACACTATTCGGCTCGCTTTCTACGCACATGAACTCCTAACATG
GCGGTTAACTCGGACGCTCCGCCGATAAATCCGGCTTCGCGCCGGTTACCTCTACG

*gcu72*

TTAGCCAGTCAACAGGAGAGAATGTGTTCCGTATCCTTCTCATTGCTCTGCTCGCCTTTGGCGTTTGGAAGGGCTATGAAAAATACCAATCTCA
ACGCCCAGCACAGCAATTTCAAGAAATTGATATTTCAAGTGGTTCTTCCAGGAGAACCCGAAGCATTGATCTAACACCAAAATTTCAGTGCGA
TGGACGAACCCATTGCTCTCAAATGACTTCTTGTGAAGAAGCCACCTTCTTTCTTAAAAATTGCCCGGGCACAAAAATGGACGGGAACAATGA
TGGTGTCCCGTGCGAGCAGCAGTGGTGCAAATAAATCAAGCCATTGGGGCTAACAATTCGTCGCAGCCGACCGCCTACGGCGGCGGCTGAAC
TCAGGGCG

*gcu76*

TTAGGCTTTGCATCCAACAGTACGTAATCTTCAAGGAGAGCTAAATGCGCTATCTGTATCTTGCAATCGTCATCCTCATGACACTCGTCGTCGT
CACATTCATGGTCCAAAATAGCGGCAGTGCGACGGTTCCCCTCCTTTCTTCTTCAGTGACGTTGCCTTTATCGCTGTTGACCTTCGGCACCTACT
TCTTAGGCATGCTGACAGGTGGCATGCTCATTGCTTTCATTCGATCATTGGTTCGCGGCGCAACCAAGCCGCAACCAGATCAGTAGATGCAGG
CATAGCCTAACAATTCGTTCAAGCCGAACTTGGTTCGTTACGCCGGCAACATGGCAGATTAAGCTTGCCATGTTGCCGGCTCCACTACGCAAG
TCGGCTTAACTCAGGCG

*gcu88*

TTAGGCCTCCCCACATCATGAACAGCATAAAGATTCGCAGAGCAACTGGATGGCAAGACAAACTGCGGGCCTACAAAGTTTTGCTCGACAGA
GTAGTCGTCGCGGAGATAACACAGGGTTGCCATGCAGATATCCCTGCAACAGCTGGCGCTCACACGGTTCAGCTAAAAATTGACTGGTGCAG
CTCTCCCTTGCTGCACGTAGAGGTGGGGAGTGAGGAAGATTTGACTCTTGAATGCGGGCCAAACGCCAAGCCGCTTCTTAGTTTGCTTTACGT

CACCTTCCTGTGTCGTAGGTACATCTGGCTAAGGCAGGCCTAACTGTTGACATATGCAGCGAACACCCAATCGTTAACCATATGTGCAGAGGG
ACGACTTCTACATCTATCCCCAGCCTAACTGGTCGTTCAACGCGGACGCCACTACAGGCCATGCCTTCGGCATTGTCATGGCCTGTAGCGGTAC
CCTCCGCACTTCGTGCTCCGGCGCCGGTTAACTAGGGCG

*gcu89*

TTGGGCCTTTCTATGACTGCTGATCCTGTACTCGAAATTTTGGCTGAAGCAAAGAAGCTTGCTCAGCGATACCGCGTCCTTACAGGCAAGCCA
CTCGGCATTACGGGAGAGGTTGCCGAGTACGAGGCGGCTCGTATCCTTGGCGTTGAACTTACGCCTGCTAGACAGGCTGGCTACGACGCTACC
GAAATCCGTGATGGTCAAACCTTCCGGCTGCAAATCAAAGGTCGGTGCGTTCTTGAAGGTAGCAAACCCGGCCAACGCATAGGCTCCATCGA
CATCAAGAAGGAGTTTGACGCTGTCTTGCTTGTTCTTCTTGATGGCAATTTTGAGGCTACTGCTATTTACGAGGCGCCGCGTGCGGCTGTAATT
GCCGCGCTTACGGCGCCTGGCTCAAAGTCTCGCAACGAGCGCGGTGCTCTGGGTATTTCCAAGTTCAAGTCTATTGGGTTCATTCGGTGGCAA
CGATCAGCCTAGGCCCAACCCGGCGCTCAACCCCGCTCCTTTCAGTCGCTGGACGCTGCGCGATAAAGCCGCGCAGCGCCGGTTAGCTCTACG

*gcu92*

TTAGAGCGCGCTATGTCCCACCTTGAATCACTCATCGTTGAATACCTTGATTGGCAGGGTTACTTGGTACGCCGCAACACCAAAGTGGGGCGG
TTAAAACATGGTGGATGGGAAATGGAACTTGATGTCATCGGTTTTAACCCGCACACCAGCGACCTTGTCCATTACGAGCCGTCTGTTGATGCC
CACACCTGGGATACGCGCGAGGCCAGATACGCCAAGAAATTTGAGGCCGCAAGAAAACTCATTTTTGCTGAGGTCTTTTCTTGGCTACCTCCG
GAAACTCCGCTACGCCAAATCGCCGTGTTTCCTTCTCACCCAAAAGGTCGAGACATCATTGCAGGCGGTCAAATCCTGTCCATTGACGAGTTC
GTTGCCGAGGTTCGTTCAGAAGTTGTCGAGTGTGGCGTTGCTTGCCGTAGTGCTATATCCGAAAACTATCCGTTGCTTCGCGTGTTGCAGCTTT
CCCATTGTGGCTACAACCGCGCTCTCTAACATTCCGCTCCAGAGGGACGCTCCGCCGGCAAGCCGGCTCCGCGCCCCTGAGCTTGTACG

*gcu113*

TTAGGTGGCTTGAGCTTCTGCTCACATAAAGGAAGCGCATTGAGAAACGTAGTCATCCATATTCCCCAAACTGATGCTGAACTAGAGGCTTGT
TTTGGCGCCTTTGCTGTATTGCGACCACATCTCACCGTGACCCAATTTCTGCCGCAGGTACGCAGGCAAGAACTTCAGGGCTACAAGATCGCG
GCCATCAAAGACGCCGAGCAAGTAGTCAGTGCTGCAGGCTTTCGCTTCTGCGAATTCCTGGCATGGGGTCGAGTGCTCTACATCGACGACCTA
ACCACTATACCCAGTGCCAGGGGTAAAGGTTACGCAGGAGTCCTTCTGGACTGGCTAACCGAACAAGCTGCTCGACAAGGCTGTGATGCAGT
GCACCTCGACACTGGTTACACGCGTCACGCCGCTCACCGCCTTTACTTGAGTAAAGGCTTTGAAATGACAAGCCATCATATGGCCAAACTCAT
TGCCAAGGACTAAGTTCGAATGGCAAATCTTCGTCGTTCAAGCACGCCACCTAACTGTAAGTTCAACGTGGACGCCAACAGCGGCCATGCCTT
CGGCATTCTATTGGCCGCTGTTGGTGCCCTGCGCCCCTGCGGGGCTCCGGCGCCAGTTAACTAGTTCG

*gcu114*

TTAGGCCTATTTGTTCAGAGCTTTAAAAATTGGCTTCGGTCAGGAAAAGCAAAAAACAACGTTTTTGCAGAAGTTTTTCACGGTATTGGCTTCT
TCAGCACAAGGCGCGCATTTTTCTGGTTGTGTCTCAAGAAACATTCTCGGCCAAATGCGCGCCAGAACATCCTTGGCTTCGTTCAAGGTCGCG
GGTTGAAAACCTCCGTTCTTGCACAGGTTTAAACCCATCGCTTCGGTCAGGTTGTTGTTTCGGTGCACCAGTTTTTGCAGTCAGAAATTCGATT
GTTCCTGGTCAGTTTGCTGCGGCGCAAAGCGCTGGTATTCTTGCGAAAAGCACTGGCCTAACCCATCAATCAACAGGGACGGTCCAAAGCTAG
CGCTTTGTCCCGCCCCTTATTTCAAACG

*gcu119*

TTAGGGTTCCTAATGGTACCTCGAACTCACGCATCAACTTCAGAGAGTCCAATTAACTGAATATGAATAAAATATTTTGCGTTTCCATATTGCT
CGCCTGCGTTGCGTTGACAGCATGTGCACGCCATCCGCTTGTGGCTGCCACAACGCTGGAAGCAACTTCATCAAGTGAGGTGAAAACAGCCAG
CATGGGCGAAACCGTTTACAACCATTACACCTGCCCGTGCAGTGAAATTTATATTGCAAAAAAATCAATAGCCATTGACGGGTTGCCAAGCAT
AACAGCCGGCTCTTTGTGGGTGGCGAAATATAGAAATACCGAGACAGGTGAAAAGTATCTTGTCAACGATAGCTACCACAACCAACTTGCCG
TAGTTTTAAAAAACGAAACAATAGCGCCTTCACGTGCAATAGCTCAATTTAGCGGAATGAAAAAACACAGAACATGGCCATTACAGTCAAGC
TCAGACTCAAATTCCATAACATTTAACGAATATCTGCCCAGTCAAAATTGGTGGCGCCTACAATATATTGGCCTAGACAAAAACGACAAAAAC
ATACTTCGATTCACCATTGAAACAAGATTTCAAAGCGAGATAGTTGGGCAAATTGAATACACTCACAACTTGAGCAATGGTAATGAATTTGTA
ATCAAGGGAGTACGATTTAAGGTTTTACAAGCGCGAAACGACAGCACACTAACTTACCAAGTGCTACAACCCTAACAAATGGTTCAAGTCGC
TCGCTTCGCTCACTCGGGACCGGCTACGGCCGGCCCCTTAACCAAACG

*gcu126*

TTAGAATATATGGAGTTTGAAATGTTAGCTGGTATAGGTATTGGGCTTTATTTGCTATCTGTGTACATCTTTGCAATGGGTTTTTCAAGCTTGAA
AGACAGTAAGAATCACGAAGAATATGGCGTTCAAAATCAGTCTCTCCATAAGAAATTCAGCTTTTTCCTGATTTTTCTGTCAATTATTATTGCT
GCATTAGCGACATTTATGGTTTTTAAGTAAGGGCGCCCCACATGGAGCTATATTATTTTTTGGCGGTTATTTTTTGCGTTGTTTCAGGAGCAAC
GCATTTTAAAATTACTAAACTCAATCCGCATACTCGTTTGCAATCATCAGCTGCTAATTTTCTTTATTTGGTGTTCTCAGCTTTTTCAAATATAG
GAATATTTGTTTTGTTTGGTATTGGTCTATACGCTTTTCCTTGGTGGCTCTCTTTTGTTTGTTTATTTTCAAGTTTATTTTTAGTTGGTTTTGTTTA
TTCATTCATCTATCCAAGACCTGCCTATGTTTTAATTTCTTTTCCATTAAGTGTTATTTTTACAATAATGGCAATTTGAAAATTCTAACAAGGCG
TAGCAACCCGCGCACTTCGTGCTCTGGACAGTTTGTACGCGGCG

*gcu128*

TTAGCTTGCTTGCGGTTATGTTGAGAAAATGGTGAAGCAAACAAAAATCATGGAAGTGCATTATGGTAAATAGTAATTCTGGTGTAGAGGGTG
GTTCTGGCTATTCATTTCAGCGTTGCTGTGTTGTTTTTCTATTATTTGAAGACTATGAAAAGTTAAACATAGAGGATTACTTTATTTGCTTAGAG
CACCATGAAGATTTTCTATTCGCATTTTTGGACGAAAATAGGCATTTAAAACAAATTGATACCTATCAAGCAAAAAAATCAAGAGATGACTGG
AAAACAGATAGTGACCTTTGTGAAATAATTGGAAAAATGACGATGGTGGGAAAAGAGCTTGTCAACGATCCTCATGACAAGTCTAATAATTA
TCAACATACCCTAAATTTTCTAACTAATCGAAATATATTATTAACAAGTAAACAAGAAAAAGGTGCTAAACAAGATAAAGTCAAAATCCAAG
TTTCAAACCGTCGAAAAAACTATTCTGATTTAACAGAGAAAATTAAGAAAAATATAGAACCAAGAATTTGCCAGTCAATACTCGAGATCACTC
AATTAAAGAATGTTTATTTTCAATATATTGATTTAGCACAAAGCTACAAGAGCTGGCAGAGAGAGCTAAAAGGCTTATCCATGGAGCACTTTG
GTCAGGAAGTGAATGATCATGAAGCTGTTATTTCAACATTGATGCGATTATTGGAAGAAGCAGAGCAAACATATAATGATAACAATAAAGTT
CTATTATCTGACCTAAGCAAAAGAGTAACTAAAGATAAAATAAGTGAAACTTTCAATATGTTTACTGTAAGTAAAAAGTCTTTTGATTTTTGG
CGAAAGTATTCGGACCAGATTTCAACCCAATTAGAACTAAAACTCCCTATACGGAGAAGAGCGCAAGAACTGCTTGAAAATTGTTTTGATTTT
TTTAAGGACCTTCAGCAAGTTGAATATCGTAAGATTTACAAGTTTGTTGAAACTAGAACAGATATTGACGAGAACCATATCTGTGAAGCTGAT
TGTATTGTGGATTTATATCACTGTTATCAACGTGAATTCCAGCCAAGATTAGAAAACTACATGGTGGCATTCGCTGTGATTGCAGCATATGTAG
AGACAAGAGGGATGTATGTCTAAATTAAACTTAAAAAGCTTATTGGTTTATTCTCCTAATGATGAGAAAGGATTTTACACAGAGTTTAGTGAA
TCAGTAAATATTATTCACGGACGAAATACATCTGGCAAAAGCACGCTTATCCAGTCAATTATATATGCTATAGGTATAAATGACTCAAAAGAT
AATTTGAATGATATTAATAGCGGTGATGTTTTTTTAGACTTGACTGTGTTTTAGAAAAAGCTGGGGAAAGTTGTAATCTGGTATTTATACGTT
CTGATGATACTTTGGTCTTGCAAAAAGGTAATGAGCCAGCTATTAGATTTGATGGCATTAACAGTAACAATTCTTTTGAGTATGGAAGGTATA
AAGAACTTCTTTCAAAATTAATTGGCTTCAATTTAGTCCTACAAAAACAATCTGAATTAGTAAGTGCTCCGTTAGAAGCAGCACTGTTGCCGTA
TTATGTATCTCAATCTGTGGGATGGGTTTATATACGTGAGTCTATTGGAGATTATAGGTTCTATAAGGATTTTAAATTTGATTACTTGGATTACT

ATTGTGGTATTGAAAGTGGTCATGAAAGAATAAAAAAATATGATCTTGAGAAAGAAAAAAAGGAATTAAACTTTGAGTTGAAGCAACTTGAT
TCATATGAGGTCAAAAATGCAGATCTGAAAGTGTCAAAGTTACTTGATGAGCGATTCAAAGGCGAGGCTGAAAATTATCTAGAAGGCTATCA
AAGTTTAAACAAAGAGTTATCAGATAAAGAGGCAGAACATACGAAATTATGTAACAAATCAAGCATGTTAAGAGGTAGGCAAAAAGTATTAT
CTCAAATAATAACTAACATAAAAAATCAAAGGCCAAAAATTGATCAGTGCCCAACATGTGATCAACACTTACCGGGCGATTTAAAAGAGTTT
TATAAGTATAGCCAAGATGTTAATGATGCCTTAAAAGAAAAGGATAGAGTAAAAGAGAATATTAAAAGCAACGCTTCGAAATTGAATTCTAT
TGAGAAGAGTTTAACCTATTTGAGAGAAAAAATAGAGAATGAATACGGGGTTTTGCGACGCTTAAAATCTGAAAATATTACATTTGATTCTTG
GCTTAATCACAATGCCAACTTACGAATGTTAAAAAACATCTCTATCAAGAAAGCTGCATGTGAAAAAAGGATTCACGGAATAAATAATGACA
TAGAAAAGCTTGGAAGTGGTGTTGATATTGACTCCTTGAGAAAAGCCAAGGAAAGAGAGTTTCTTTCTATATTTAAACGTAAGGCTTCCGCAC
TAGGTGTTAAGCTACCAAAAGAAACCAAATATCAGGATCTGTATTCTATAAATTCATTCCCGTACCAAGGGGTGGAGTTGCACCAGTTGCTAA
TGGCCTATAATTTTTCATTCTACGAAATGGTCATAAAAGCACCAGCAATCCATAGCTTTCCATTTCTACTAGATGCTGTTTTTAAAGAAGATAT
TGATTCTGAGAGCCGAAGTAATATATTTAAATTCCTTTCGCAAGAAACTAAATCTAGTGGGCAAGTGGTTTTTTCTGTGGCTGAGTATAAGGG
GGATGAAACATCATCTAATCCTCTTTTCAATGTTGATGCCATTAAAAAGGAATATTTTCCTGAAGACACAAAATTAATATGTATTGGTGATAGT
AAATCTAAACGGGCATTTTTATCAAAAGCAGCCTTAATAGATAGCTCATTAGTTGACAATACAATGAGGTTGTTGGAAACAGTCTAACTTAAT
ATTTCAATATCCAAGCTAACAAGCCAAAGCACGCGGACTCAGCAAAGCTGAGCCGGT

*gcu135*

TTAGCGTTTTAGGAGGGTAAAGCAATGAGTTTGGAAGATAAGTTGTCTGGATGGACAGGCCCTTCAAGTAGCACGGAGCAGGATAAGCAAGA
TAGAACGGAGCGGATGATTCGCCAAGCGATAGATGCACATGAACCATTCAATAATTGTTCATTGAAAGTTTTTGCGAAGGGGTCTTACGCTAA
CAATACTAATGTGCGGGCAGATAGTGATGTAGACATAGCAGTGGAGTGCACGGAATGTCGATACTGGGGGGAATCTGAAAAAGGTAATCACC
CTCCTAGTAGTGGATCATATGAGGGAATTTGGACCCCAGAAAAGTTAAGAGCAGAGCTTCTTTCTGCTATGAATACAAAATTTCCGGGTCAGG
TAGACTCTTCGGGATCTACTGCGATACAGATAAATTCTAACTCAGCACGCGTAGATGCCGATGTAGTTCCATGTTTTAGCTATCGCTACTATAT
GAACCATGGATCGAGAGATGGAACAAAAATATTCAAAACTGATGGCAGCAGCATCGTTAACTATCCAGTTCAACAAAAGGAAAATGGGATA
GCAAAGAATAATCGAACCGGATATGCCTATAAAAAAGGAGTGCGCCTACTAAAACGCGTTGAAAATGTAATGGCACAAGATGGCACATTTCG
TGAGTTGCCATCATTTTTCATGGAGTGCCTAGCATATAATTGCCCAGACAATACATTTTCTCACCCAACATGGACAGAGTGTCTACGAGCCATG
CTTTTGCATATTTGGAATAACCTTCAAGGTGATGAGCCGACCTCTGGACGCTGGTTAGAGGTGAGTGAGTGCGTTTTCCTGTTTCACTCAAATC
AAAAATGGACTCGCAAGGATGGTCGCGAGTTTGCTAAAGCGGCGTGGAATTACTTTGGATTCAAATGATGCGTAACGTGACAATTAAAAGCT
CGCTGTATCTTCTAGTCGGAATTTCAGCAGTTGCTTGGTTTTCGCTCGCTTACTTTAGCGGTCTTGATTTGTCAAAGGTAAAAGATTTCTTTGGG
CTTGTACCAAAGGTTGTAACTATTGATATTTTAGTAGTTGCAGTATTTGTTAAATGGGGCTGGAAACTTAAAGTATTTAGAGGGTGGCTTGTTC
CTTTTCCTAATTTAAATGGAAGCTGGGTGGGTTTTATATACTCCGACTGGAAGAATCCCGAAACAGGAGAAAAGCCACCTCCCATACCCGTAA
TGCTCACAGTAAACCAATCTTTCTTTCATATTCATTGCATGATGCGCACGAGTGAAATGGAAAGCTCTTCATATTCTGAGGGCTTTCTTATCGA
ACCGGATCGACAAATTAAAAACATAGCATATTCCTATACTAGCAAGCCCAGATTAAGTTTGAATGAACGGAGTATTCCACATGATGGAACAA
CAGTATTTAAGATCATTGAAAAGCCTAAGCAAAAACTAGTAGGCAGGTATTGGACGGAGAGATTAGCAAAGGGTGAAATCGTCCTTCAGTAT
CACTCTAAAGAAATGCTTGAAGAACTTCCGGAAGAGTTGGGAGATCATCCCGTAACGGAAGATGAAAATCGACGCTAACAATCGGCTGCACG
GCGACCGATTTTCCGCCGCTTCGCGGCTCCAAACCGGTGCGTGAGCTGGGCG

*gcu142*

TTAGGCGATTTCCCCGATTAAGGAGTATGGATGGGTATTGAGAGTGAATTGGTAGGCTTCCTAGAAAGCACTATCAAGGATGGTGCGAATAA
GGCTAGGGATATAGAGATTGTAAAATTCTACTACGGCCTAAATGAGTCGCCTTGGCCCACTTTAGAGGAAACTGCAAGCAGGTTTAGTGTTGG
TACAAGAGAAAGAATTAGGCAATTACTCAATTCCAAATTTAGAGATAATGTCAGCAAGCATTCTATACCGTCTTTAAACACTTTGGTTGAAAT
TTTACAGTCCAGAGAGTATTGGCTGACATCAGAATTAGAACGTGAAATTTGTGACGCTGGGCTAATCGACAAGGAAACCCATTTAAAGGGAA
TATTTAACCTAATTGATGATGTAAGTCTTAATTGTGAATTCGAATTTTACACCCCAGATCTAAAGCGAGCCACGAGAAACTCAATTTTATCTAG
CAAAAATATATTCTTGCTCAGGACGTCTAGTGTCAAGGATATCGAAAATCTATTCAAAAAAGCTCAAGGCTTACCTGGGCGATGTGGAATTGC
TAAATTAAATTATCTGGAACAAGAACTTGCTCAGTACTACACGCTCGTTTCATTGCTAATTGAAAGCTCACCAACGTCTTGGGTTAGAGTGATT
GGAGACGATTACTGGTATATTTTTGAGAATAGAGATAACACCATTATCAACTATTGCGAGAAGGTATTTGGTGTTATTGAGCATTGCGACGCC
GCCAGATTGGCAAATACATTTAGGAATGCTTTAGATGGGAGAACCTATAAGTATCCTTACCCACCTACAGAAATTATTGAGGAATACTTGAGA
AGTTCTGTCTACATGGTTAACACTGATTCTGGATTAAAGTTCATAGGTGAAACCACAAGGCTGAATGAAATTGAAGAGGACATAGTATCTTTC
TTTGATTCTGGTAGAAACGCATCATTTCCTGAGCTTCGAGAATACTTATCTCAAAAAGGTTATGGGCAGCCGCATATATCAAAATCTACAAATT
TTTCGCCTTTGGTATATGTTGATAAGGCCAAAGGTCGAAAGCACTATGTGTTCAGTCTCATTGGACGTAGAGTTTCAGAGCTGGTTGATAATTC
AGATATAGATATATATGAATTGTATCTCCGTAGATTGCGAGCTTTGCTTGATGTTGGGACTGATGAAACAAGAGAGCAAACAACGAGAAAAG
AGCAGCACATATTACAAGAATGGCTATTTAAAGATAAGACGCATGAAGATTGTGCTATTTGCGGTAAAGAGTTCAGCGTGAAATCCTTGGTCA
CTGCACATAAAAAGCCGAGATCAGACTGTAATGATGCAGAACGCTTAGACCCATATATTGTTATGCCAGTGTGCGTGATGGGGTGCGACTACC
TTTATGAGAATATGTATATCTTTATTGAAGGCACTGAAATTAAGCGCGGCCAATATTTTTCGAATGCAAGGGCTGAATCTGATTTTATTGAAAA
TCTAGTTGGGCGAGAGGTTGATTCAAAATGGCTTCTGGGCAATCCGTCCTATTTTCGTTCGCCTAACAAAGCATTGCAGCGGACAAGCCGCTG
AACGCGGCG

*gcu149*

TTATACGAAGAGCAAGGAATGAAATGAATCGGTATCATTTAGCACAAATTAATATAGCTCATGCGCAAGCCCCAATGGATTCTGCCTTAATGA
AAGGGTTTGTTGATCGTTTTGCTGAAATTAATGCACTCGCGGATAGTTCGCCCGGTTTTGTCTGGCGGCTTCAAACTGAAGACGGCGACGCTAC
TGCACTACAAGCTTTCGATGATCCATTATTGCTTGTAAATATGAGTGTTTGGGAAAATATTGATTCATTAAAAAACTATGTTTATAAATCACTT
CACGTTGAACTAATTCGTGATCGTAATGCATGGTTTAATAAAATGACTAAGCCTCATCAAGTGCTTTGGTGGGTTCCAGCGGGGCACATTCCA
TCTGTAACAGAAGGAAAGGAAAAGTTAGAGCTATTGCAAAAGGTGGGGCCAAGCATAGCCGCCTTTACATTTGCAAAGCCATTTAGCTATGA
GTCATAAATCGTATAACAAGGCGTTGCAACACCGTTTCGCTTCGCTACACTCGACAGTTTGTAAGCAGCGCATTTAGCGAATCGCTGCGCGAA
GTTTTTCGCACATGCGCTGCTTACAAACTGCGGTTGAACTTCACG

*gcu163*

TTATGTAGCAGAATAATATCTATGCACCTCCTAAGGCCAACCAGAAGAAAAATTTCCACCACAGTACTTTTTATAAGTATTGGTTTTGTTTTTT
CAGTCGTCAGCGACCTAACCTCAAAAGCATACAAAGAATACGTCAAGAATACCCACTTCGAGGGCCAGCTTCTTGAAGATTACGAAAATATA
ATAAAACAAGGACCACTAACACCCGCAGTGCCAGACAACCGAATTTACGAACTCGTAAAAAAAATCCCACCAGAGGCGCACAAAAAAGTTG
AGCTAGCCGGATTATTTAACATTTGGTTCACGGTTATTTTTGGGCTTCTAATTACATACCTTGGAGCGTGTGTTGCACATCGCGAATCTAAATC
GCGTGATGCCACATAACAATTGGTTCAAGTCGCTCGCTTCGCTCACTCGGGACCGGCTAAAGCCGGCCCCTTAACCAAACG

*gcu167*

TTAGGCTTTAAAACCCTCACTTCCGTTGCTAAGTTGAAGCTAGATATATTTCACCTGATGATTAAATACGATGACAACACATATTGATAAATTA
GCCTTATTCTCGATCTTAGACAAAAAATTACTGGTTGTTCGCTCCAAAGGAAAATCTCTTTTTTATCTCCCAGGCGGCAAACGTGAACTCAACG
AAACTGACCATGAAGCACTGTTACGTGAAATATCTGAAGAATTAGCGATCAATTTAATACCTTCAACAATAACCTATGCCGCAACACTTGAAG
CTCAAGCTGATGGCAAACCTGAAGGCTTGCTAGTTAAATTAACCTGCTATTTCGCTAACTATGATGGGGCTCCAACGCCTGCCGCTGAAATTG
AAGAGTTACGTTACATTGACAGCAATGATATTGAGATATGCTCAAAAGCAGCCATCTTATGCGTTGATTGGCTCAAAGCAAGTAACTACATCA
ATTAGTGCTGACAATATTCATCACGTAGAATACAAGCGTAGCCACAAGGCATGACAGAGATGAACACAAATGAAACCGTATGAATGGATATT
GTTCGATGCTGATGAAACACTTTTTGAATTCGACGCTTTACGTGGCTTAAAACTTATGTTTTCACGCTTTGGTATCAACTTTTCTGATGCTGATT
TTTCAGAATATCAATTAGTTAATAAACCACTCTGGGTCCAATACCAAAATGGTGAAATATCAGCAACGCAACTCCAGTATCAACGCTTTCAAT
CTTGGGCTGAACGACTTGATGAAACACCCAAAACATTGAATAGTGCTTTTCTGTCCGCGATGGCTGACATCTGTGCACCATTACAGGGTGCGG
CCAATTTGCTGAATACGCTTCGTGGCAAAATGAAACTCGGCATTATTACCAATGGCTTTACTGAACTTCAGCAAGTACGCCTTGAACGTACTG
GGTTTCGAGACCATTTTGATGTGCTGATTATTTCAGAGCAAGTAGGTGTTGCTAAACCACATCCTGAGATTTTCGAGTATGCGCTTTCTGCTAT

GGGTCATCCCTCCCGAGAACGCGTTCTCATGGTCGGTGATAATCCTGACTCAGACATATTGGGTGGTCTTAATGCAGGACTACATACGTGCTG
GGTCAATGCAGACAATAAACCTGAACCAACTGATATAAAACCACACTATCAGGTTTCATCATTATCAGAGCTTGAATCACTTTTGGTACAGTC
CTGAGTATTAAAAAGCGCCTAACAAAATTTTCATGTCGGACTGCACTACGTGCAGCCGCATAAAACGAAG

*gcu167.2*

TTAGGCTTTAAAACCATCACTTCCGTTGTTAAATTGAAGCTCGATATATTCACCTGATGATTAAATACGATGACAGCACATATTGATAAATTAG
CCTTATTCTCGATCTTAGATAAAAAATTACTGGTTGTTCGCTCCAAAGGAAAATCTCTTTTTTATCTCCCAGGCGGCAAACGTGAACTCAACGA
AACTGACCATGAAGCACTGTTACGTGAAATATCAGAAGAATTAGCAATCAATTTAATACCTTCAACAATAACCTATGCAGCAACACTTGAAGC
TCAAGCTGATGGCAAACCTGAAGGCTCGCTAGTTAAATTAACCTGCTATTTCGCTGATTATGATGGTGCACCAACACCTGCCGCTGAAATTGA
AGAATTACGTTACATTGACAGCAATGATATTGAGATATGCTCAAAAGCAGCCATCTTATGCGTTGATTGGCTCAAAGCTAGCAACTACATCAA
TTAGTGCTGACAACAATATTCATCGCGTAGAATACAAGCGTAGCCCAAGGCATAACAGAGATGAACGCAAATGAAATCGTATGAATGGATAT
TGTTCGATGCTGATGAAACACTTTTTGAATTCGACGCTTTACGTGGCTTAAAACTCATGTTTTCACGCTTTGATATCGATTTTTCTGATGCTGAT
TTTTCAGAATATCAATTAGTCAATAAACCGCTTTGGGTCCAATATCAAAATGGTGAAATCACAGCAACTCAACTCCAGTATCAACGCTTTCAA
TCTTGGGCTGAACGGCTTGATGAAACACCCAAAACATTGAATAGTGCTTTTCTGTCCGCGATGGCTGACATCTGTGCACCATTACAGGGTGCA
GCCAATTTGCTGAATACGCTTCGTGGCAAAATGAAACTTGGCATTATTACCAATGGCTTCACTGAACTTCAGCAAGTACGCCTTGATCGTACTG
GGTTTCGAGACCATTTTGATGTGCTGATTATTTCAGAGCAAGTAGGTGTTGCTAAACCACATCCTGACATTTTTGAGTATGCGCTTTCTGCTAT
GGGCCATCCATCTCGAGAGAAAGTTCTCATGGTCGGTGATAATCCTGACTCAGACATATTGGGTGGTCTTAATGCAGGACTACACACGTGCTG
GGTCAATGCAGACAATAAACCTGAACCGACTGATATAAAACCACACTATCAGGTTTCATCATTATCAGAGCTTGAATCGCTTTTGGTACAGTC
GTGAGTATTAAAAAGCGCCTAACAAAATTTTCATGTCGGATGGCACTACGTGCCACCGCATAAAACGAAG

*gcu182*

TTAGGCCCCTGAAAAGCATCGCTTGACCAGCTATGCGCCAATGGCGTATAGTTCGCTGCATGCTCACCATCATCGAGTCGCCGCTCTTCAGCA
AGATTTGGGCTGACTACTGGTCAGTGGAGGAGCACGAAGAGTTCACGGTCCACATCGCCGCTAACCCCGAGGCCGGCGATCTGGTTCCCGGCT
CGGGCGGTTGCCGCAAGGTCCGGTGGACTCGCCCTGGCATGGGCAAGCGTGGCGGCGTCCGAGTCATCTATACCGTACGCCTGAAGCGCGGA
ACAGTAGTGTTGCTCACCATCTACTCCAAGGGCGCGAAAGAGAACATTTCCGCACATGTACTACGCCAGATCGCGGAGGAACTGGATCATGA
CTAGCAAGAAGACAATTTCGGCGCGTGATGCCGAGCGCAACATCGGAGACGAGTTGCTGCAGGCTATTCGCGACGTCAAGAGTGGCAAGTAC
GGCGCCAAGTACAGAGTTGAGGCCAATGATATTGTCAAGGCGCGCCTCAAGTCGGGCCTTTCTCAGGCTCAGTTTGCTGCCGCTCTGCGCATC
TCATCCCGCACCCTTCAGCAGTGGGAGCAAGGGCGCCGTCAGCCTTCTGGTGCAGCCGAGACTCTCCTTAAGATCGTGTCTCGCCATCCCGAG
GTTCTTCGCGAGGTTGCAGGGGCCTAACAATTCATTCAAGCCGATCCCGCTTCGCGGGACGGCTTAATTCAGGCG

*gcu189*

TTAGCGGCCTGTAGGGAGATAGGAATGCTCAAAAAGGACAAAGAGGACTTCCTTGCCTTAGTGTCTGGTGATGCGACGCTGACACAGAAGGC
CGTTGAGAGCCTGAAATCCAGAGAGAGGGGGCATTTTGAAGACCATCCAATGTCTTCAAAATTCGTAAAGCAATATCTGAATGAACTAGTTG
ATGTCTGGAGAAATAGCCCTAACGACGGGGCTAGAGAGTGGACATTGCAATTCATCGCAGACGCCGGGGTTGCAAACGAACATACAAAGCCC
CTTATCTGCGCTGCGCTATCTGATCGCTCCTGCATGTTTCTCCCAACGGTCATCTACCTGATGAGCACGTCGCCAGAATTGTTCGCAGATAGCG
GAAGCTATCTAGTTCCCCTGACGTCACACCCGGACAGAGAAGTTCGCTGGCGGGTCGCGTATTTCATATCCAAGTTGAAGTCGCCGGATCAAG
ACATGAAGAAGTGCATCGAGTATCTGAAGAGCGACAGAGACGGAACCACGCAGCTCTATGTCAGAGCGTGCGCAGGCCGCTAACAATTCGCT
GCAGGCTGACGGCCCTGTCGGGCCGCAGCCTGAGCTCAAACG

*gcu199*

TTAGGCGCTAGGAGGGGTAATGAGCTACGACCTGATGGTCTTTGACCCAGACGCAGCCCCAAAAGATCGTGAATCTTTTGTGGCCTGGTACGA
CAAGCAGACGGAGTGGTCAGAGGATCACAGCTACAACGATCCAGTGGTGAGCACACCTTCGCTGCAAGCTTGGTTCCGAGAGATGATTCAGT
TCTTCCCGCCTATGAGCGGCCCTCACGCAGTGGACTCAGATGCATCTGAGGTCACGGATCACTCCGTCGGAAAGACCGTTATCTATTCAGCAT
TTGCTTGGTCAGTTGCAGAACAGGCATATAACAAGATGCGAGAGCTCGCCATCAAGCATCGCGTTGGATTCTTTGACGTAAGCTCAGATGACG
GTGAAATTGTTTTCCCGTTAACGCCATGAGGGCAATCGGCTTGGCGCCTAACAGTTCATTCAAGCCGAACCCGCTTCGCGGGTCGGCTTAACT
CAGGCG

*gcu217*

TTATGCACGTCCACTAAAGAAGTTTTACATGCCTGAAAAATTAATGCAGCAATAGAGTTAGCGAACGTTGGTACGTACTACAACGTCCTTTTG
AACGTGTAAAGCTCAAAAGGGTAAACCGGCTGATAAAGAGGATGTATATTATGTCAAAAAATCGAAGCGAGGACGTTTGGTTAGATCCTGAA
ACCGGGCTGATGTGGCAAACGGCAACAAGCGATAATAAGATAGATTGGAAAGATGTAGGTAGGTATGCTGAGCAATTAAACCAATTAAATTA
TGGAGGCTATAGCGATTGGAGAGTGCCGACTGTATTTGAATTAGAAACGCTAAATACTGAGAAACCTTATTTCATAAAAGGTGGCCCTAGTTC
AGGAATTTTTATTAAAGAGCCGCTTTTGCATTCTATGGATTTTGACCGTCAGTCTTTTTGGACATCAAGCCAAGCTGATGAAATAGATGGAAAT
AGAGCGCGCACATTCCTTGTCAATTTTTTTACGCAAAGAGAAAGCCACATAGTGAATCACGTAGTGGCTATGCTTGTTATTCCAAATATGTCA
GGTGTGTTCGTTACTGATATGAAAACCTTCATTTTCAACCACGTTTCTTAATAAAACTATTTCGCAGCTGGTTGGCAAAAAAGTTTCTAATTGG
TCAAGTAACAAAAAATATTTTGCATGGCACAAAGATCGATGGGCAGAAAATGCATAACAAATCGCTCAAGCACCGCCACTTCGTGGCTGGAC
AGTTTTTAAGTTGCGCTTTTGTGGATTTGCTGCGCAAAAGTATTCCACAAAACCACAACTTAAAAACTGCCGCTTAGCTCGGCG

*gcu223*

TTGGGCGTCATGATCCGGAACCTTCACGAATATGAGTCGTGGCTTGACGAACTAGACAAGAAGCTCTTCGTCAAGAACAACGCAATTCATTCC
GAATTCGGCCCTGGTTTTTCGGTGGCGTTGAGCGCATTCCACACAGAAGAAGAAATCGTCGCTTACTCTGTTACATTTGCCCGCACTCTGGACC
AACATCTGTTTGGATTGCCAATCAAGTATCTAGTCGAGCGTTTTGTTCGGTTAATTGTAAATGTCCAGCATCTTCCGTTTGAGTCAGAAGAGCT
AGCGCGAAAGGCGAGTGTGGCAGTGTCACAAAGACATTCGGTTTGGGTTCCACCAAATACGTAAAACAAAGCCACATGCACACGCCGCCCAA
CACTACGGTCAAGGCCGCTCCCTTCGGTCGCTGGACGCTGCGCGATAAAGCGCAGCGCAGCGCCCCTTACCTTCAACG

*gcu228*

TTAGCGAGCATCCTCAAGATTTAGCGCATGCCCCAGACAGACCTTTATGGTTTCGCTCTCCTGGCATCCGACATGGCGCCCGATGTCGAGTGC
ATTGACGGATTGCTGTTCTTGGACCGCGGTCACCTAGGACACGACGCATTGCGCGCAAGAGTGCAGACATTCTCTAGCCCGATTGAAGCGCAA
TCCTGGATGAACCTCGTTCCGATCGATGACTTCATTGATTGTGGTGTAGCGAATTGGTCCAGCTCAGACAGCGCACTAAGCGCTCTCGTCGAC
ACATATCGGCGATCATGGCTCTCGATTGTCCGGGCCAAATTCGGCGATGTTCCTGGAGTTTCGGTGGACATTCTGGTGGACGACGACGCGGGC
GACGTTCTTCTCCGATTGACCCAGACGCAGCGGATCGCGTGATGCTCGCTAACAACTCGCTGGAGCGCAGCCGTGCTCGATAAGGGGCCCATT
CACGATGGCTGCTCACGGCGCGCTCAGCTCCGGCG

*gcu232*

TTAGACCTCATGAATTTCGTATCCCCACAGGGCGGCATCGAATTTGAAGTCCCGGATGACTGATGGCGCTTCTGTGATATGACGTCTTGGAAC
AGAGGTTCCGACTACTATCCCTACGCCAATCCCGGTGGCGACATCGAAGACGTTGAGGTCGTGTCGATCTCGGACATCGATCCACCAGCACGA
AGCGTTGCGTCCCTCTGTTCAAGAAATACAAGTTCGTGCCGGTGCTACTTGCTTTGCAGTCGCCGGAATGCGCGCTGCCACCGATAACGGTCC
ATGCAATCGCGCGTCCCGGCGGTGCGCGGTACAGAGTCCAGAACGGATTCCACCGCTACTACGCATCCGTTGCTGTGGGTTATCGGAGCATTC
CGGTTATCGTTCTACCCGAGATGCCGTCGTGAGGTCTAACCACGCGCTGCACTCGGACGCGCCTGGGCTGTCACGCCCTTTGCAATGCAAACC
GCGCGACAGCCTGCGGCGCGCCGGTGACTGCGAGCGTTAG

*gcu233*

TTATAGCTTTAAAAGGAGTTCCAATTGAGTAATGTAATAGCAGTCTTTGCGAGTGCAAGACGTAATGGGAATACAGGAAAATTTATTGACTGG
ATAGCGGGTGAGCTTGGAATGGAGGTGGTTGATATTTCTGAAAAAAATATAACGCCCTATGACTACGATCACAAGAATATTGGCGATGATTTT
CTTCCAGTAATACATAAACTTCTTGGGTACGAAAAAATCATTTTTGTAACACCTGTGTATTGGTATAGCCCCAGTGCTCAAATGAAGACCTTTA
TTGACAGAACATCTGATTTTTTGGACTTAGATGAGTTAAAGGATATAGGCAGACGGTTGCGCGGAAAAACAGCTTATATTGTATGTACTTCAA
TTAGCGCCGATGCTGATAGCTCGTTTCTTGGCTCATTAAAAGACACTTTTCGTTATCTGGGCATGCAATATGGCGGTTATGTTCACGCTAATTG
TGAAAATGGTTATGTTCAAGATAATTATAAGCAAGATGTTGAAAACTTTTTGCGTACTGTCAAATCCCAAGAGCGGCTATAACAAGTCGTTCC
AGCACCGCGCACTTCGTGCGCTCGACAGTTTGTAAGTTGCTTTTTTGTGGTTTTGCTACGCAAAAGTACCCCACAAAAAACCAACTTACAAACT
GCGGCTGAACTTGGCG

*gcu236*

TTAGGCATTAGAACTTCACGGCGGATTTAAAGAAAAGGAGAAGACATCGTGGCTCAATGCACTGCCCCAGTGAATGGTCATCGCACAGCGAG
TGCTGCGGCCAACTGCCCAGCTTGTCGCTATCGCTCCAGTGGCTACCGAAGCAGTTATGGCTACGGAGGTGGTTACGGTTCGTCCTACTCTTCA
GGCGGCAGTAGCTACAGCGGCGGCGGTGGGCGGTCTACTAGCGGGGGCAGTTCTAGGCCGCGCTGGTCGCGCGCCAACTCAACGGTTTTCTA
CACACCAGCGCAGGTGGTGTCACTTACGCCTGTCCGTGAAGCCGTCGAAGCGCAAACCGCAGCGAAGCCGGATCTCCGCGATTGCTTCCTTTG
CCATGCTTGGGATGATCGCCAGGGAGCCGCCAAGCAATTGCACGATCTGCTTGAAGGAGCTGGGGTCAAGGTGTGGTTCAGCGAAAAGGACC
TTGGTCTTGGCGTTCCGATGATGCGCGCCATAGACAAGGGCCTAGCAGCCTCAAAAATTGGGCTTGTCTTGGTGACCCCCGCGCTGCTCGCAC
GGCTACCGAAAGAGGGCGTCGCCGACAAGGAGCTATCAACGCTATTGCAGGGCAATCGGCTGGTGCCGATCGTACACGGCACTACGTACGAC
GCGCTCCGGGATGTGAGCCCCATGCTCGCTTCGAGAAGCGGTCTGGACACATCGGAAGATTCGATGTCAGTGGTAGCTACCAAAATTGCAGA
ACTAGTAGCAATTTGGAGCTGACCCTTGAAAGATGAGGGCTGCCAGCTAATGCCTAACACGGCATTGCAGCGGACTGCCTTCGGCAGCCGCTG
AACTCCAACG

*gcu238*

TTAGGTTGACGCCTTCGGCGCTAGGAGGAAATGTGAAAGAACTAATATCATTTGTTATCGTAATGGGGTTGATTGATAGTTTAAATCCTTTTTC
AGTTGGACTACAGATAGCGCTTCTTCCCCTAACTAAAAAGTCTCATCATACTTTGTATTTTGTACTAGGCACATTTGTTACTTATTTTATTGGCG
GAATTTTAATATATTATGGTTTTGTTTCTATTTTTAAATCTTTTTGGTCGGATCTAAACATAGACTTTACTAAAACCCCATATACAATAATTGAG
CTTGCTCTCGGCCTTGGACTTCTATCATATGCCATTTTTGTTTCCTTTATAAAAAAGCAGAATAAAGATATAGATAAGAAGGAATTTTCAGTTA
AACCATTTTCACTTTTTTTATTAGGAGCAACTGGAACCTTATTTGATCTTCCAACAGCAATTCCCTACATCGCTGTCCTAGGGAAAATGGGAAC
AATGGAACTAAAACCCACATTAACGATCATTCTTCTAGCATTTTATTGCGTTATATATCTTCTGCCAATGCTAGCAATTTTTATTATCCATCATA
CATTAAGAGAAAAATCTGAAGAATTAATTAACAAAATTAAAAAGAGTTTTGATAAAATTTCGAATGTTCTCGCTAAAATATTTAGTTTTGGAT
TTGCTGTTTTTTTAATAATAGATTCAATACTTAGTATTACAGGAAATACCATTAATTATTAACCTAACAAATGCTTCAACTTGACTTGCCCTTTG
TCACGGTTTGTGCTACTCGCTTCGCTCGCTTGCGCACAAACCGCGCCAATCCCTCGCTTCGCTCGGGAACGGGCTGCGCAAGTTAAGCAAATG

*gcuA*

TTAGGCGTCATGGGTCAAACGCTCACAATCGAACCAAGCGGCGAGAGTCGCGGGCACTGCGACTGCTGTGGGAATTCCTCTCGAACCGTTTGG
GGTTATGTACAGGACTCAGAGAGGACAATCTGCGCATACTTTGTTCAATGGACTGTGGGCGCTCCAGATCACTTTCCCAACTTTGACTTTCTCA
TCGGAACATGGGGCGACGATGCAGTAAGTGACAAAGTTCTTTCCTCGTGGCTATTCAATCCCGAAGCAGGTTCATTCATGATCATTGATGCTT
CTAGTCGTCCTGCAGCGGCATCGGATATGTGCAACCACGCTCTTAGTCGTGCCGAGGTTCTTGCCGCACCGGGTATGAAGTCTTTGGCGTCCCA
ATGCCTAGATGCGGTCTGGATCCAAGACGAGCGCGTTGCGGAAATTCGGGGTTGGAAGAATGACGCCTAACCGGTCGTTCGAGCGGACTGCC
CTCGGCAAGCCTCGGTCAGCCGCTCAACTTCAACG

*gcuF*

TTAGCACCACTGAAACCCAGCTTTATTTAGCTCATGTTTATTCAAACGGCATTTAGCTTTTCAGGCGTTATTCAGTGCCTGTTTTGCCTTTTTTC
CGGGCTTCGCCTGCATGGGCTGCGCAGGTTTTCAGTCTTTTTGGCCTCTAGCCCTTGCGTAGCAAGCGCAAGCAGCTATCGTTTTTGCAGTGCT
GTGCCGCCTCGGTGGCGCAGCGTTTTTTCACGGTTAGCGCCCGTCGCCAAATTCAAGTTATCCGTTTTGGCTTCTGGTTCTAACATTTCGGTCA
AGCCGACCCGCATTCTGCGGTCGGCTTACCTCGCCCG

*gcuI*

TTAGGTGCACACATGCCAAGAATTTTTACTCAACTCATAGCGTTACTTCTATTGGTTGCGTGCAAACCTGAATCTGTACCGCTTAATGAATCAT
TTACCTGTTCTGAAAACCCCATCGAATTAGAACAGATCTCACAACTATTGAAAAATGAGCATGTGCCATTCACTCAAGAAACAGTTTTGGTCG
ACAGGGTATCATTTACTTCAGCGGGCGTAAAAAGTGTTCTCCCTAAGGGGGCTACGTGCATTTTCTACTCACATAAATACAGCAATATTACGA
ATAAAATTAACAAAGAAGTCATTGATGACTTTCCGCCAATCTGGGGTAGAGCTACCTCTTGGGACGATCGAAACGCCCAATTGGTTGAAAAAT
TATCGCAACACGGCATCCAGACAACAAGGCGCGTTTATAGAGGTAAAGAATGGATCGCATGGCCCATAGAAGATGTCGAGTTGGCCGAAAAT
ATTTTAGGCTTCAATGAGAAAATGAAACAGTATTTTAAAGAGCAGCGGCTTAAAATGTATGCACCTAACCAGGCGCTCAAGGCGCAGCCTTCG
GCTACCGTGGACGCCGCAAGCAGCGCCTCTTAGCTCTACG

*gcuJ*

TTAACTTGAGGGGAATTAAACATCGTGCATGATTTTAACGGTGCACAAATCTTTGTTGGGGCTAAAGTTAAAGTTCTATTTATTGATCAAACGA
TTATGAATGGACTGCCATCAAGCGAAGTAAATGACATTAATTCTATGCTGGGTGAAATATTCGAAGTTTACGAAATAGACGAATATGGCCAAG
CATGGGTTGAAAAGTGGTGGCATTTAGATGAAGGTAAATCGTATTCGCACTCTTTAGGCTTGTCCTCAACAGAGCTTGAGTTAGTAGAAAAAT
AAACCAACAATAGTGCATTTAAGTCAAAAGTTAACAAGGCGCTCCAGCAATGCGCACTTTGTGCGCTCGACTTATAAACTGCGGCTGAGCTTA
GCG

*gcuN*

TTAGGAATTCAATGGACGTAGACCTGCAAAGCATTGGAATACAAACCGTCCAGGAAGCATTGGAGAAAGGTTCACTCATTTTTAATGAAAAA
ACCAAAACCGATGACTTCAACAGGGCTTGTGATCACATACTTGTGCTGCTAGTAGATGCATTTCAGTGCTTTGATCGTGGCTCTTGGGGTACAT
CGGTATTTCTTTCAATTACTGCAATTGAAGAAGTAGCAAAAGCAGAGGTTGGCCTTTATCGAAGAGAAGGCAAAATCGGAAAAGCTAAGCGC
GGCAAAGATAAGCTTTTCAACCATCAAGAAAAACACCGCATGGCTATATTGCCTACAGTATTTATGAGTAAGCGCTTAGAAGAGGCTTTAGGC
AAAGAAAAATGCGCCGAACTACTAAAAGAGGCTGCACATGGGGAGTTTAGAAATCTCAGAGAGTCATCCTTGTATTTTTCAAATGAAAACGG
CCAATTTGTCACTCCTGCAAACGTTGTATCCCAAAACAGGGCAAAAGAATTTTTGCTTTTAGCGTTGGAAGCCGCAGATGATCGCCTCGTAGG
TTACACGAACCATACCGGAATCTTAGAGGCTAAAATTAACGAGATCTTCAGCGTTGTGGCCCATTCCTAACAAATGGTTCAAATCGCTCGCTC
CGCTCGCTGGGACGGGCTAAAGCCGCCCCTTAACCAAACG

*gcuP*

TTGGGCAACAAGAAAACCGATATGAACGTACGCACTTGCACTGAATCTGACGTCGCCTCTATCGCAGTCGTATTTACTGAGTCTATTCATGTA
CTTGGAGCGTCTCACTATGACGCTTCGCAAAGGAATGCGTGGGCACCGCGTCCCGCAGATATAGAGGCTTGGTCAGCTCGCTTATCTGGCCTA
CAGACTCTTCTAGCAATTGAGGGAGATGCGGTTATCGGGTTCATCTCTTACGAGCTTAGCGGCCACATCGAGTTTCTTTACACCGCACCGGGTT
CCGAGCGTCGGGGCGTCGCGTCTGTTCTGTACCGTGAGGTTGAGAAAGCCCTCCCAGGTGTTTCGCTCTTCACAGAAGCCAGTCTGGTCGCCA
AGCCCTTTTTCCTGCGGCATGGTTTCAGTGTAGTTGAGGAGCAAAATGTCTCCCGTGGAGGCGTCATGTTCCGTAGGTATGCAATGCGCAAGG
CAGTTGTCGCCCAACATGGCGCTCAAGCCGACCGGCCAGCCCTGCGGGCTGTCCGTCGGCTTAGCTAGGGCG

*gcuQ*

TTAGAGCAAAAATGAACTGCGCACGCTGTGGCCGCCAGAATTTCGTAGATGCACATCGTTGCGCTCACTGCGGCCACCACTTCTTTGACGGTA
GTGACGTGGTGAATCTTCTAGACAAGGCTCAGAAGCGCACTTGGTTAAGGCCGCGAAACATCGCCATATTCCTTGCGGCCTGGGCGCTACTGG
TGGTGGAGATCGTCACCTACGCGGGTAAAGCGCCAAGTTCTCCTCAGGGCTGGGCGGCGCTCATACTGCTCGGGCCTATTGTGTACGTCGTAC
TAGCAACCTGTGCAGATACAATTTTTCGAAAGCCATGAATACAGTAAATCGCTCATGCTCTAACCAATGCTTCCAGCCGACGCTACGGCCTTA
CGGCCTCCGCGCGTCTGAAGCTGGGCG

*gcuWGS1*

TTCGGCGCACGAGGGGCCACGGTCAAGGTTGACGCTACCGATGAACACGCAGCCGTCGTGCGGCTGTTCAGCCATGTGGTGAATGGGCAGAG
AGCAAGTTGTGTAGGATTGACCGTTGCCGTACAGTCGCCGCGATGCTCAGCCAGTTTAGGCCCGTGTCGGCACGACTGGACGTTGTAAAGTTT
TTTTATCGACAAGGAGGAAGAAGCATGAAATTTCAGCGATTGACCTCGGTACTAGTGTTCGTACTCGCCGGCAGCCTGTTTCTGACTGCATCC
GTTGCCTCGGCCGGGCCGAAGGAGGACGTTGCCGCCGCGACCATGAAATGGGCCCAGACTTTTGGACAAAACGATCCTGACAAAGTTGTTCC
TCTCTATGCCACTGATGGAGTTCTCTGGGGAACTATCTCTCCTACCGTGCGAGCGGATCGAGCGGCGTTGCGCGACTACTTCGTCACGGCATTC
AAGGTCTTGCCCAAGGTCACATTCGGCGAGCAATTGATCCGCGTTTATGGCGGCACCGCTGTCAACACCGGCTACTACACGTTCTCGTATAAC
AAGGACGGCGAAGCCAAGACGCTGCCAGCGCGTTACAGCTTCACATTCGTGAAAGAGGGCGAGAACTGGATAATCGTGGACCATCACTCATC
GGCGATGCCCGCCCCTCTTCGGTAGGCACACGCCTAGGGCCGGAGGGCCATGCTCGAACAGGATCGCGGATGACCTGGGATATGGCCGACAA
GCGAACTTTTCGAGCCCCGCCGAACTTTAAATGCAGCGGACCGCCTTCGGCGGCCGCTGATTATCGACG

*gcuWGS2*

TTAGGGGCACATGAGAGTTCTTGCAGCGTCCTTGGCTTTCTTATTGCTAACTGGTTGCGCACATTGGTTCGGCCCTGCTGACGGCGCCTTTTAT
GCAGTTGGCTCTGCGCCGGAGAGTCATTCTTGCAAGCTATCCGTGGCTCCTGTTGGCTCTAGCAACCCTCCAACGGAGTGGAGCGTTGTAGGC
AACTTCCGACAGAGTGTTGTCGTCAATCATTCGCGCAAGGGGCATAGGGTGGCTCTCATTTGCAACGGTGAGCTGGCTGCCGAGCGAACCTTC
AAGTACGGTCGTGATGTAGGTATTGGCGGGGAACTTGCGGTCAATGGCAGTGCCCCCTAACAATTCATTCAAGCCGAAGCCGCTTCGCGGCTC
GGCTTAATTCAGGCG

*gcuWGS10*

TTAGGCATACCAGATAGCACGCACCTACTAACCAACAAGCCATCATCTAAAAATGCAGGAAACAACGTACCCTAATCGGATTGACATTGGAA
CGCTCCAAGTTGCGGACACATGGAAGAAAAAATTCCGCGCTCTTGAGCAGGCGGGCGGCCCAAAGCTCCCACACCTCAAGACGCTGCCATTC
AAGGAGCGATTCAGCGTTACCTTCAACTTTCTTGGATTCATATTCGGACCGTTGTATTATTTGGCCAAGGGCATGTGGAAGAAGGCGCTATTTC
TGTTTGCCGCCTGCGTTGCAGCCGTTCTACTTATCGGCTCAGCCTTAGACATGGCCGGGTTACCCGAGGTGGCCAACTCACTAGGGTATGGGGT
CTCAGCATTCTTTGGGGTTCGGTCAAATATCGACTACTACAAAAAAATGGTGTTGGGTGAGAATGGCTGGTGGTAACATTAAAAATATCCGAA
ACTTTGCCCATGCCTAACTGGGCGTTCAACGTGGACGCCAACAGCGGCCATGCCTTCGGCATTTTATTGGCCGCTGTTGGTGCCCTGCGCCCCT
GCGGGGCTCCGGCGCCAGTTAACTAATTCG

*gcuWGS21*

TTAAATTAGGGAAGCTTATGAGCAACGACCCAAGAACAGAACCTCTAGTGGAGTGGAATCGGCTGGCTCGGGAAAATACCGAAAATGCAATT
GTTTCATCTATGTTTGAAGCATCTGCTAAAGCAAGCGAACCCCTGGAGAAATTTTCCACATGGCTGCTTGTTGGCGCTGCAGCAATCGCATCTT
TTTTAATCGTAAACTCAGATCAAATAATTAATTTGGTCCGCAAAGTTGGCTATTTATCTTGCGGTGCGTTCTTATGCCTTTCGTGCATTTTTGGC
TTAATTTCAAAAATATACGCATTACGTTGTAAAATTGGTATTGAAGTTAGCTCTGCCGTAAGAAATACCTTCTCTATACATTTGGCAAACTACG
AAAAGGAAGAAGAAAAAATTCAAGAAAGTGCAAGTTTCTGGGGAATTACACTTGAAACCGGTGTTCGTATTGAAAGAGTGCTTAATGAATTT
TACAAGCCACTACCAAAAATAATCGCATGGTATTCACGCCGTATATTGGAAAAACACAAAGGCAATCCTCAAGTAGGTCATATATTGCTTATT
TCCATGCTAAATAAACAAGGCTTATTTACAATGCTTCAAGCCTTGTCGTTTCTAGGTTTTTTAATTGCAGGCTTTGCTTTTGCAGCAAAAATTTA
ACAAGGCACTGCACCTGACGAATTGTAAGTAGCGCTTATGCAATATCGCTACGCTAAATTTATCGCAACGCGCTACTTACAATTCGCAGGTGA
GCGCAACG
